# Benchmarking imputation methods for discrete biological data

**DOI:** 10.1101/2023.04.06.535892

**Authors:** Matthieu Gendre, Torsten Hauffe, Catalina Pimiento, Daniele Silvestro

## Abstract

Trait datasets are at the basis of a large share of ecology and evolutionary research, being used to infer ancestral morphologies, to quantify species extinction risks, or to evaluate the functional diversity of biological communities. These datasets, however, are often plagued by missing data, for instance due to incomplete sampling limited data and resource availabilities. Several imputation methods exist to predict missing values and have been successfully evaluated and used to fill the gaps in datasets of quantitative traits. Here we explore the performance of different imputation methods on discrete biological traits i.e. qualitative or categorical traits such as diet or habitat. We develop a bioinformatics pipeline to impute trait data combining phylogenetic, machine learning, and deep learning methods while integrating a simulation framework to evaluate their performance on synthetic datasets. Using this pipeline we run a wide range of simulations under different missing rates, mechanisms, and biases and different evolutionary models. Our results indicate that a new ensemble approach, where we combined the imputation results of a selection of imputation methods provides the most robust and accurate prediction of missing discrete traits. We apply our pipeline to an incomplete trait dataset of 1015 elasmobranch species (including sharks and rays) and found a high imputation accuracy of the predictions based on an expert-based assessment of the missing traits. Our bioinformatic pipeline, implemented in an open-source R package, facilitates the application and comparison of multiple imputation methods to make robust predictions of missing trait values in biological datasets.

## Introduction

A substantial part of ecology and evolution research relies on biological traits that can be measured or described across individual organisms or species. For instance, trait datasets can be used to infer ancestral morphologies and to model phenotypic evolution in a phylogenetic context, for instance the emergence of gigantism in elasmobranchs (Pimiento et al., 2019) or the evolution of climatic niche in walnut trees (Zhang et al., 2022). Traits are also used to evaluate the functional diversity of modern and past biological communities (Jeliazkov et al., 2020; Pimiento et al., 2020a), and to quantify species extinction risks and conservation priorities (Pimiento et al., 2020b). Biological datasets can include life history traits (e.g. body size, reproductive strategy, ecological traits Cooper and Purvis, 2010), ecological traits (e.g. size of the home range, diet; Galán-Acedo et al., 2019), and spatial and climatic features (e.g. geographic range sizes, temperature tolerances; Johnson et al., 2021; Schneider et al., 2019). Traits can be classified in two categories: continuous variables (e.g., body mass) and discrete features (e.g., presence/absence of an anatomical feature, diet, or habitat). Discrete features can be nominal (e.g., herbivorous or carnivorous diet), ordinal (e.g., shade tolerance gradient in plants), or quantitative which are variables composed of integers with a finite number of states (e.g. number of meals per day; Kaliyadan and Kulkarni, 2019).

Unfortunately, trait datasets are often incomplete, and more so when resulting from large compilations spanning many species. trait data may be missing for several reasons: for instance, a species is rare or occurs in habitats with limited accessibility and cannot be sampled for measurements (Nakagawa and Freckleton, 2008), or species or taxonomic group is less “charismatic” than others and less well represented in the available literature. Additionally, biological samples (e.g. from herbarium or museum specimens) might be incompletely preserved, or the trait is difficult to quantify (Penone et al., 2014). Incomplete trait data is almost inevitable when dealing with extinct species, for which the fossil record can inherently only provide partial information (Ronquist et al., 2012). This is problematic because incomplete data might reduce the statistical power of our inferences, and even prevent some analyses that cannot handle missing data, such as calculating functional diversity indices (Mason et al., 2005; Villéger et al., 2008), and it may result in spurious results due to non-random extinctions, such as in the case of flightless-ness evolution in birds (Sayol et al., 2020).

In statistical research, the distribution of missing values can be characterized by three types of missing mechanisms (Johnson et al., 2021, see Supplementary Information Figures S7–9): (1) Missing completely at random (MCAR) where a random sample of data is missing, (2) Missing at random (MAR) where the missing values are related to other observed traits in the dataset, but independent of the unobserved data themselves, and (3) Missing not at random (MNAR) where missing data relates only to unobserved data, which means that missing values depend on factors not included in the dataset (Mack et al., 2018; Papageorgiou et al., 2018). For instance, if in a body mass dataset, species with small body mass are disproportionally missing, the distribution of missing values is MNAR. However, if the dataset also includes body length for all species and length is correlated with mass, then the distribution is MAR. In this study, we introduce a fourth type of missing mechanism describing the case in which species belonging to particular clades are more likely to be missing trait data. This captures the case in which the taxon sampling in a trait dataset is uneven. Hereafter, we refer to this distribution of missing data as phyloNA.

An MCAR pattern will typically lead to an unbiased interpretation of the data because missing values are randomly distributed, but it will reduce the sample size and the statistical power of data analysis. In contrast, instances of MAR, MNAR, and phyloNA induce a bias. The bias induced is potentially known when values are MAR or phyloNA, while for the MNAR mechanisms the bias is unknown because it cannot be explained by the observed data.

Several strategies have been developed to deal with datasets containing missing data. The simplest strategy to overcome the problem, is the “complete-case analysis”, which consists of removing traits or species with missing data prior to the analysis (Johnson et al., 2021). This method, however, presents two main drawbacks. First it decreases the sample size thus reducing the statistical power of follow-up analyses (Nakagawa and Freckleton, 2008). Second, it can induce a bias in the estimated parameters of the statistical analysis (Papageorgiou et al., 2018; Debastiani et al., 2021; Johnson et al., 2021), if the distribution of missing data is not MCAR.

Another way to deal with missing values, is to use imputation methods to fill the gaps. Data imputation consists of replacing a missing value with a value predicted by a statistical method. There exist a variety of methods for data imputation ranging from a simple principle where missing values are replaced by the mean of the trait to sophisticated machine learning approaches.

In biological datasets, missing trait data are often imputed using phylogenetic information, where the missing traits are predicted while considering the shared evolutionary history among species. However, not all traits are conserved in a phylogeny, and different degrees of phylogenetic signals have been observed in life-history traits and ecological traits (Kamilar and Cooper, 2013). Phylogenetic information can be integrated in the imputation through the use of evolutionary models as implemented in phylogenetic comparative methods (Harmon and Open Textbook Library, 2019), or in the form of additional predictive variables (e.g. from an eigenvector decopomosition of the phylogeny, Diniz-Filho et al. (1998)) in the dataset using machine learning methods (Debastiani et al., 2021).

Several studies have assessed the performance of different imputation methods applied to continuous traits (Penone et al., 2014; Johnson et al., 2021), showing that both phylogenetic comparative methods and machine learning approaches can provide reliable imputation of quantitative traits in many, but not all cases, and depending on the level of phylogenetic signal in the trait data. Less is known, however, about the performance of imputation methods with discrete traits, even though many biological traits are discrete in nature, for instance, host-specificity in fungi (Põlme et al., 2020), morphological structures like flower color (Cappellari et al., 2022), feeding type (Froese Rainer and Pauly Daniel, 2022) or the substrate on which the species lives (Meiri, 2018).

Here, we evaluate the performance of a range of imputation methods on discrete nominal trait data and used simulations to develop guidelines for handling incomplete biological datasets. We tested five imputation approaches: a phylogenetic comparative method based on a Markov model of trait evolution (as implemented in corHMM; Beaulieu et al., 2013), the k-nearest neighbour approach (kNN; Fix and Hodges, 1951), a non parametric method based on random forests (missForest; Stekhoven and Buhlmann, 2012), multinomial logistic regression (MICE; van Buuren, 2007), and a deep learning approach based on generative adversarial networks (GAIN; Yoon et al., 2018). In addition, we evaluated an ensemble method based on hard voting (Bruzzone et al., 2002), aggregating the output of the three machine learning methods coupled with phylogenetic imputation. To facilitate the comparison and benchmarking of the different imputation methods, we implemented a bioinformatic pipeline to perform simulations and imputations under different strategies. Our framework can pre-process the data and used them to identify the approaches predicted to yield the highest accuracy and reliability under different scenarios. We evaluate our imputation pipeline with an empirical data from the elasmobranch clade, which includes sharks, rays, skates, and sawfish, benchmarking it against an expert-based assessment of the missing trait as an empirical measure of accuracy.

## Materials and Methods

We developed a framework to assess the performance of different imputation methods in filling gaps in biological trait data based on simulations (Fig. 1). Our pipeline involves four steps: (1) simulating a phylogenetic tree and trait data under different evolutionary scenarios, (2) generating gaps in the data according to different missing mechanisms and rates, (3) imputing missing values using a range of imputation methods and strategies, and (4) comparing the accuracy of the imputations across methods.

**Figure 1:**
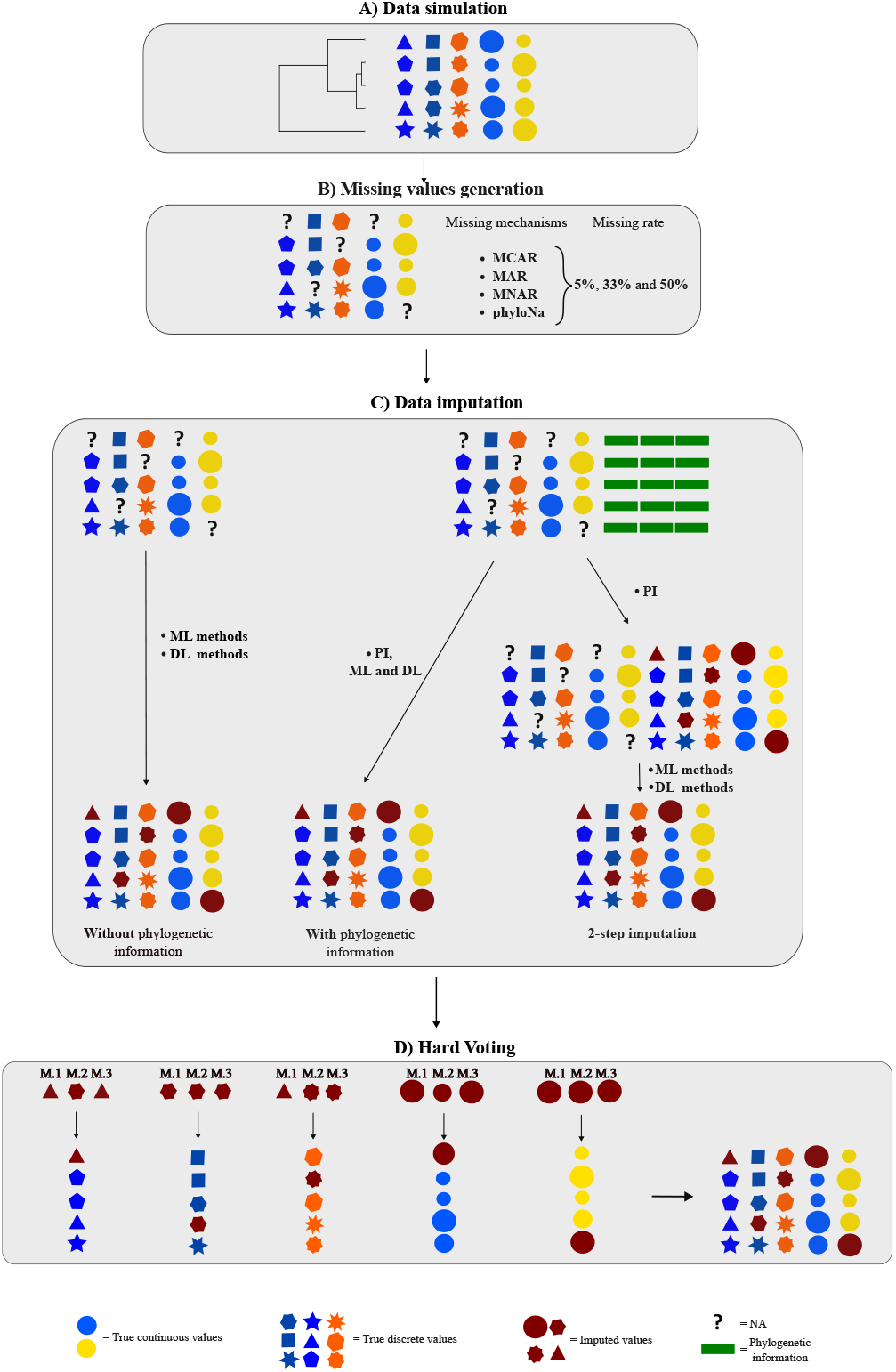
A pipeline to compare the performance of different imputation methods to impute discrete traits. **A)** Simulation of a phylogenetic tree and data with discrete and continuous traits. **B)** Generation of missing values according to different mechanisms (MCAR, MAR, MNAR, phyloNA). **C)** Imputation of missing data through three different strategies. **D)** A Hard Voting classifier combining the output of different methods {M1, M2, …}.

For the imputation step we included phylogenetic imputation (PI), non-parametric or semi-parametric machine learning methods (ML), and a parametric deep learning method (DL). We evaluated four strategies for prediction. The first strategy used ML and DL methods with the trait data without any phylogenetic information as input. The second strategy incorporated phylogenetic information to the data set (as detailed below), which previous studies on continuous traits have shown to improve the accuracy of the imputations (Fisher et al., 2003; Penone et al., 2014; Blomberg et al., 2020; Johnson et al., 2021). The third strategy used the output of the PI methods along with the trait data as input for ML and DL imputation methods. Finally, the fourth strategy were based on a hard voting ensemble that combines the output of different imputation methods to make a majority prediction (Bruzzone et al., 2002).

### Phylogenetic and trait data simulation

We generated simulated datasets reflecting what real biological data might look like, with phylogenetic information and trait data including continuous and discrete traits with some degree of covariance among traits.

For each simulation we generated a phylogenetic tree according to a stochastic birth-death process in which births (speciations) and deaths (extinctions) occur stochastically based on birth and death rates in a continuous-time Markov process. We simulated phylogenetic trees based on the number of extant species set to 100, with a birth rate of 0.4 and a death rate of 0.1. The extinct species were subsequently dropped from the tree. We then rescaled all trees to a height of 1 (Supplementary Information, Figure S7a). We then used the phylogeny to simulate datasets composed of 13 continuous and nominal traits. The first was a 3-state nominal trait and it was the one we focused on to benchmark different imputation methods, while the other traits were treated as auxiliary variables to perform the imputation. Thus, all the evaluations of imputation accuracy presented in the **??** refer to this trait. The auxiliary traits included three nominal and three continuous uncorrelated traits that evolved independently of the first trait and of each other, and three nominal and three continuous traits that were correlated to the first one.

In each dataset, we simulated uncorrelated nominal traits based on a Markov evolutionary model. For comparison, we also tested scenarios in which the nominal traits were generated under a threshold model (Felsenstein, 2005, see Supplementary Information). We drew the transition rates for the Markov model randomly from a uniform distribution 𝒰 (0, 0.5). For the threshold model, we simulated continuous traits according to a Brownian motion (BM) and discretized them based on threshold values randomly drawn from a uniform distribution spanning the trait range (Revell, 2014). We then assigned random labels to each interval to transform the continuous trait into a nominal one.

Uncorrelated continuous traits were generated as independent instances of a BM process. Under the BM model, the trait evolves according to a stochastic process in which the expected change in trait value after time *t* is normally distributed: 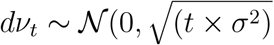 (Felsenstein, 1985). For each trait we drew with a rate parameter *σ*^2^ randomly drawn from a uniform distribution 𝒰 (1× 10^*−*4^, 0.5). We simulated correlated nominal traits so that one of the states is correlated with a state of the trait of interest. The pairs of correlated states were selected at random. The algorithm generates for each correlated trait a probability matrix of size *number of states S*_*i*_ × *S*_*j*_, with *S* set to number of states of traits *i, j* and a given correlation strength (here set equal to 0.8) between each state. Then, the traits are sampled with the corresponding weights present in the probability matrix. Continuous traits that are correlated with the trait of interest, were randomly drawn normally distributed vectors, with mean and standard deviation set equal to the trait of interest and a correlation strength of 0.8. All continuous traits were *z*-standardized prior to the analyses to have a a mean of 0 and a standard deviation of 1. The standardization is an important step in ML and DL to homogenize the weight of each trait in the analysis.

We explored how the strength of the phylogenetic signal impacts the performance of different imputation approaches. To achieve that, we altered the branch lengths of the tree based on the *λ* and *κ* models (Pagel, 1999b,a). The *λ* transformation affects the trait correlation between species due to shared evolutionary history, i.e. the phylogenetic signal in the trait (Cavender-Bares et al., 2020), such that with *λ* = 0, the phylogenetic tree results in a star phylogeny, which means that the internal branches have a length of 0 (see Supplementary Information, Figure S7b; Harmon and Open Textbook Library, 2019). As a result, the simulated traits evolve independently among the species. The second transformation is based on the *κ* model, that alters the branch lengths to change the evolutionary process from a gradual one (*κ* = 1, where trait change is proportional to evolutionary time) to a punctuated one (*κ* = 0, where trait change is proportional to the number of cladogenetic events). We repeated our trait simulations using rescaled trees with *λ* = 0.0001, to simulate traits with little phylogenetic signal, and *κ* = 0, to simulate traits under a punctuated evolution model.

In total, we simulated 900 datasets under the three different evolutionary scenarios, each performed 100 times with three different missing rates. In the Supplementary Information, we evaluated the same scenarios with the nominal traits simulated by the threshold evolutionary model.

### Missing values simulation

We simulated missing values (here after NAs) under different patterns following the MCAR, MAR, MNAR distributions described in the literature (Johnson et al. (2021)) and a phyloNA distribution. NAs occurred in all 13 traits, even though we focused our assessment of the imputation only on the first trait. The only exception is for values that were MAR, in this case missing values were simulated only in the first trait. The missing values were simulated according to three fractions, 5%, 33% and 50% (Supplementary Information, Figures S8–10) using missMethods 0.4 (Rockel, 2022).

In nominal traits, the missing values were generated such that each state remained present in the trait after NA assignment except for phyloNA missing values. We simulated the phyloNA values by randomly selecting a clade that is larger or equal to the missing rate and set all trait values to NA for a number of randomly chosen clade members to satisfy the specified fraction of missing data.

### Imputation of missing values

We tested five methods to impute NAs that can be grouped into phylogenetic imputation (PI) methods, machine learning approaches, and deep learning methods. The machine learning and deep learning methods can impute missing values in mixed dataset (continuous and discrete variables), while PI uses different models depending on the nature of the trait, i.e., Markov or BM models for discrete and continuous traits, respectively. PI methods were based on the libraries Rphylopars 1.1.0.9004 (Goolsby et al., 2017) and corHMM 2.8 (Beaulieu et al., 2013) for the R 4.2; statistical programming environment (R Core Team, 2013), while machine learning methods were based on the missForest 1.4 (Stekhoven and Buhlmann, 2012), kNN 6.1.1 (Kowarik and Templ, 2016) and MICE 3.14.0 (Buuren and Groothuis-Oudshoorn, 2011) packages. We also used a deep learning method based on generative adversarial networks and implemented in the GAIN Python library (Yoon et al., 2018). Additional details about the imputation methods and settings are provided in the Supplementary Information.

### Imputation strategies

We tested and compared different imputation strategies (Fig. 1). First, we imputed the traits using ML and DL methods only based on trait data, without phylogenetic information. Second, we included the tree in the imputation and ran PI as well as ML and DL, for which the phylogenetic information was provided through phylogenetic eigenvectors (Diniz-Filho et al. (1998); Penone et al. (2014)). We performed the eigenvector decomposition using the PVR 0.3 package (Santos, 2018) and included a number of eigenvectors capturing 95% of the tree variance. Third, we combined PI with ML in a two-step approach where the dataset was augmented with the output of a first phylogenetic imputation prior to ML and DL imputation. Since corHMM estimates relative probabilities associated to each state for the imputed species, we included the full probability vector as input for the second imputation. Finally, we used a hard voting ensemble method (hereafter: HV), which aggregates the output of multiple models and returns the most voted state as the final output. Based on the performance of the individual models, we chose to aggregate three models: missForest without phylogenetic data and MICE and kNN based on the 2-steps strategy.

### Error calculation

We benchmarked the different imputation methods and strategies across all simulations in terms of accuracy and imputation error, which we defined as the proportion of correct (or erroneous) imputations out of the total number of imputed values. We calculated the accuracy and error for each simulated dataset and then summarized them as mean and standard deviation across replicates.

### Empirical application

We tested our pipeline on an empirical trait dataset of Elasmobranchii, which we obtained from Fishbase (Froese Rainer and Pauly Daniel, 2022) using the rfishbase 4.0.0 package (Boettiger et al., 2012), and combined with a comprehensive phylogeny (Stein et al., 2018). The trait dataset included 18 traits for 1015 extant species, which are all present in the phylogeny. Eight traits were discrete: The traits “Fresh water”, “Brackish”, and “Salt water” were binary traits, the trait “Water column” described the position in the water column (demersal or pelagic), “Migration” was a trait of mass relocation (amphidromous, oceanodromous, or potamodromous), “Dangerous” provided information about the species’ threat to humans (harmless, other, poisonous to eat, traumatogenic, or venomous), “Feeding type” indicated whether the species was mainly herbivorous or carnivorous (i.e. mainly animals and a trophic level of 2.8 or higher) or plants/detritus+animals (i.e trophic level 2.2–2.79), and “Feeding habit” described the feeding habit of each species (browsing on substrate, filtering plankton, hunting macrofauna (predator), selective plankton feeding, variable). The discrete traits had 2 to 7 states and the amount of missing values ranged from 0 to 91% depending on the trait (Supplementary Information, Table S28). In addition, we discretized the traits “Maximum depth” and “Length”, following previous work (Paillard et al. (2021)). The trait “Maximum depth” was converted to a binary trait: *x <* 200 m (epipelagic) and *x >*= 200 m (mesopelagic) and the trait “Length” to three states: small (*x <* 150 cm), medium (150 cm =*< x <* 300 cm), and large (*x >*= 300 cm).

The accuracy of imputed traits cannot be calculated for empirical data sets because the true values are unknown. We therefore evaluated the accuracy in two ways: first we compared our predictions with an expert-based approach, where we one of the authors (CP) filled the gaps based prior knowledge and before seeing the predictions generated by the imputation pipeline. Second, we used a random sampling approach where we added an additional 1% of missing values completely at random on top of the already missing data in each state. Using this mechanism, we introduced an amount of missing values proportional of the empirical frequency of the states, thus preserving the original pattern of missing data (Supplementary Information, Table S28). We repeated the addition of missing values and trait imputation 10 times to calculate the mean and standard deviation for the expected imputation accuracy. Elasmobranch traits were imputed using MICE+PI, missForest, kNN+PI, and HV.

## Results

### Imputation accuracy with simulated data

To summarize our imputation results, we first combined all simulation settings with the same missing data rate (5, 33, 50%), making an overall ranking of the tested methods. The ranking was then dissected across different simulation scenarios. Overall, we found that the approach combining machine learning with a phylogenetic comparative method outperformed all others imputation strategies, irrespective of the level of phylogenetic signal, the model of evolution, the missing mechanisms, and the amount of correlation in the dataset (Fig. 2 and Supplementary Information Tables S2–11). For the simulated datasets composed of six traits correlated and six independent of the focal trait, the ensemble method showed for a missing rate of 33% an average accuracy of 68.9% (confidence interval: 1.3), which was 2.6% higher than missForest alone, 1.6% higher than missForest+PI, 0.6% greater than knn+PI and 3.9% higher than using phylogenetic imputation (Supplementary Information Table S3). The relative difference in performance between HV and the other methods remained consistent when varying the fraction of missing data to 5% or 50% (Supplementary Information, Tables S2–3).

**Figure 2:**
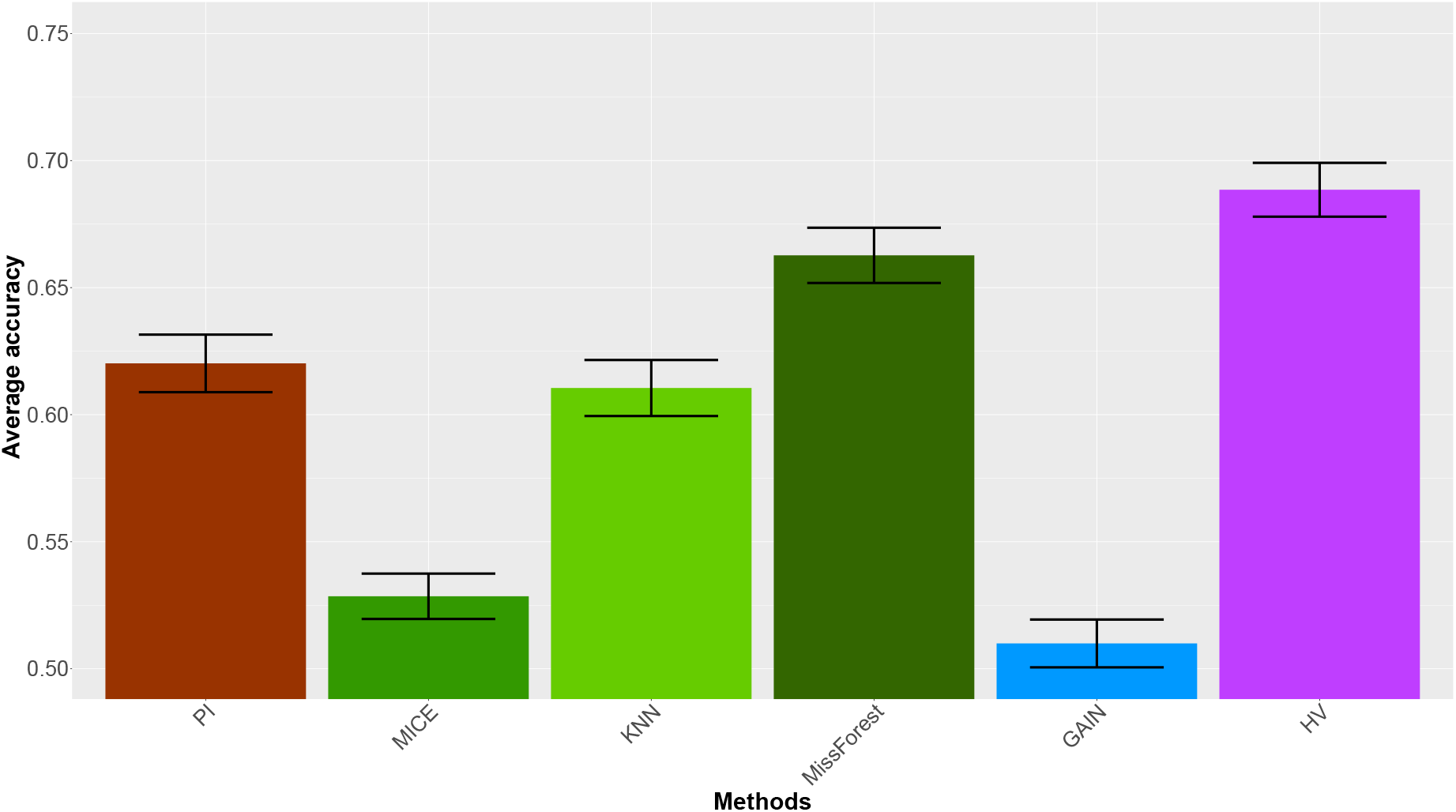
Barplot representing the average accuracy and the confidence interval of PI, MICE, kNN, missForest, GAIN, and HV. The Y axis represents the range of accuracy values aggregated for each replicate based on a 33% missing rate. The deep learning method is represented in blue, the machine learning methods in green and the phylogenetic imputation method in burgundy. We averaged the results of the datasets containing strongly correlated traits (*ρ* = 0.8) and independent traits (*rho* = 0.0). In both scenarios, the MK model was used to simulate discrete traits.

Using our 2-step imputation approach generally improved the accuracy and performed better than when phylogenetic eigenvectors were used as additional features. For instance, with a missing rate of 33%, the kNN+PI approach resulted in an averaged accuracy 1% higher than kNN and 2.3% higher compared to the kNN+P. The average accuracy of MICE+PI was 5.7% higher than MICE+P but only 0.1% greater than MICE, respectively. In contrast, and regardless the missing rate, GAIN did not benefit from the 2-steps strategy, but all variants of these approaches (with and without phylogenetic information or the 2-steps strategy) differed by 1.5% in accuracy for GAIN (Supplementary Information Table S3). However, when the missing rate was low with 5%, the 2-step strategy did not substantially improve the imputation.

When imputation missing values in datasets composed of strongly correlated traits, the ensemble approach improved the imputation accuracy over phylogenetic imputation method regardless of phylogenetic signal and across most missing mechanisms (Fig. 4). For instance, the ensemble method improved the accuracy by about 15% when the phylogenetic signal was strong and the missing values were MAR. (Fig. 4). In the near absence of phylogenetic signal (*λ* = 0.0001), HV returned the largest improvement in accuracy compared to PI when the missing data are MAR (27%, Fig. 4). The HV method outperformed phylogenetic imputation in almost all the scenarios, including those with missing rates of 5% or 50% (Supplementary Information, Tables S15 and S27). When it is not the best method, for instance, in the case of phylogenetically clustered missing data (phyloNA), it achieved a performance comparable to that of the best approach (PI; Fig. 4).

Based on the aggregated results (Fig. 2), missForest came second to HV in accuracy among all scenarios but its performance was more variable across the scenarios. With a missing rate of 33% and missing data distributed according to MCAR or MAR, the missForest results were comparable to HV (Supplementary Information, Table S17 and S19) in presence of a strong phylogenetic signal. In contrast, when the missing values were MNAR, missForest returned slightly less accurate imputations (Supplementary Information, Table S21) and when the missing values were phyloNA with a strong phylogenetic signal the accuracy of miss-Forest decreased by 12.5% compared to HV (Supplementary Information, Table S23. We observed similar patterns across different missing rates (Supplementary Information, Tables S2–4, S12–15 and S24–S27).

When the focal trait was independent of all the other traits (i.e *rho* = 0.0) the most accurate methods were PI and missForest+PI followed by HV, regardless the missing mechanism and the phylogenetic signal, (Supplementary Information, Tables S17b, S19b, S21b, and S23b). For instance, in a scenario where the missing data was MAR and the phylogenetic signal was strong, PI returned an accuracy of 81.5% (sd: 16.3), which was 4.3% greater than HV (Supplementary Information, Table S19b).

As expected, an increasing missing rates was associated with a decrease in accuracy of the imputations. For instance, a MNAR value was imputed with an accuracy of 86.4% (sd: 25.9) when the missing rate was 5% (Supplementary Information, Table S14), while at 50% the accuracy decreased to 62.1% (sd: 31.9) (Supplementary Information, Table S26). the accuracy decreased with an increasing missing rate regardless of the model of trait evolution and all missing mechanisms except phyloNA, where the imputation was robust to the missing rate (Fig. 3 and Supplementary Information, Figure S11).

**Figure 3:**
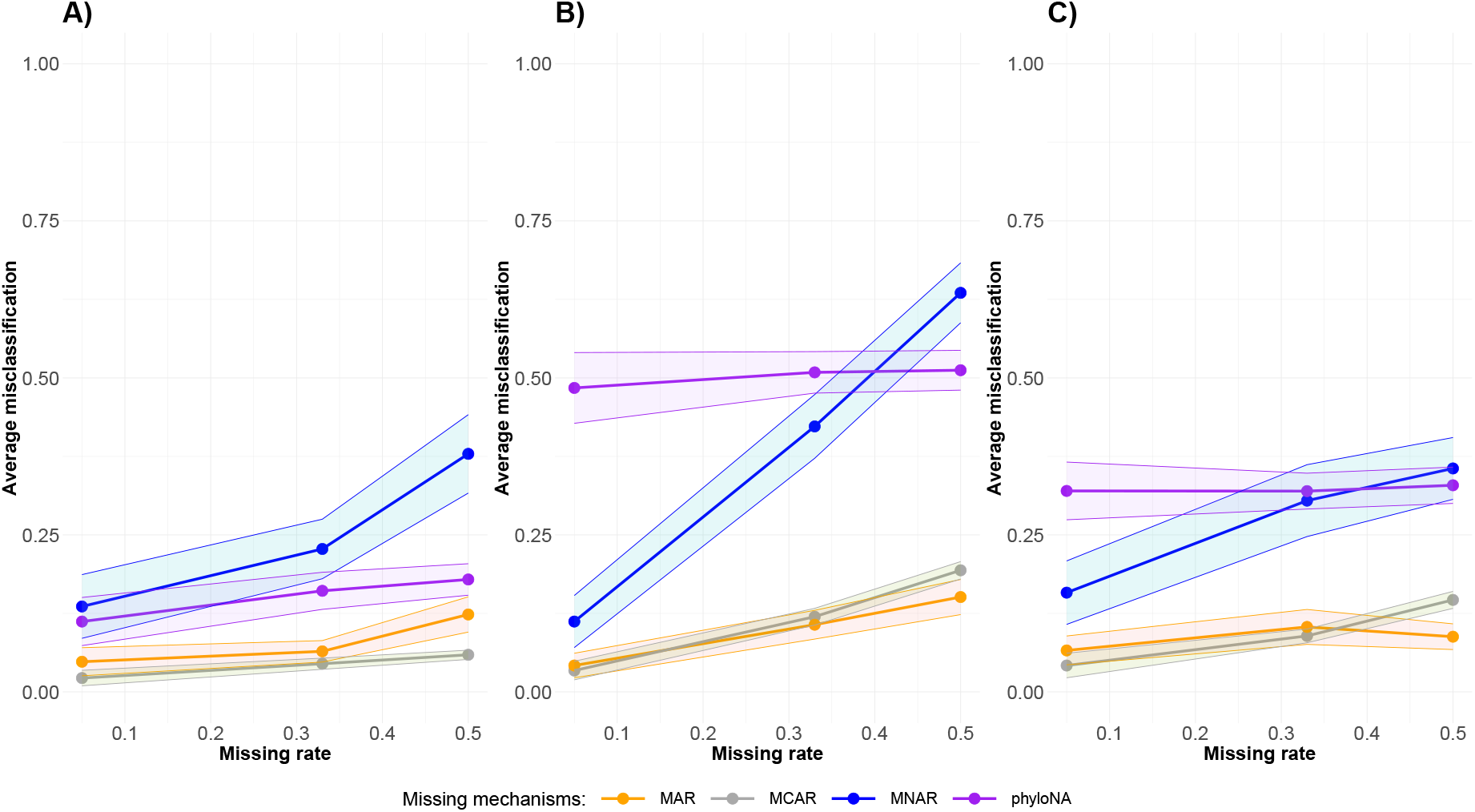
Effect of missing mechanisms and missing rates for different strengths in phylogenetic signal. **A)** Traits simulated with high phylogenetic signal of *κ* = 1 and *λ* = 1. **B)** Traits simulated with *κ* = 0 and *λ* = 1. **C)** Traits simulated with *κ* = 1 and *λ* = 0.0001. The dots represent the average misclassification of the hard voting approach and the shaded polygon represents the 95% confidence interval. The data were simulated through a Markov model and contained correlated traits (*ρ* = 0.8).

**Figure 4:**
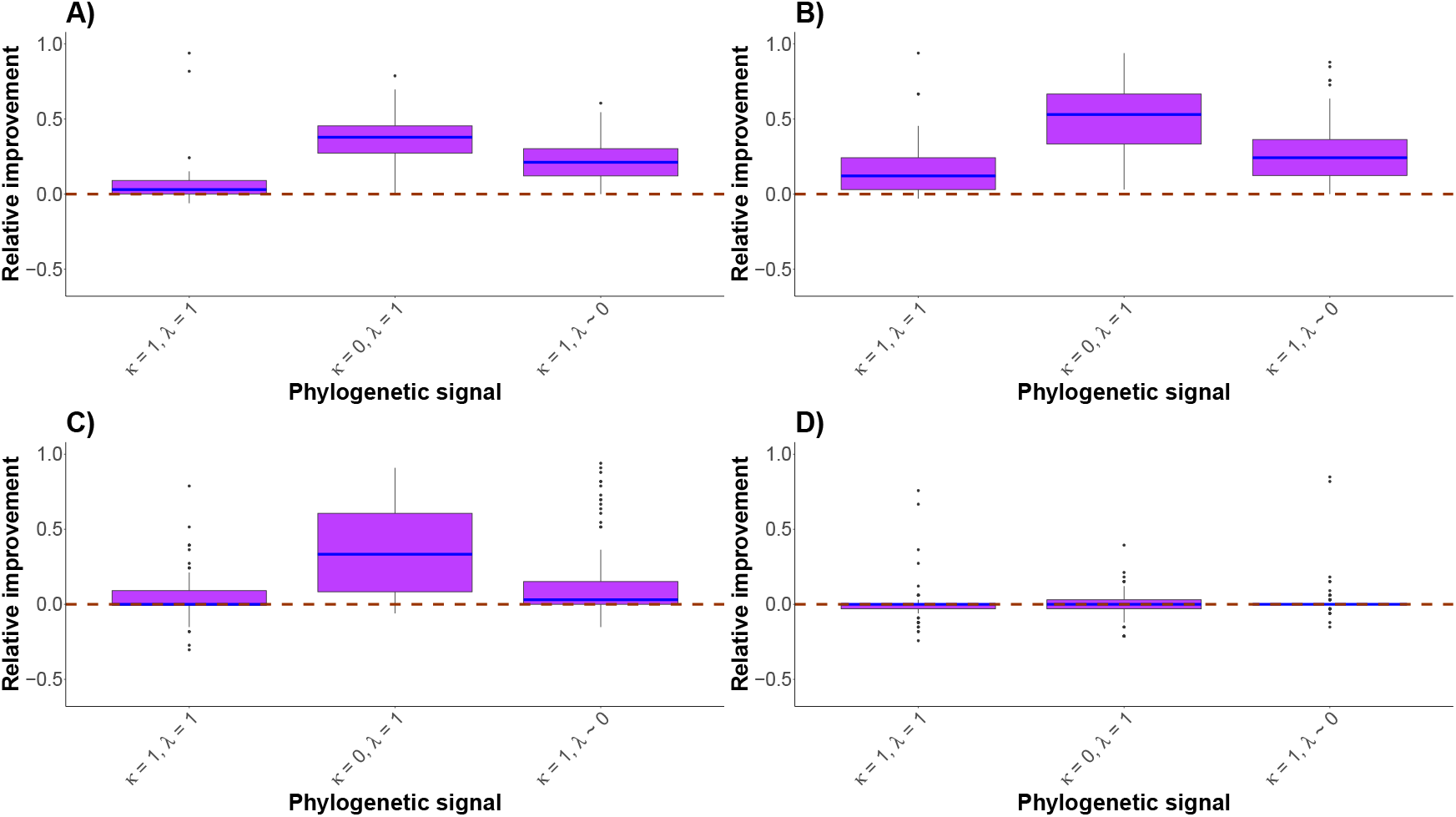
Improvement in performance of the hard voting (HV) method over phylogenetic imputation (PI) for different strengths in phylogenetic signals and four missing mechanisms (MCAR (**A**), MAR (**B**), MNAR (**C**) and phyloNA (**D**)). The dashed line indicates no difference in performance between the PI method and HV. The blue line represents the median. Data were simulated by an Markov model with a missing rate of 33% and contained strongly correlated traits (*ρ* = 0.8).

### Impact of biases in the missing data

Another factor that strongly influenced the imputation performance was the missing mechanism (Fig. 3). Missing values based on the MCAR or MAR mechanisms were imputed more accurately and with less variation across replicates than those based on MNAR or phyloNA. For instance, an MCAR value was imputed with 95.5% (sd: 4.5) accuracy (Supplementary Information, Table S17a)) with a strong phylogenetic signal, whereas the MNAR value was imputed with 77.2% (sd: 24.2) accuracy (Supplementary Information, Table S21a)).

### Impact of phylogenetic signal and trait correlations

The amount of phylogenetic signal in the imputed trait had a limited effect on the performance of HV imputation when the missing values followed the MAR or MCAR missing mechanisms. The accuracy varied by 7.5% between simulations with strong or a low phylogenetic signal (*κ* = 0 or *λ* = 0.0001) (Supplementary Information, Tables S17a and S19a). The effect was instead stronger when missing values were based on phyloNA or MNAR, in which case the accuracy of HV imputation decreased by 19.5–34.8% between a strong and a low phylogenetic signal (Supplementary Information, Tables S21a and S23a). We observed a similar effect at missing rates of 50% (Supplementary Information, Tables S26–27), while at 5% missing rate the imputations were accurate even in presence of MNAR values (Supplementary Information, Table S14).

The phylogenetic signal had a greater impact on the accuracy when the traits were weakly correlated (Supplementary Information, Tables S17b, S19b, S21b, and S23b). In the situation where the missing values were MAR, the accuracy varied by 32.8% between the simulations with a strong or a low phylogenetic signal (*κ* = 0) (Supplementary Information, Table S19b)).

The presence of a strong correlation among traits improved the imputation accuracy. MAR missing values were imputed with an accuracy of 93.5% (sd: 8.7) in the presence of a stron correlation among the traits and 77.3% (sd: 16.4) when the traits were independent (Supplementary Information, Table S19)). However, this difference was less pronounced for values missing according to a phyloNA mechanism (Supplementary Information, Table S23).

The model of trait evolution also had some impact on the accuracy of the HV imputations (Fig. 3, Supplementary Information, Figure S11). With 33% of missing data, a MNAR value was imputed 12.2% more accurately when it followed a Markov model compared with a threshold model (Supplementary Information, Table S20). The difference in performance due to the underlying evolutionary model was even stronger for simple phylogenetic imputation, showing that this method is more sensitive to model misspecification than ML. This difference of accuracy was not observed in all the situations, in case of 5% of missing data or when the missing mechanism were MAR, the values were imputed with the same accuracy (Supplementary Information, Tables S12–15, S18, and S25).

### Imputation of the Elasmobranchii dataset

The accuracy returned based on the expert-based approach ranged between 81% for the Maximum depth trait and 100% for the Feeding type trait (Fig. 5). In addition, all the accuracies from the expert-based approach exceeded the frequency of the most present state of each trait (Fig. 5), indicating that the accuracy did not only result from the imbalanced empirical state frequencies. In the random sampling approach, the imputation of the Maximum depth and Length traits produced similar accuracies to the expert-based approach (Fig. 5). However, for the Feeding type and Feeding habit traits we estimated a predicted accuracy of 83.3% and 57.3%, respectively, which were 17.3% and 42.3% lower than expert-based estimations (Supplementary Information, Table S29). This difference could be explained by the unbalanced distribution of the states in the traits (Supplementary Information, Figure S38).

**Figure 5:**
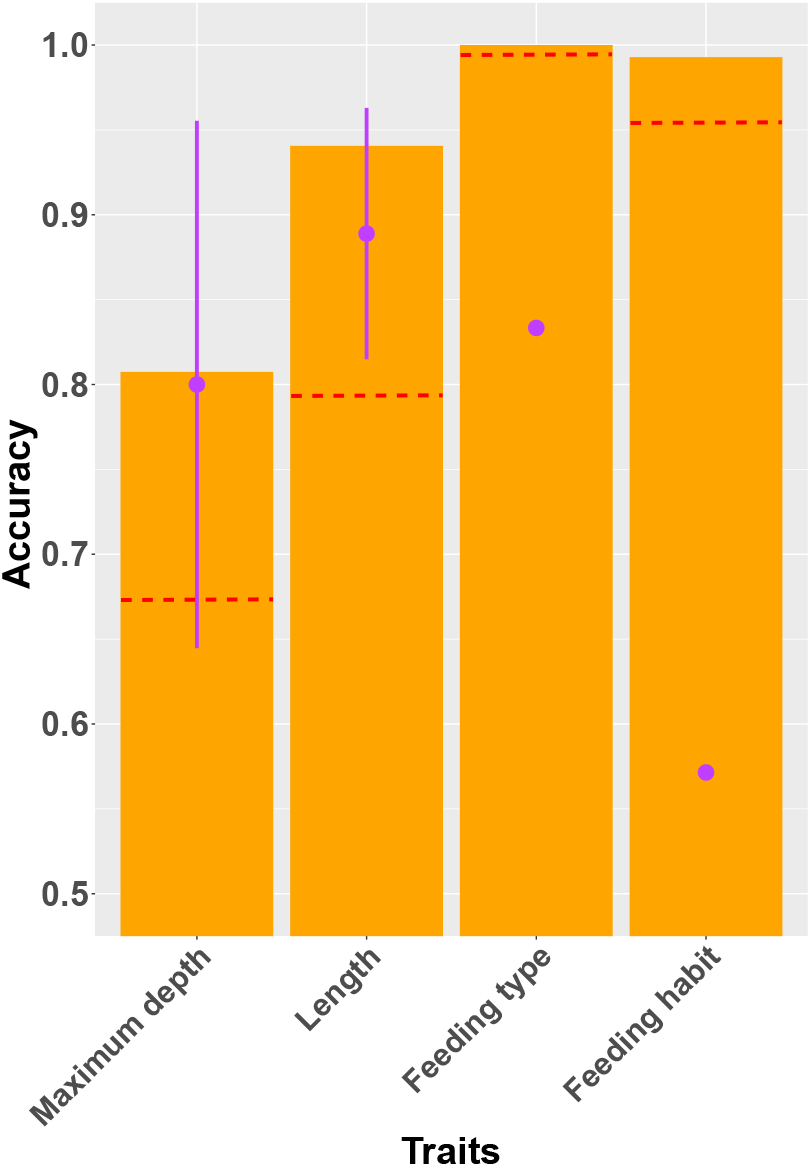
Imputation accuracy of four Elasmobranchii traits obtained by with hard voting. The orange bars represent the accuracy obtained by comparing the imputed data to the expert filled data. Purple dots display the mean accuracy comparing the true values with the imputed ones when an additional 1% of the non-missing data were omitted. Purple lines are the standard deviation across 10 replicated omissions. The red lines represent the frequency of the most present state.

## Discussion

The possibility to impute biological traits from incomplete datasets can help us filling knowledge gaps with realistic and accurate predictions, which can be used in downstream analyses of ecology and evolution. This is especially important when filling these gaps by direct observation is difficult, such as when the species is rare or extinct. For example, while soft tissues hold great importance in determining key features of animal ecology, they are rarely preserved in the fossil record and accurate imputation could help predicting these pieces of the puzzle in long-extinct taxa. For instance imputation can be used to predict phenotypic traits in dinosaurs like feathers or night vision that are only rarely preserved in the fossil record (Choiniere et al., 2021) of the conservation status of under-represented taxonomic groups in the Red List of the International Union for the Conservation of Nature (Silva et al., 2022).

With our extensive simulations we showed that discrete biological traits are most robustly imputed using a combination of phylogenetic and machine learning methods. The improvement in accuracy of the HV method when the missing values are MCAR, MAR, or MNAR compared to individual imputation methods likely derive from the ability of different methods to pick up slightly different signal from the data. While our HV approach was the most robust across the different simulation settings, changes in its performance do reflect previously observed patterns. For example, as expected, the accuracy of the model decreases as the missing rate increases (Debastiani et al., 2021; Johnson et al., 2021; Penone et al., 2014) because there is less information in the data.

Previous work showed that the missing mechanism has a strong impact on the accuracy of the imputation (Goberna and Verdú, 2016; Johnson et al., 2021) of missing values in continuous traits. Our results show that this is indeed the case for discrete traits too. While in an MCAR scenario the missing values are unbiased, thus leading to the highest prediction accuracy, deviations from that, such as the MNAR scenario, returned a generally worse performance, with the worst bias induced by the MNAR mechanism (Johnson et al., 2021). A potential solution to improve the accuracy of MNAR values would be to apply methods designed specifically for this type of missing mechanism (Molenberghs et al., 2014).

Phylogenetic signal quantifies the tendency for related species to resemble each other (Blomberg et al., 2003). Thus, when the phylogenetic signal is low, species relatedness has smaller predictive power, potentially leading to lower imputation accuracy (Debastiani et al., 2021; Swenson et al., 2017; Molina-Venegas et al., 2018; Goberna and Verdú, 2016; Kim et al., 2018). Our results also highlighted that the HV ensemble method was robust to changes in phylogenetic signal when the dataset were composed of strongly correlated traits (Supplementary Information, Tables S12–27). The accuracy is likely due to the correlations between the focal trait and some of the other traits in the simulated dataset as this has been shown to compensate for the lack of phylogenetic signal (Debastiani et al., 2021). Indeed, our simulations of uncorrelated traits showed a stronger impact of phylogenetic signal on the accuracy of the predictions.

We found the mode of evolution of the discrete trait to have a substantial impact on the accuracy of the imputation. For instance the accuracy returned for a trait simulated through a threshold model was of 12.2% greater when it followed a Markov model compared with a threshold model (Supplementary Information, Table S20). This variation in accuracy likely result from the adequacy of the implemented phylogenetic imputation methods, which assumed relatively simple Markov models in our experiments. This highlight the importance of using evolutionary models that reflect adequately the data, when performing phylogenetic imputation.

The analysis based on an empirical dataset of 1015 elasmobranch species confirmed that the ensemble method is a viable way to impute missing values (Fig. 5). The imputation method were more accurate than simpler approaches such as filling the missing values by the most frequent state of the trait or randomly. This finding was also true for traits that were highly imbalanced like Feeding type and Feeding habit (Supplementary Information, Figure S38).

Our random-sampling approach allowed us to evaluate the imputation performance providing a useful tool to assess the predicted accuracy even for empirical datasets and in the absence of an independent assessment of the missing data. This method, which added at least 1% of missing values in each state, demonstrated that HV was conservative and is not overestimating the confidence we can have in the imputed values (Fig. 5). This assessment is important to evaluate to what extent a trait is imputable and for instance showed that for the highly imbalanced Feeding habit trait the expected accuracy is low. This is not unexpected considering that the imputation methods tend to amplify the bias by filling in the gaps with data from the most common state (Somasundaram and Reddy, 2016). Therefore, this explains why the accuracy of the random-sampling approach was low. The accuracy might not be an optimal metric to evaluate the performance of a classifier when the data are strongly imbalanced, whereas a receiver operating characteristic curve (ROC), for example, could be a better metric (Chawla et al., 2002). However, these alternative metrics are more challenging to interpret for traits with more than two states. Thus, an independent assessment of the imputations based on alternative expert-based opinions, when applicable, remains a valuable way to evaluate the quality the imputation.

In conclusion, we showed that imputation of missing discrete values is a powerful and promising method for filling gaps in biological datasets and that ensemble methods aggregating different phylogenetic and machine learning methods are most likely to perform well across different datasets and settings. Our pipeline facilitates the use and comparisons across different imputation methods within a single package, building upon the extensive machine learning and phylogenetic libraries available in the R statistical environment. This will hopefully facilitate the exploration of the missing traits in empirical biological datasets and, within the limits of statistical imputation, help to accurately fill these knowledge gaps.

## Data availability statement

Code to perform simulations, imputations, and repeat all analyses is available at https://github.com/Matgend/Phylo_Imputationandresults at https://doi.org/10.5072/zenodo.1181410.

## Acknowledgements

TH and DS received funding from the Swiss National Science Foundation (PCEFP3_187012). DS additionally acknowledges funding from the Swedish Research Council (VR: 2019-04739), and the Swedish Foundation for Strategic Environmental Research MISTRA within the framework of the research programme BIOPATH (F 2022/1448). CP is funded by a PRIMA grant from the Swiss National Science Foundation (no. 185798).

