## Supporting Information for "Benchmarking imputation methods for discrete biological data"

### Benchmarking imputation methods for discrete biological data - Supporting Information

This is the "Supporting Information file of the paper called "Benchmarking imputation methods for discrete biological data". In this manuscript, you will find more details on the simulated dataset, on the distribution of the missing values, the imputation methods, and all the extra figures and tables generates through the analysis.

**5 Outputs of the empirical data** **61**

### 1 Simulation

#### 1.1 Trait data

We simulated datasets composed of 100 species and 13 traits. The traits can be continuous or discrete. The discrete traits are all composed of 3 states. The first trait of each dataset, is the trait of interest on which the performance of the imputation methods is evaluated. Additionally to the first trait, the dataset includes 3 continuous and 3 discrete traits that evolve independently and 3 continuous and 3 discrete traits which are correlated to the first trait. In the study, we simulated 6 different scenarios each 100 times. Below, there is the distribution of the dataset for each scenario.

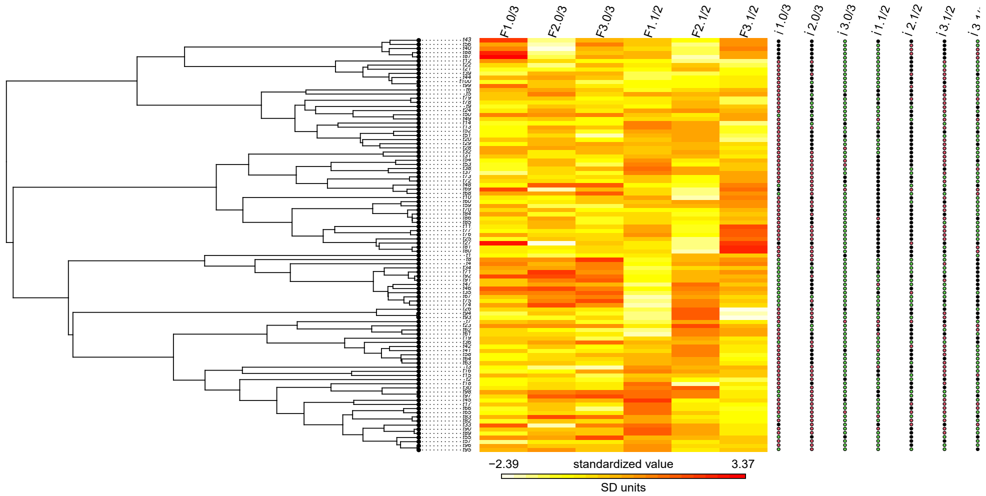

Figure 1: Distribution of simulated traits according a MK model and a phylogenetic tree build with $\lambda = 1$ and $\kappa = 1$. On the left, the phylogenetic tree used for the imputation. The tree is build with $\lambda = 1$ and $\kappa = 1$. The R functions `phylo.heatmap()` and `dotTree()` of the package phytools(*v1.0-1*; Revell (2012)) has been used to display the distribution of the continuous and categorical traits.

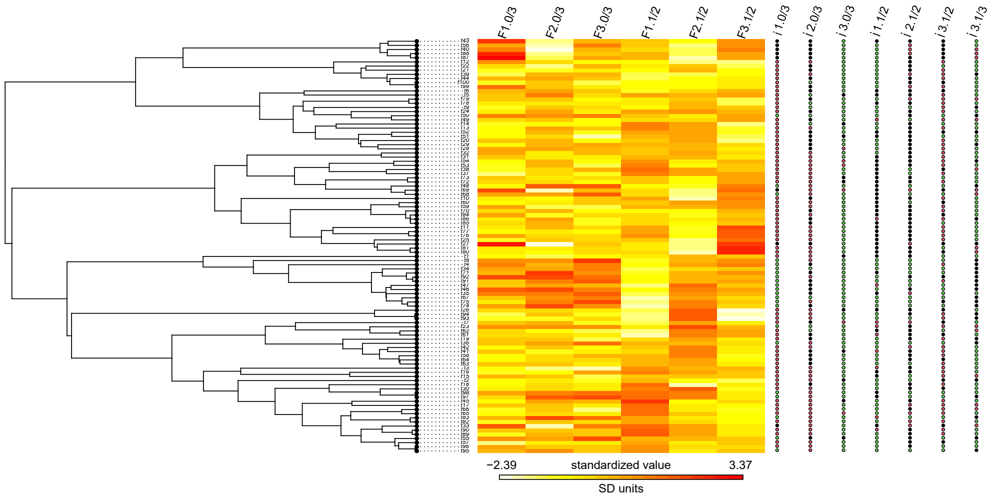

Figure 2: Distribution of simulated traits according a MK model and a phylogenetic tree build with $\lambda = 1$ and $\kappa = 0$. On the left, the phylogenetic tree used for the imputation. The tree is build with $\lambda = 1$ and $\kappa = 1$.

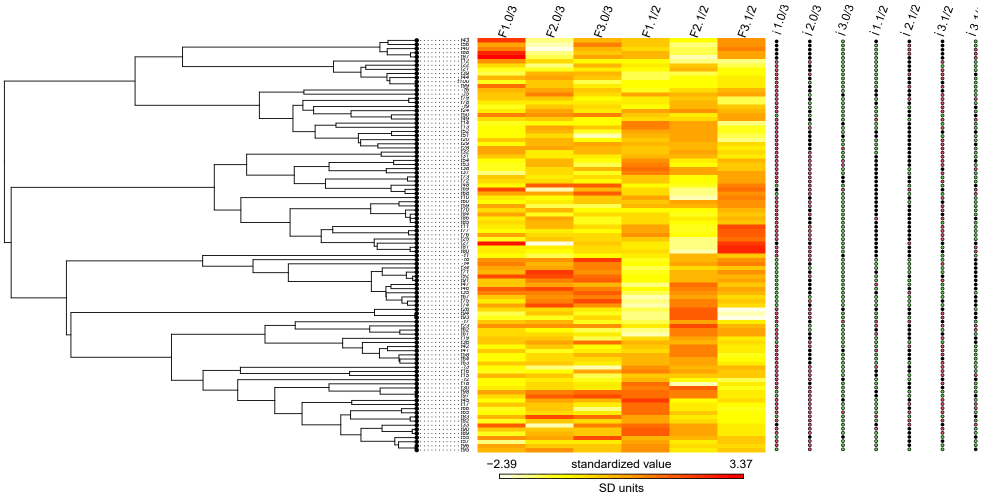

Figure 3: Distribution of simulated traits according a MK model and a phylogenetic tree build with $\lambda = 0.0001$ and $\kappa = 1$. On the left, the phylogenetic tree used for the imputation. The tree is build with $\lambda = 1$ and $\kappa = 1$.

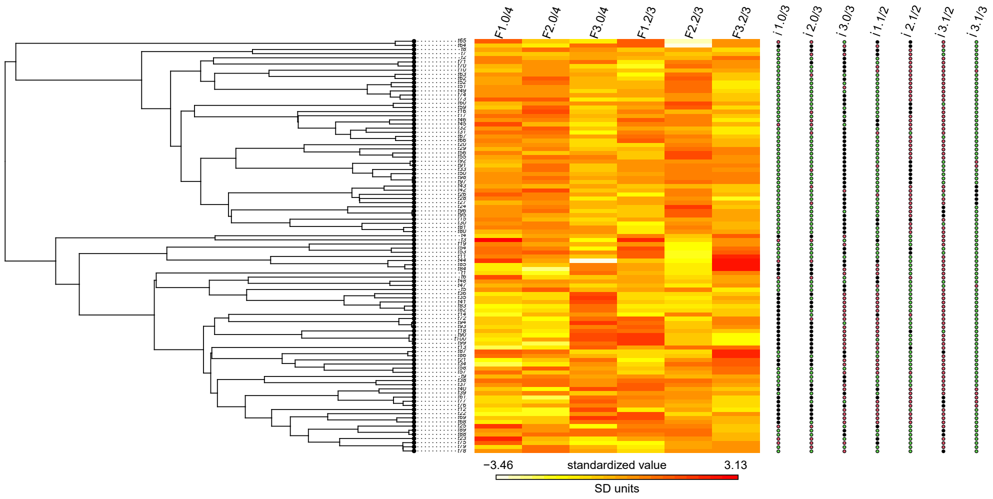

Figure 4: Distribution of simulated traits according a threshold model and a phylogenetic tree build with $\lambda = 1$ and $\kappa = 1$. On the left, the phylogenetic tree used for the imputation. The tree is build with $\lambda = 1$ and $\kappa = 1$.

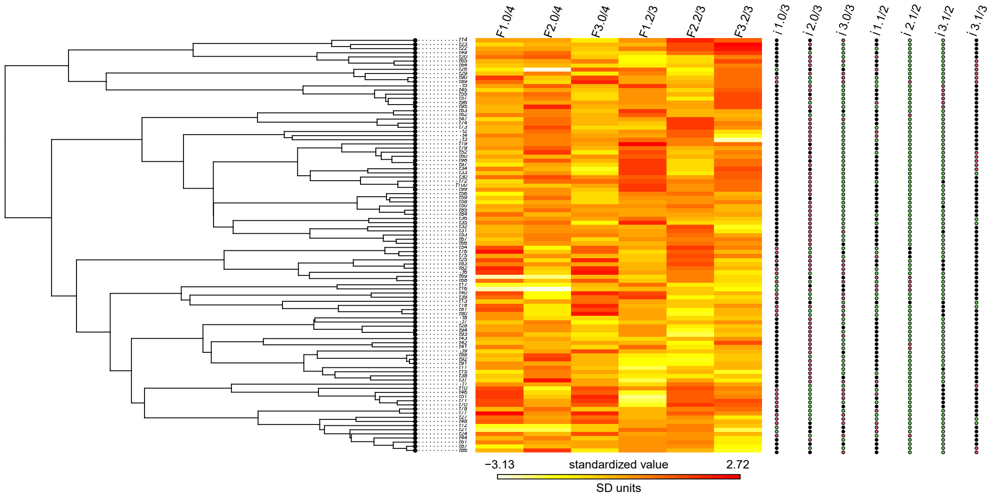

Figure 5: Distribution of simulated traits according a threshold model and a phylogenetic tree build with $\lambda = 1$ and $\kappa = 0$. On the left, the phylogenetic tree used for the imputation. The tree is build with $\lambda = 1$ and $\kappa = 1$.

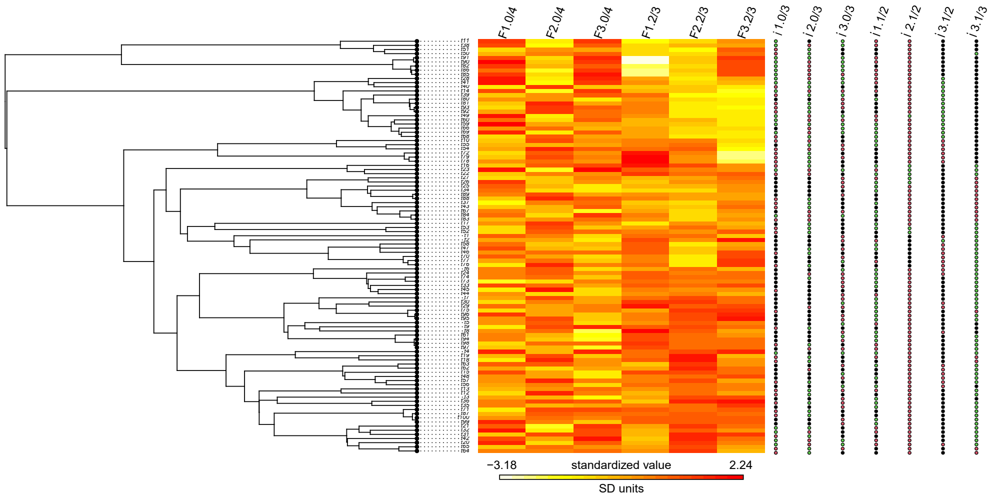

Figure 6: Distribution of simulated traits according a threshold model and a phylogenetic tree build with $\lambda = 0.0001$ and $\kappa = 1$. On the left, the phylogenetic tree used for the imputation. The tree is build with $\lambda = 1$ and $\kappa = 1$.

In addition to the Markov model, we also simulated discrete traits according to the threshold model (Felsenstein, 2005; Revell, 2014). The scenarios simulated are the same except to the model of evolution Tab. 1.

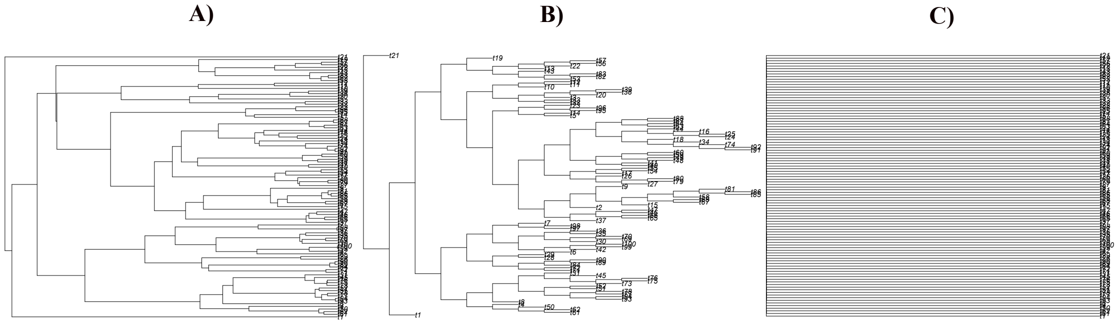

Figure 7: Illustration of the type of trees used for the different simulations. a) Phylogenetic tree with a strong phylogenetic signal. b) Tree transformed by a $\lambda = 0.0001$. c) Tree transformed by a $\kappa = 0$

Table 1: Three scenarios of how traits uncorrelated with the nominal trait of interest were generated. The correlated traits and the uncorrelated continuous traits are all simulated in the same way in the three scenarios as described in the method.

| Scenario n° | $\lambda$ value | $\kappa$ value | Evolutionary models |
|---|---|---|---|
| 1 | 1 | 1 | |
| 2 | 0.0001 | 1 | Threshold model |
| 3 | 1 | 0 | |

#### 1.2 Missing values simulation

The missing values have been simulated according 4 different missing mechanisms: missing completely at random (MCAR), missing at random (MAR), missing not at random (MNAR), and phyloNA.

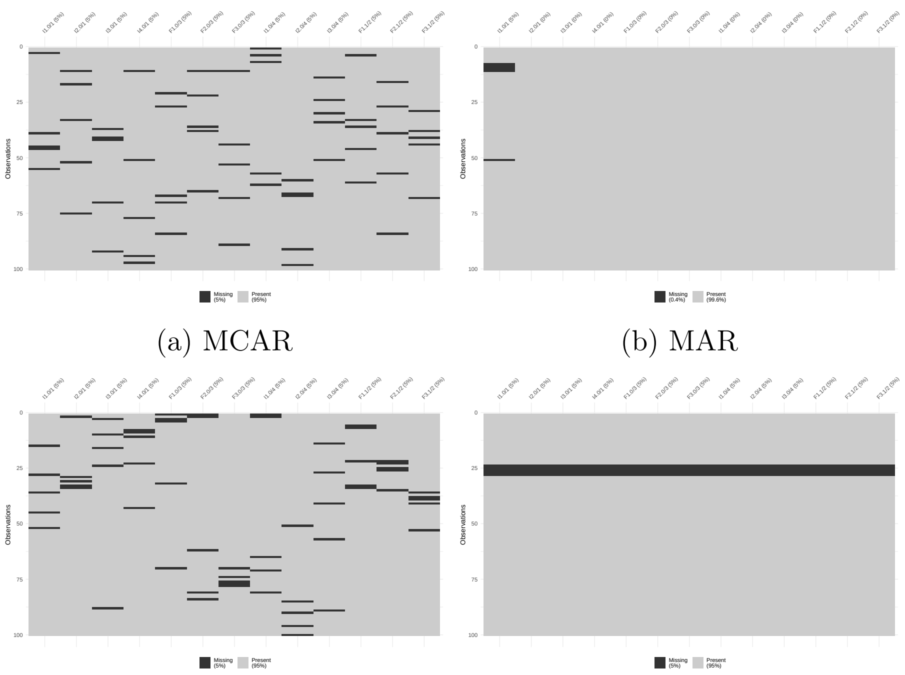

(a) MCAR (b) MAR

(c) MNAR (d) phyloNa

Figure 8: Representation of the missing values distribution in the simulated dataset under 4 missing mechanisms (MCAR, MAR, MNAR and phyloNa according to a missing rate equal to 5%. The black squares represent the missing values while the gray squares the observed values. These plots have been generated using the R function `vis_miss()` from the R package visdat 0.5.3 (Tierney, 2017)

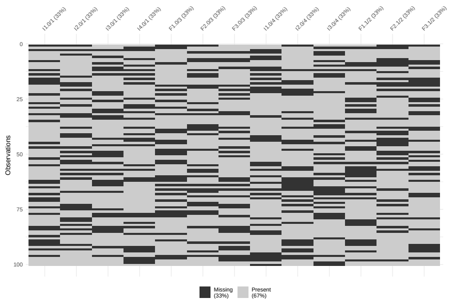

(a) MCAR

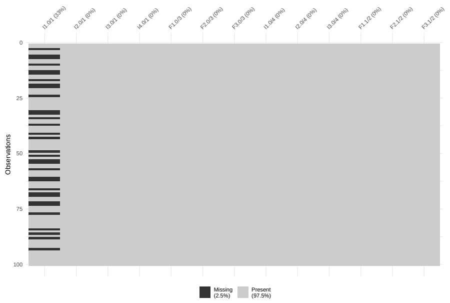

(b) MAR

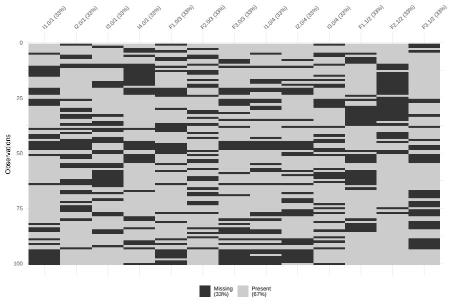

(c) MNAR

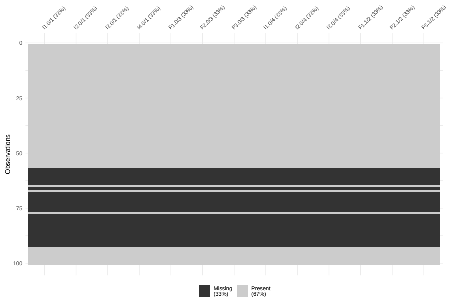

(d) phyloNa

Figure 9: Representation of the missing values distribution in the simulated dataset under 4 missing mechanisms (MCAR, MAR, MNAR and phyloNa according to a missing rate equal to 33%. The black squares represent the missing values while the gray squares the observed values. These plots have been generated using the R function `vis_miss()` from the R package visdat 0.5.3

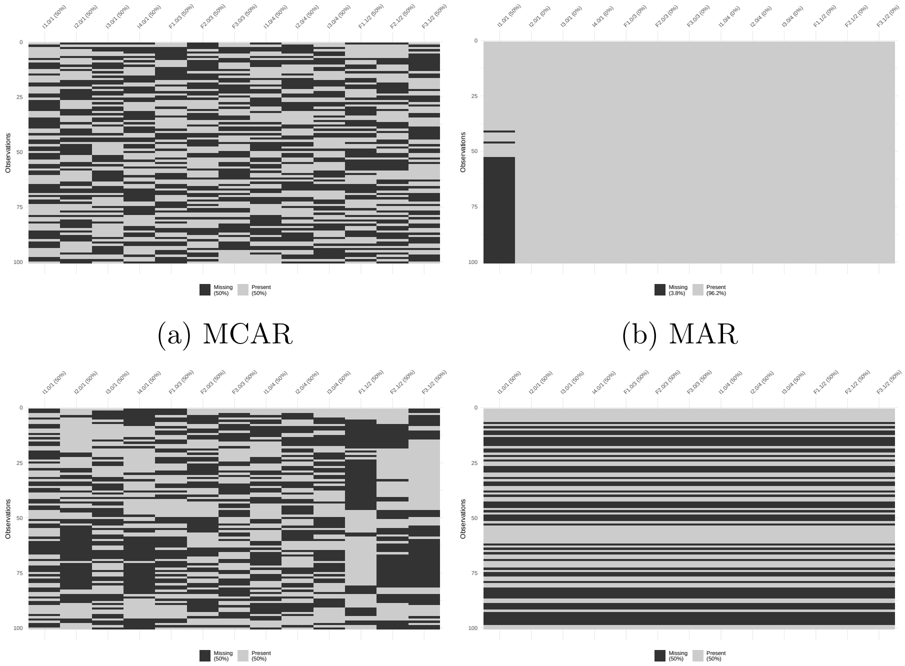

(a) MCAR (b) MAR

(c) MNAR (d) phyloNa

Figure 10: Representation of the missing values distribution in the simulated dataset under 4 missing mechanisms (MCAR, MAR, MNAR and phyloNa according to a missing rate equal to 5%. The black squares represent the missing values while the gray squares the observed values. These plots have been generated using the R function `vis_miss()` from the R package visdat0.5.3.

### 2 Imputation methods

#### 2.1 Phylogenetic imputation methods

Phylopars is a maximum likelihood phylogenetic imputation method, which is available in the R package Rphylopars 1.1.0.9004 (Goolsby et al., 2017). This method is only applicable to impute missing values of continuous variables. The approach can be described in two steps: 1) The phylogenetic covariances and the phenotypic variances are obtained through maximum likelihood estimation (MLE) according to a provided evolutionary model. The phylogenetic covariance matrix captures the evolution similarity between species (Harmon and Open Textbook Library, 2019) while the phenotypic covariance matrix describes the intraspecific variability due to a variation of the trait value among the individuals of a same species. 2) Both phenotypic and phylogenetic covariances are then used to calculate the covariances between the observed values and missing values. The imputed values are then calculated from a linear equation having as slope an estimated value from a multivariate normal distribution and as intercept a value sample from the observed values of the trait containing the missing value (Bruggeman et al., 2009).

For the study this PI method has only been used for the "2-steps" strategy. The missing value have been simulated according to the BM or the OU model. The model having the smallest Akaike information criterion (AIC) have been selected to impute the missing data. The phenotypic covariance was not considered by the function because only one individual per tips was simulated.

The phylogenetic imputation method developed in the R package corHMM 2.8(Beaulieu et al., 2013) is a method which imputes missing values through a Markov model with or without hidden states taking as input a trait dataset and a phylogenetic tree. In this study, we decided to use the method without hidden states ($rat.cat = 1$) (Beaulieu et al., 2013). The method optimizes the rates of the Q matrix using maximum likelihood estimation according to a model of evolution and then return the probability of each states of both ancestral and tips (get.tip.states = 1).
There are two main elements which composed a CTMC-FSS model, an instantaneous rate matrix (Q) a vector of state frequencies and a frequency at the root (Boyko and Beaulieu, 2021). The term finite state space means that the estimation can be of a finite number of state (3). Q gives the instantaneous transition rate from a state to another and can be modeled as ER, SYM, or ARD. From these two main elements, a phylogenetic tree is added. The phylogenetic tree is important to build the transition-probability matrix which gathers the probabilities of change between each state over a time interval *t*. (means that longer an edge is, larger the probability to see a state change is) (Boyko and Beaulieu, 2021) and the frequency at the root is estimated from the transition rate ($root.p = "yang"$).

The strategy has been to run the imputation according the three transition matrices (ER, SYM, and ARD) and to select the one with the best AIC score which means the model with the largest likelihood. Once the model selected, missing values were imputed according to the state with the highest probability. The PI methods have been applied using the `pi_continuous_traits()` function and the `pi_categorical_traits()` function from the R package TDIP 1.0 (Gendre Matthieu, 2022) which call respectively, the function `phylopars()` from the package Rphylopars 1.1.0.9004 (Goolsby et al., 2017) and the function corHMM from the package corHMM 2.8 (Beaulieu et al., 2013)).

#### 2.2 MissForest

MissForest is a non-parametric machine learning method used to impute missing values for continuous and categorical data. The method is based on the random forest (RF) algorithm which consists of the construction of an ensemble of decision trees through a two-stage randomization procedure (grow tree using bootstrap sample and random feature selection) (Breiman, 2001). The imputation is separated in two steps: 1) the RF is trained on the observed values of the dataset, and 2) The missing values are predicted using the trained RF. In more details, the missing values of a trait are imputed through a RF which have been trained with the response values being the observed values of the trait itself and the predictor values being the observed values of the other traits. These two steps procedure is iterated until a stopping criterion or the maximum of iteration which is fixed by the user is reached or if the difference between the newly imputed data and the old imputed data increase for the first time.

The missForest methods have been applied using the function (`missForest_phylo()`) from the TDIP 1.0 package (Gendre Matthieu, 2022) which apply the function `missForest()` from the R package missForest 1.4 (Stekhoven and Buhlmann, 2012). The various parameters have been set as following: in case the stopping criterion is not reached, the maximum number of iterations is equal to 10 (maxiter = 10), the number of trees designed in each forest is equal to 100 and the last parameter is the number of traits randomly sampled at each split (using the best variable/split point among the mtry) during the creation of the trees from a bootstrapped dataset. The mtry parameter is equal to the root square of the number of traits.

#### 2.3 kNN

The k nearest neighbor imputation is a non-parametric machine learning method where an aggregation of the k nearest neighbors is used to impute the missing value. The function (`kNN_phylo()`) from the TDIP 1.0 package (Gendre Matthieu, 2022) package which calls the function `kNN()` from the VIM 6.1.1 package (Kowarik and Templ, 2016) has been used. This function uses an extension of the Gower distance (Gower, 1971) which is different according the type of variable to handle. In case of nominal variables, a binary measure is applied. If the state is the same, then the distance is equal to 0 otherwise the distance is equal to 1. For continuous variable, the distance is defined as the absolute distance divided by the total range (range = 1 because traits are scaled). Once the distances calculated, the values of the k neighbors are aggregate to generate the imputed value.

For categorical traits we used the function `maxCat()` from the VIM 6.1.1 package (Kowarik and Templ, 2016). This function picks the state value of the k neighbor having the smallest distance. For continuous variables, we used the `weightedMean()` function of the laeken 0.5.2 package (Alfons and Templ, 2013) which consists of filling the missing value computing the weighted mean of the k neighbors. For the study, $k$ has been set to 2. The function does not take in account the imputed value of other traits or species to impute the next missing values.

#### 2.4 GAIN

Generative Adversarial Imputation Nets (GAIN) is a generative unsupervised deep learning method. This approach is an adaptation of the Generative Adversarial Nets (GAN) framework. GAIN is composed of two neural networks, a Generator and a Discriminator which are trained against each other. The Generator ($G$) is a neural network (NN) that aims to impute missing data. It takes as input a matrix composed of observed and unobserved values from the true data ($\tilde{X}$), a binary matrix ($M$) whose 1 corresponds to the components of the matrix which are observed and a matrix of noise ($Z$). Those three matrices have the same size of the true dataset ($X$) (the dataset in which we want to impute the missing values). The output of the NN is a matrix where the elements equal to 0 in the second input matrix are imputed ($\hat{X}$). The Discriminator($D$) is the second NN of the approach. The goal of this NN is to discriminate which value from the Generator matrix is imputed or observed in the true dataset. The input of $D$ is the $\hat{X}$ matrix and a hint matrix ($H$). The hint is a matrix of the same size than $M$ and is composed of three values, 1, 0 and 0.5. The aim of H is to reveal to D which value is imputed or observed in the original dataset. The value 0.5 means that the hint does not know if the value is imputed or observed. The output matrix is a probability matrix whose value is closer to 1 the more the value $i$ of $\hat{X}$ is considered as observed. The generator is trained to minimize the probability that the discriminator predicts $M$ while the discriminator is trained to minimize the probability to correctly predict $M$. The model returns three loss functions, one for the generator, one for the discriminator and one quantifying differences between the true and the imputed data. The R function which imputes values according the GAIN method is called `gain_phylo()` from the TDIP 1.0 package (Gendre Matthieu, 2022). The inputs are: 1) the dataset containing missing values in which, the categorical traits are converted in one-hot encoding variables using the R function *one_hot()* from the package mltools 0.3.5 (Ben Gorman, 2018), 2) the amount of phylogenetic variance that we want to include, 3) a phylogenetic tree, 4) the batch size, 5) the hint rate, 6) the hyperparameter alpha, and 7) the number of iterations.

For the study, we set the batch size to the 20% of the total number of traits, the hint rate to 90%, which is the amount of the mask that the hint provides to the discriminator, the hyperparameter equal to 100, and the epochs equal to 10'000. The GAIN algorithm is coded in python and available on the GitHub page from (Yoon et al., 2018). We have made some changes to the algorithm provided. Firstly, the algorithm is now a python class. Secondly, we defined the H matrix as defined in the original paper. The B matrix is thus draw from a binomial distribution. We used the R package reticulate 1.24 (Ushey et al., 2022) to link the python code to the R framework.

#### 2.5 MICE

Multivariate imputation by chained equations (MICE) is an approach which creates multiple imputation of missing values being categorical or continuous according to an imputation method. The methods used to impute the missing values are different. For our study, we used the default methods, which are predictive mean matching (PMM) in case of continuous variables and/or categorical variables having more than 20 different states, logistic regression in case of binary variables, multinomial logistic regression in case of categorical variable with more than 2 states, and proportional odds logistic regression for ordinal variables. Because the simulate dataset contains only nominal variables having 3 states and continuous traits, the method used are multinomial logistic regression and PMM. The parameters of the imputation models are estimated from the observed values of the dataset in an iterative way.

Multinomial logistic regression is a model being part of the generalized linear model class (Buuren and Groothuis-Oudshoorn, 2011). More precisely, the model used to impute the missing values is the Bayesian polytomous regression model. The algorithm works as follow, firstly, it fits categorical responses as a multinomial model using the `multinom()` function from the R package nnet 7.3-15(Venables and Ripley, 2002) which applies multinomial log-linear models via neural networks and secondly, from the estimated model, computes the missing categories(states). The model assumes that the parameters of the model ($\beta$) follow a multivariate normal distribution (Buuren and Groothuis-Oudshoorn, 2011). The missing values present in a continuous variable are imputed through the predictive mean matching (PMM) method. PMM is a semi-parametric method which is similar to a regression model except that the missing value are not filled according to a linear model prediction but are filled with values that are present in the dataset and are close to the linear model prediction (Buuren and Groothuis-Oudshoorn, 2011) (empirical evaluation). PMM has the propriety that it ensures that the value imputed are plausible (Buuren and Groothuis-Oudshoorn, 2011).

The function `mice_phylo()` from the package TDIP 1.0 (Gendre Matthieu, 2022) which use the function `mice()` from the R package MICE 3.14.0 (Buuren and Groothuis-Oudshoorn, 2011) has been used. The function takes as arguments in addition of the dataset with missing values, the number of imputations which is equal to 1 and the maximum number of iterations which we set to 5 (default value). Once the imputed dataset is generated, the algorithm apply the function `complete()` from the MICE package which replaces the traits containing missing values with the corresponding imputed traits. The way that we use MICE is simply to test the imputation method implemented in the packages and not to use the entire MICE framework as described in the literature (Buuren and Groothuis-Oudshoorn, 2011).

### 3 Implementation

The pipeline is build around the TDIP 1.0(Gendre Matthieu, 2022) package that we build for the study. This package calls many functions from various packages. In this section we will list all the functions used in the different stages of the pipeline.

A)To simulate the trait data and the phylogenetic tree, we implement the function `data_simulator()` from the TDIP package. To simulate the trees, the function calls `pbtree()` function from the R package phytools 1.2-0 (Revell, 2012). The tree are rescaled using the function `rescale()` from the package geiger 2.0.10 (Alfons and Templ, 2013). For the simulation of the trait data, the uncorrelated continuous traits are simulated with the `mvSim()`) function from the mvMORPH 1.1.6 (Clavel et al., 2015) package, the uncorrelated discrete traits with the `get_random_mk_transition_matrix()` and `simulate_mk_model()` function from the R package castor 1.7.3 (Louca and Doebeli, 2018). The continuous traits correlated to the trait of interest are simulated using the function `rnorm_pre()` from the faux 1.1.0.9004 (DeBruine, 2021) package.

B) Once the trait data simulated, the second step is to generate missing values. The missing values can be MCAR, MAR, MNAR, or phyloNA. We implemented the `na_insertion()` function of the TDIP package that generates these four type of missing values at once. For the generation of MCAR, MAR and MNAR values the R package missMethods 0.40 (Rockel, 2022).

C) The third step is the imputation. To impute data, we wrote the function `missing_data_imputation()` from the TDIP package which impute data according to the strategies and the method that the user provides. The TDIP package provides also a way to apply each imputation method separately. Once the data imputed, we used the function `hard_voting()` from the TDIP package to apply the ensemble method. The code of the pipeline is available on GitHub [1] and the TDIP source code [2] too.

### 4 Outputs of the simulations

#### 4.1 Accuracy tables

In this section, you will find all the tables displaying the imputation accuracy of the various methods tested in the study.

Table 2: Average accuracy and standard deviation in parenthesis of imputations across all scenarios with data simulated by a MK model. The missing rate is 5%. Sample size is 600 for each imputation method.

| Methods | No phylogeny | With phylogeny | 2-steps | Ensemble |
|---|---|---|---|---|
| PI | - | 0.641(0.345) | - | - |
| MICE | 0.614(0.314) | 0.535(0.322) | 0.601(0.323) | - |
| KNN | 0.661(0.347) | 0.682(0.317) | 0.726(0.311) | - |
| MissForest | 0.713(0.327) | 0.696(0.31) | 0.706(0.322) | - |
| GAIN | 0.543(0.323) | 0.546(0.321) | 0.54(0.321) | - |
| HV | - | - | - | 0.735(0.315) |

Table 3: Average accuracy and standard deviation in parenthesis of imputations across all scenarios with data simulated by a MK model. The missing rate is 33%. Sample size is 600 for each imputation method.

| Methods | No phylogeny | With phylogeny | 2-steps | Ensemble |
|---|---|---|---|---|
| PI | - | 0.62(0.283) | - | - |
| MICE | 0.529(0.223) | 0.473(0.211) | 0.53(0.235) | - |
| KNN | 0.61(0.276) | 0.638(0.25) | 0.683(0.257) | - |
| MissForest | 0.663(0.272) | 0.65(0.255) | 0.673(0.265) | - |
| GAIN | 0.51(0.235) | 0.525(0.238) | 0.517(0.232) | - |
| HV | - | - | - | 0.689(0.265) |

[1] https://github.com/Matgend/Phylo_Imputation
[2] https://github.com/Matgend/TDIP

Table 4: Average accuracy and standard deviation in parenthesis of imputations across all scenarios with data simulated by a MK model. The missing rate is 50%. Sample size is 600 for each imputation method.

| Methods | No phylogeny | With phylogeny | 2-steps | Ensemble |
|---|---|---|---|---|
| PI | - | 0.599(0.283) | - | - |
| MICE | 0.472(0.193) | 0.446(0.176) | 0.502(0.208) | - |
| KNN | 0.58(0.262) | 0.61(0.244) | 0.649(0.254) | - |
| MissForest | 0.627(0.268) | 0.623(0.252) | 0.649(0.267) | - |
| GAIN | 0.495(0.232) | 0.504(0.23) | 0.49(0.227) | - |
| HV | - | - | - | 0.654(0.262) |

Table 5: Table representing the average accuracy and the standard deviation obtained from the imputation of all the scenarios from data simulated by a MK model and containing strongly correlated traits ($\rho = 0.8$). The missing rate is 5%. Sample size is 600 for each imputation method.

| Methods | No phylogeny | With phylogeny | 2-steps | Ensemble |
|---|---|---|---|---|
| PI | - | 0.64(0.349) | - | - |
| MICE | 0.752(0.287) | 0.632(0.331) | 0.72(0.291) | - |
| KNN | 0.778(0.318) | 0.757(0.284) | 0.827(0.247) | - |
| MissForest | 0.857(0.243) | 0.783(0.275) | 0.776(0.286) | - |
| GAIN | 0.595(0.323) | 0.587(0.308) | 0.59(0.317) | - |
| HV | - | - | - | 0.87(0.226) |

Table 6: Average accuracy and standard deviation in parenthesis of imputations across all scenarios with data simulated by a MK model and containing strongly correlated traits ($\rho = 0.8$). The missing rate is 33%. Sample size is 600 for each imputation method.

| Methods | No phylogeny | With phylogeny | 2-steps | Ensemble |
|---|---|---|---|---|
| PI | - | 0.615(0.288) | - | - |
| MICE | 0.638(0.213) | 0.547(0.217) | 0.629(0.214) | - |
| KNN | 0.689(0.276) | 0.699(0.234) | 0.756(0.228) | - |
| MissForest | 0.778(0.237) | 0.704(0.247) | 0.719(0.253) | - |
| GAIN | 0.543(0.24) | 0.554(0.235) | 0.539(0.233) | - |
| HV | - | - | - | 0.795(0.22) |

Table 7: Table representing the average accuracy and the standard deviation obtained from the imputation of all the scenarios from data simulated by a MK model and containing strongly correlated traits ($\rho = 0.8$). The missing rate is 50%. Sample size is 600 for each imputation method.

| Methods | No phylogeny | With phylogeny | 2-steps | Ensemble |
|---|---|---|---|---|
| PI | - | 0.583(0.289) | - | - |
| MICE | 0.559(0.201) | 0.508(0.186) | 0.573(0.192) | - |
| KNN | 0.634(0.268) | 0.651(0.238) | 0.698(0.241) | - |
| MissForest | 0.724(0.256) | 0.648(0.258) | 0.675(0.268) | - |
| GAIN | 0.518(0.239) | 0.515(0.23) | 0.508(0.232) | - |
| HV | - | - | - | 0.737(0.243) |

Table 8: Table representing the average accuracy and the standard deviation obtained from the imputation of all the scenarios from data simulated by a threshold model and containing strongly correlated traits ($\rho = 0.8$). and contained strongly correlated traits ($\rho = 0.8$). The missing rate is 5%. Sample size is 600 for each imputation method.

| Methods | No phylogeny | With phylogeny | 2-steps | Ensemble |
|---|---|---|---|---|
| PI | - | 0.618(0.333) | - | - |
| MICE | 0.755(0.291) | 0.659(0.325) | 0.733(0.282) | - |
| KNN | 0.786(0.315) | 0.758(0.291) | 0.822(0.261) | - |
| MissForest | 0.841(0.273) | 0.798(0.278) | 0.764(0.288) | - |
| GAIN | 0.581(0.307) | 0.576(0.308) | 0.574(0.314) | - |
| HV | - | - | - | 0.853(0.247) |

Table 9: Table representing the average accuracy and the standard deviation obtained from the imputation of all the scenarios from Data simulated by a threshold model and containing strongly correlated traits ($\rho = 0.8$). The missing rate is 33%. Sample size is 600 for each imputation method.

| Methods | No phylogeny | With phylogeny | 2-steps | Ensemble |
|---|---|---|---|---|
| PI | - | 0.557(0.272) | - | - |
| MICE | 0.656(0.221) | 0.577(0.207) | 0.64(0.185) | - |
| KNN | 0.693(0.261) | 0.683(0.221) | 0.734(0.226) | - |
| MissForest | 0.754(0.249) | 0.671(0.256) | 0.666(0.265) | - |
| GAIN | 0.507(0.225) | 0.53(0.233) | 0.516(0.225) | - |
| HV | - | - | - | 0.771(0.221) |

Table 10: Table representing the average accuracy and the standard deviation obtained from the imputation of all the scenarios from Data simulated by a threshold model and containing strongly correlated traits ($\rho = 0.8$). The missing rate is 50%. Sample size is 600 for each imputation method.

| Methods | No phylogeny | With phylogeny | 2-steps | Ensemble |
|---|---|---|---|---|
| PI | - | 0.532(0.264) | - | - |
| MICE | 0.574(0.205) | 0.534(0.18) | 0.583(0.172) | - |
| KNN | 0.609(0.262) | 0.622(0.233) | 0.657(0.247) | - |
| MissForest | 0.683(0.281) | 0.614(0.263) | 0.623(0.276) | - |
| GAIN | 0.488(0.23) | 0.494(0.226) | 0.486(0.224) | - |
| HV | - | - | - | 0.694(0.256) |

Table 11: Table representing the average accuracy and the standard deviation obtained from the imputation of all the scenarios from data simulated by a MK model and containing only independent traits. The missing rate is 33%. Sample size is 600 for each imputation method.

| Methods | No phylogeny | With phylogeny | 2-steps | Ensemble |
|---|---|---|---|---|
| PI | - | 0.619(0.285) | - | - |
| MICE | 0.435(0.175) | 0.406(0.172) | 0.437(0.217) | - |
| KNN | 0.527(0.257) | 0.575(0.257) | 0.604(0.272) | - |
| MissForest | 0.542(0.257) | 0.595(0.257) | 0.621(0.279) | - |
| GAIN | 0.465(0.225) | 0.487(0.235) | 0.473(0.225) | - |
| HV | - | - | - | 0.581(0.269) |

Table 12: Table representing the average accuracy and the standard deviation obtained from the imputation of MCAR values. The missing rate is 5%. MK corresponds to the MK model and TM to the threshold model. Sample size of each scenario is 100 for each imputation method. The data contained some strongly correlated traits ($\rho = 0.8$).

| Methods | MK | MK, $\kappa = 0$ | MK, $\lambda = 0.0001$ | TM | TM, $\kappa = 0$ | TM, $\lambda = 0.0001$ |
|---|---|---|---|---|---|---|
| PI | 0.936(0.098) | 0.542(0.248) | 0.692(0.215) | 0.792(0.205) | 0.732(0.219) | 0.574(0.295) |
| MICE | 0.862(0.179) | 0.902(0.132) | 0.86(0.162) | 0.902(0.126) | 0.886(0.14) | 0.892(0.154) |
| MissForest | 0.974(0.068) | 0.966(0.076) | 0.964(0.082) | 0.974(0.068) | 0.96(0.094) | 0.966(0.086) |
| KNN | 0.956(0.083) | 0.948(0.093) | 0.926(0.123) | 0.948(0.097) | 0.948(0.097) | 0.942(0.107) |
| GAIN | 0.908(0.146) | 0.688(0.206) | 0.702(0.239) | 0.78(0.19) | 0.796(0.184) | 0.694(0.223) |
| MICE+P | 0.77(0.215) | 0.828(0.18) | 0.69(0.235) | 0.826(0.17) | 0.81(0.208) | 0.79(0.191) |
| MissForest+P | 0.964(0.077) | 0.89(0.134) | 0.884(0.148) | 0.928(0.112) | 0.936(0.124) | 0.904(0.138) |
| KNN+P | 0.958(0.087) | 0.828(0.184) | 0.86(0.152) | 0.912(0.137) | 0.888(0.154) | 0.834(0.171) |
| GAIN+P | 0.874(0.155) | 0.672(0.232) | 0.71(0.219) | 0.792(0.191) | 0.784(0.216) | 0.702(0.221) |
| MICE+PI | 0.89(0.157) | 0.824(0.198) | 0.738(0.218) | 0.88(0.173) | 0.848(0.161) | 0.774(0.236) |
| MissForest+PI | 0.968(0.074) | 0.908(0.143) | 0.862(0.167) | 0.932(0.121) | 0.934(0.133) | 0.892(0.149) |
| KNN+PI | 0.962(0.079) | 0.936(0.106) | 0.906(0.143) | 0.95(0.096) | 0.95(0.096) | 0.95(0.096) |
| GAIN+PI | 0.868(0.169) | 0.698(0.261) | 0.726(0.227) | 0.79(0.2) | 0.742(0.226) | 0.706(0.228) |
| HV | 0.978(0.063) | 0.966(0.076) | 0.958(0.1) | 0.968(0.079) | 0.966(0.086) | 0.968(0.079) |

Table 13: Table representing the average accuracy and the standard deviation obtained from the imputation of MAR values. The missing rate is 5%. MK corresponds to the MK model and TM to the threshold model. Sample size of each scenario is 100 for each imputation method. The data contained some strongly correlated traits ($\rho = 0.8$).

| Methods | MK | MK, $\kappa = 0$ | MK, $\lambda = 0.0001$ | TM | TM, $\kappa = 0$ | TM, $\lambda = 0.0001$ |
|---|---|---|---|---|---|---|
| PI | 0.804(0.259) | 0.466(0.33) | 0.606(0.326) | 0.664(0.286) | 0.63(0.313) | 0.472(0.357) |
| MICE | 0.87(0.181) | 0.926(0.135) | 0.888(0.164) | 0.884(0.171) | 0.9(0.162) | 0.894(0.162) |
| MissForest | 0.954(0.11) | 0.954(0.098) | 0.952(0.107) | 0.956(0.105) | 0.958(0.111) | 0.956(0.101) |
| KNN | 0.922(0.139) | 0.934(0.114) | 0.924(0.12) | 0.916(0.143) | 0.92(0.148) | 0.924(0.136) |
| GAIN | 0.806(0.242) | 0.594(0.304) | 0.704(0.291) | 0.708(0.261) | 0.68(0.274) | 0.618(0.305) |
| MICE+P | 0.68(0.309) | 0.806(0.239) | 0.678(0.271) | 0.732(0.279) | 0.758(0.263) | 0.722(0.29) |
| MissForest+P | 0.87(0.202) | 0.864(0.188) | 0.782(0.245) | 0.88(0.197) | 0.85(0.245) | 0.832(0.244) |
| KNN+P | 0.868(0.197) | 0.814(0.215) | 0.806(0.21) | 0.854(0.218) | 0.858(0.242) | 0.756(0.279) |
| GAIN+P | 0.794(0.24) | 0.6(0.276) | 0.648(0.254) | 0.716(0.262) | 0.642(0.282) | 0.64(0.284) |
| MICE+PI | 0.832(0.212) | 0.838(0.194) | 0.67(0.255) | 0.812(0.214) | 0.824(0.195) | 0.782(0.277) |
| MissForest+PI | 0.84(0.234) | 0.708(0.317) | 0.636(0.321) | 0.73(0.258) | 0.716(0.279) | 0.686(0.335) |
| KNN+PI | 0.9(0.181) | 0.89(0.162) | 0.826(0.212) | 0.888(0.187) | 0.858(0.221) | 0.85(0.215) |
| GAIN+PI | 0.776(0.255) | 0.582(0.293) | 0.652(0.304) | 0.718(0.258) | 0.64(0.307) | 0.614(0.331) |
| HV | 0.952(0.114) | 0.958(0.1) | 0.934(0.117) | 0.95(0.111) | 0.946(0.117) | 0.926(0.16) |

Table 14: Table representing the average accuracy and the standard deviation obtained from the imputation of MNAR values. The missing rate is 5%. MK corresponds to the MK model and TM to the threshold model. Sample size of each scenario is 100 for each imputation method. The data contained some strongly correlated traits ($\rho = 0.8$).

| Methods | MK | MK, $\kappa = 0$ | MK, $\lambda = 0.0001$ | TM | TM, $\kappa = 0$ | TM, $\lambda = 0.0001$ |
|---|---|---|---|---|---|---|
| PI | 0.788(0.306) | 0.308(0.379) | 0.464(0.481) | 0.69(0.326) | 0.556(0.367) | 0.33(0.412) |
| MICE | 0.846(0.188) | 0.92(0.145) | 0.842(0.183) | 0.878(0.161) | 0.878(0.193) | 0.858(0.18) |
| MissForest | 0.856(0.258) | 0.914(0.166) | 0.844(0.245) | 0.884(0.242) | 0.874(0.244) | 0.878(0.244) |
| KNN | 0.808(0.29) | 0.878(0.195) | 0.834(0.261) | 0.86(0.266) | 0.836(0.266) | 0.85(0.272) |
| GAIN | 0.728(0.281) | 0.478(0.288) | 0.48(0.334) | 0.586(0.318) | 0.546(0.324) | 0.468(0.304) |
| MICE+P | 0.796(0.226) | 0.85(0.162) | 0.706(0.281) | 0.82(0.235) | 0.8(0.231) | 0.792(0.216) |
| MissForest+P | 0.756(0.317) | 0.724(0.302) | 0.622(0.395) | 0.802(0.303) | 0.802(0.305) | 0.73(0.372) |
| KNN+P | 0.762(0.336) | 0.678(0.288) | 0.658(0.353) | 0.792(0.287) | 0.77(0.288) | 0.682(0.341) |
| GAIN+P | 0.686(0.283) | 0.458(0.275) | 0.492(0.331) | 0.6(0.305) | 0.52(0.289) | 0.486(0.287) |
| MICE+PI | 0.87(0.2) | 0.822(0.213) | 0.76(0.254) | 0.824(0.25) | 0.782(0.231) | 0.75(0.238) |
| MissForest+PI | 0.832(0.259) | 0.794(0.309) | 0.638(0.389) | 0.81(0.289) | 0.768(0.32) | 0.762(0.315) |
| KNN+PI | 0.794(0.307) | 0.84(0.259) | 0.748(0.317) | 0.838(0.285) | 0.814(0.289) | 0.794(0.297) |
| GAIN+PI | 0.69(0.307) | 0.484(0.298) | 0.486(0.324) | 0.622(0.307) | 0.554(0.301) | 0.452(0.306) |
| HV | 0.864(0.259) | 0.888(0.211) | 0.842(0.258) | 0.874(0.258) | 0.85(0.261) | 0.854(0.244) |

Table 15: Table representing the average accuracy and the standard deviation obtained from the imputation of phyloNA values. The missing rate is 5%. MK corresponds to the MK model and TM to the threshold model. Sample size of each scenario is 100 for each imputation method. The data contained some strongly correlated traits ($\rho = 0.8$).

| Methods | MK | MK, $\kappa = 0$ | MK, $\lambda = 0.0001$ | TM | TM, $\kappa = 0$ | TM, $\lambda = 0.0001$ |
|---|---|---|---|---|---|---|
| PI | 0.904(0.183) | 0.496(0.304) | 0.678(0.246) | 0.732(0.298) | 0.664(0.324) | 0.576(0.269) |
| MICE | 0.394(0.233) | 0.302(0.204) | 0.396(0.209) | 0.348(0.247) | 0.35(0.235) | 0.38(0.217) |
| MissForest | 0.686(0.38) | 0.51(0.282) | 0.7(0.217) | 0.552(0.367) | 0.546(0.367) | 0.582(0.265) |
| KNN | 0.414(0.409) | 0.366(0.279) | 0.416(0.329) | 0.454(0.369) | 0.424(0.37) | 0.4(0.295) |
| GAIN | 0.394(0.301) | 0.312(0.245) | 0.328(0.242) | 0.396(0.273) | 0.324(0.249) | 0.354(0.241) |
| MICE+P | 0.35(0.316) | 0.268(0.25) | 0.168(0.188) | 0.322(0.287) | 0.33(0.281) | 0.21(0.194) |
| MissForest+P | 0.834(0.256) | 0.51(0.289) | 0.666(0.224) | 0.674(0.303) | 0.662(0.317) | 0.562(0.263) |
| KNN+P | 0.822(0.274) | 0.486(0.283) | 0.528(0.3) | 0.658(0.322) | 0.6(0.345) | 0.476(0.294) |
| GAIN+P | 0.36(0.291) | 0.336(0.236) | 0.398(0.272) | 0.356(0.276) | 0.336(0.28) | 0.344(0.254) |
| MICE+PI | 0.78(0.245) | 0.316(0.226) | 0.27(0.196) | 0.606(0.307) | 0.572(0.304) | 0.332(0.255) |
| MissForest+PI | 0.878(0.222) | 0.524(0.277) | 0.694(0.215) | 0.704(0.308) | 0.666(0.315) | 0.57(0.272) |
| KNN+PI | 0.886(0.209) | 0.54(0.284) | 0.672(0.242) | 0.72(0.306) | 0.672(0.306) | 0.566(0.276) |
| GAIN+PI | 0.368(0.299) | 0.344(0.24) | 0.39(0.233) | 0.334(0.277) | 0.338(0.272) | 0.362(0.247) |
| HV | 0.888(0.196) | 0.516(0.287) | 0.68(0.234) | 0.702(0.308) | 0.658(0.31) | 0.566(0.279) |

Table 16: Table representing the average accuracy and the standard deviation obtained from the imputation of MCAR values. The missing rate is 33%. MK corresponds to the MK model and TM to the threshold model. Sample size of each scenario is 100 for each imputation method. The data contained some strongly correlated traits ($\rho = 0.8$).

| Methods | MK | MK, $\kappa = 0$ | MK, $\lambda = 0.0001$ | TM | TM, $\kappa = 0$ | TM, $\lambda = 0.0001$ |
|---|---|---|---|---|---|---|
| PI | 0.897(0.136) | 0.512(0.16) | 0.682(0.144) | 0.756(0.142) | 0.708(0.145) | 0.564(0.191) |
| MICE | 0.758(0.114) | 0.796(0.085) | 0.73(0.091) | 0.801(0.1) | 0.785(0.089) | 0.791(0.092) |
| MissForest | 0.939(0.051) | 0.905(0.06) | 0.926(0.054) | 0.918(0.059) | 0.912(0.064) | 0.921(0.053) |
| KNN | 0.823(0.097) | 0.735(0.098) | 0.784(0.092) | 0.772(0.093) | 0.77(0.097) | 0.762(0.11) |
| GAIN | 0.765(0.111) | 0.516(0.153) | 0.582(0.148) | 0.651(0.126) | 0.63(0.147) | 0.549(0.147) |
| MICE+P | 0.718(0.126) | 0.705(0.101) | 0.578(0.124) | 0.738(0.109) | 0.713(0.101) | 0.637(0.119) |
| MissForest+P | 0.928(0.055) | 0.753(0.109) | 0.77(0.103) | 0.881(0.075) | 0.857(0.08) | 0.77(0.107) |
| KNN+P | 0.887(0.067) | 0.671(0.112) | 0.749(0.112) | 0.819(0.083) | 0.797(0.091) | 0.696(0.125) |
| GAIN+P | 0.791(0.102) | 0.51(0.134) | 0.575(0.149) | 0.729(0.109) | 0.662(0.121) | 0.537(0.147) |
| MICE+PI | 0.839(0.124) | 0.713(0.096) | 0.643(0.116) | 0.808(0.101) | 0.771(0.106) | 0.677(0.122) |
| MissForest+PI | 0.948(0.047) | 0.833(0.089) | 0.803(0.1) | 0.89(0.082) | 0.864(0.089) | 0.827(0.11) |
| KNN+PI | 0.932(0.05) | 0.802(0.086) | 0.82(0.073) | 0.885(0.068) | 0.867(0.067) | 0.819(0.084) |
| GAIN+PI | 0.752(0.114) | 0.512(0.128) | 0.598(0.132) | 0.684(0.117) | 0.646(0.127) | 0.562(0.147) |
| HV | 0.955(0.045) | 0.88(0.068) | 0.911(0.056) | 0.924(0.056) | 0.908(0.056) | 0.896(0.06) |

Table 17: Tables representing the average accuracy and the standard deviation obtained from the imputation of MCAR values. The missing rate is 33%. MK corresponds to the MK model. Sample size of each scenario is 100 for each imputation method. **(a)** Data contained some strongly correlated traits ($\rho = 0.8$). **(b)** Data composed exclusively of independent traits.

(a) Strong correlation ($\rho = 0.8$)

| Methods | MK | MK, $\kappa = 0$ | MK, $\lambda = 0.0001$ |
|---|---|---|---|
| PI | 0.897(0.136) | 0.512(0.16) | 0.682(0.144) |
| MICE | 0.758(0.114) | 0.796(0.085) | 0.73(0.091) |
| MissForest | 0.939(0.051) | 0.905(0.06) | 0.926(0.054) |
| KNN | 0.823(0.097) | 0.735(0.098) | 0.784(0.092) |
| GAIN | 0.765(0.111) | 0.516(0.153) | 0.582(0.148) |
| MICE+P | 0.718(0.126) | 0.705(0.101) | 0.578(0.124) |
| MissForest+P | 0.928(0.055) | 0.753(0.109) | 0.77(0.103) |
| KNN+P | 0.887(0.067) | 0.671(0.112) | 0.749(0.112) |
| GAIN+P | 0.791(0.102) | 0.51(0.134) | 0.575(0.149) |
| MICE+PI | 0.839(0.124) | 0.713(0.096) | 0.643(0.116) |
| MissForest+PI | 0.948(0.047) | 0.833(0.089) | 0.803(0.1) |
| KNN+PI | 0.932(0.05) | 0.802(0.086) | 0.82(0.073) |
| GAIN+PI | 0.752(0.114) | 0.512(0.128) | 0.598(0.132) |
| HV | 0.955(0.045) | 0.88(0.068) | 0.911(0.056) |

(b) Low correlation

| Methods | MK | MK, $\kappa = 0$ | MK, $\lambda = 0.0001$ |
|---|---|---|---|
| PI | 0.924(0.062) | 0.499(0.155) | 0.673(0.165) |
| MICE | 0.61(0.114) | 0.395(0.109) | 0.489(0.144) |
| MissForest | 0.788(0.104) | 0.456(0.142) | 0.595(0.175) |
| KNN | 0.711(0.138) | 0.442(0.15) | 0.614(0.163) |
| GAIN | 0.74(0.113) | 0.409(0.123) | 0.516(0.152) |
| MICE+P | 0.688(0.123) | 0.416(0.098) | 0.417(0.093) |
| MissForest+P | 0.896(0.069) | 0.507(0.137) | 0.589(0.164) |
| KNN+P | 0.835(0.083) | 0.472(0.145) | 0.626(0.152) |
| GAIN+P | 0.788(0.102) | 0.424(0.122) | 0.548(0.157) |
| MICE+PI | 0.766(0.139) | 0.365(0.115) | 0.387(0.112) |
| MissForest+PI | 0.918(0.063) | 0.498(0.153) | 0.677(0.152) |
| KNN+PI | 0.878(0.066) | 0.481(0.154) | 0.649(0.163) |
| GAIN+PI | 0.76(0.102) | 0.437(0.126) | 0.524(0.158) |
| HV | 0.865(0.077) | 0.454(0.155) | 0.606(0.172) |

Table 18: Table representing the average accuracy and the standard deviation obtained from the imputation of MAR values. The missing rate is 33%. MK corresponds to the MK model and TM to the threshold model. Sample size of each scenario is 100 for each imputation method. The data contained some strongly correlated traits ($\rho = 0.8$).

| Methods | MK | MK, $\kappa = 0$ | MK, $\lambda = 0.0001$ | TM | TM, $\kappa = 0$ | TM, $\lambda = 0.0001$ |
|---|---|---|---|---|---|---|
| PI | 0.783(0.192) | 0.388(0.25) | 0.627(0.23) | 0.555(0.225) | 0.545(0.241) | 0.445(0.275) |
| MICE | 0.727(0.197) | 0.859(0.127) | 0.751(0.161) | 0.82(0.143) | 0.823(0.136) | 0.811(0.157) |
| MissForest | 0.931(0.094) | 0.908(0.115) | 0.92(0.133) | 0.903(0.118) | 0.919(0.1) | 0.927(0.095) |
| KNN | 0.889(0.129) | 0.865(0.13) | 0.871(0.171) | 0.878(0.145) | 0.892(0.111) | 0.878(0.133) |
| GAIN | 0.676(0.225) | 0.595(0.229) | 0.646(0.221) | 0.602(0.217) | 0.612(0.24) | 0.534(0.234) |
| MICE+P | 0.573(0.244) | 0.76(0.174) | 0.583(0.202) | 0.695(0.188) | 0.685(0.204) | 0.676(0.207) |
| MissForest+P | 0.843(0.165) | 0.739(0.24) | 0.793(0.212) | 0.754(0.201) | 0.789(0.186) | 0.746(0.226) |
| KNN+P | 0.861(0.143) | 0.706(0.197) | 0.797(0.2) | 0.784(0.168) | 0.799(0.168) | 0.718(0.218) |
| GAIN+P | 0.695(0.227) | 0.536(0.223) | 0.64(0.224) | 0.598(0.23) | 0.609(0.233) | 0.582(0.225) |
| MICE+PI | 0.701(0.21) | 0.728(0.143) | 0.603(0.165) | 0.732(0.154) | 0.711(0.14) | 0.671(0.174) |
| MissForest+PI | 0.822(0.163) | 0.633(0.312) | 0.701(0.214) | 0.644(0.237) | 0.642(0.247) | 0.614(0.302) |
| KNN+PI | 0.88(0.143) | 0.773(0.22) | 0.81(0.209) | 0.8(0.18) | 0.803(0.185) | 0.774(0.225) |
| GAIN+PI | 0.683(0.218) | 0.552(0.22) | 0.636(0.229) | 0.596(0.223) | 0.611(0.215) | 0.568(0.246) |
| HV | 0.935(0.087) | 0.893(0.117) | 0.897(0.142) | 0.89(0.119) | 0.89(0.107) | 0.893(0.125) |

Table 19: Tables representing the average accuracy and the standard deviation obtained from the imputation of MAR values. The missing rate is 33%. MK corresponds to the MK model. Sample size of each scenario is 100 for each imputation method. **(a)** Data contained some strongly correlated traits ($\rho = 0.8$). **(b)** Data composed exclusively of independent traits.

(a) Strong correlation ($\rho = 0.8$)

| Methods | MK | MK, $\kappa = 0$ | MK, $\lambda = 0.0001$ |
|---|---|---|---|
| PI | 0.783(0.192) | 0.388(0.25) | 0.627(0.23) |
| MICE | 0.727(0.197) | 0.859(0.127) | 0.751(0.161) |
| MissForest | 0.931(0.094) | 0.908(0.115) | 0.92(0.133) |
| KNN | 0.889(0.129) | 0.865(0.13) | 0.871(0.171) |
| GAIN | 0.676(0.225) | 0.595(0.229) | 0.646(0.221) |
| MICE+P | 0.573(0.244) | 0.76(0.174) | 0.583(0.202) |
| MissForest+P | 0.843(0.165) | 0.739(0.24) | 0.793(0.212) |
| KNN+P | 0.861(0.143) | 0.706(0.197) | 0.797(0.2) |
| GAIN+P | 0.695(0.227) | 0.536(0.223) | 0.64(0.224) |
| MICE+PI | 0.701(0.21) | 0.728(0.143) | 0.603(0.165) |
| MissForest+PI | 0.822(0.163) | 0.633(0.312) | 0.701(0.214) |
| KNN+PI | 0.88(0.143) | 0.773(0.22) | 0.81(0.209) |
| GAIN+PI | 0.683(0.218) | 0.552(0.22) | 0.636(0.229) |
| HV | 0.935(0.087) | 0.893(0.117) | 0.897(0.142) |

(b) Low correlation

| Methods | MK | MK, $\kappa = 0$ | MK, $\lambda = 0.0001$ |
|---|---|---|---|
| PI | 0.815(0.163) | 0.482(0.188) | 0.673(0.176) |
| MICE | 0.386(0.197) | 0.336(0.1) | 0.358(0.143) |
| MissForest | 0.715(0.2) | 0.441(0.155) | 0.61(0.19) |
| KNN | 0.691(0.194) | 0.433(0.155) | 0.608(0.18) |
| GAIN | 0.603(0.237) | 0.402(0.16) | 0.501(0.233) |
| MICE+P | 0.344(0.186) | 0.302(0.111) | 0.237(0.094) |
| MissForest+P | 0.772(0.166) | 0.462(0.154) | 0.641(0.171) |
| KNN+P | 0.735(0.176) | 0.433(0.144) | 0.617(0.177) |
| GAIN+P | 0.637(0.228) | 0.402(0.153) | 0.531(0.217) |
| MICE+PI | 0.469(0.197) | 0.285(0.096) | 0.244(0.086) |
| MissForest+PI | 0.808(0.167) | 0.48(0.189) | 0.686(0.15) |
| KNN+PI | 0.798(0.167) | 0.469(0.178) | 0.677(0.16) |
| GAIN+PI | 0.619(0.233) | 0.423(0.18) | 0.548(0.21) |
| HV | 0.773(0.164) | 0.445(0.166) | 0.619(0.187) |

Table 20: Table representing the average accuracy and the standard deviation obtained from the imputation of MNAR values. The missing rate is 33%. MK corresponds to the MK model and TM to the threshold model. Sample size of each scenario is 100 for each imputation method. The data contained some strongly correlated traits ($\rho = 0.8$).

| Methods | MK | MK, $\kappa = 0$ | MK, $\lambda = 0.0001$ | TM | TM, $\kappa = 0$ | TM, $\lambda = 0.0001$ |
|---|---|---|---|---|---|---|
| PI | 0.726(0.266) | 0.215(0.272) | 0.555(0.364) | 0.474(0.289) | 0.422(0.304) | 0.26(0.367) |
| MICE | 0.693(0.129) | 0.608(0.136) | 0.642(0.142) | 0.679(0.151) | 0.663(0.124) | 0.633(0.152) |
| MissForest | 0.739(0.248) | 0.56(0.268) | 0.682(0.288) | 0.624(0.291) | 0.631(0.275) | 0.596(0.285) |
| KNN | 0.749(0.266) | 0.602(0.271) | 0.695(0.294) | 0.673(0.286) | 0.692(0.279) | 0.659(0.294) |
| GAIN | 0.625(0.243) | 0.352(0.228) | 0.516(0.271) | 0.491(0.247) | 0.459(0.235) | 0.384(0.249) |
| MICE+P | 0.649(0.118) | 0.596(0.102) | 0.486(0.108) | 0.64(0.111) | 0.603(0.111) | 0.549(0.106) |
| MissForest+P | 0.711(0.27) | 0.329(0.256) | 0.569(0.293) | 0.512(0.289) | 0.489(0.289) | 0.352(0.313) |
| KNN+P | 0.749(0.27) | 0.509(0.258) | 0.663(0.303) | 0.626(0.289) | 0.629(0.274) | 0.549(0.302) |
| GAIN+P | 0.657(0.234) | 0.378(0.229) | 0.508(0.251) | 0.491(0.269) | 0.473(0.263) | 0.39(0.268) |
| MICE+PI | 0.775(0.123) | 0.608(0.117) | 0.631(0.12) | 0.672(0.121) | 0.633(0.14) | 0.573(0.126) |
| MissForest+PI | 0.743(0.272) | 0.449(0.314) | 0.666(0.335) | 0.561(0.293) | 0.531(0.301) | 0.445(0.374) |
| KNN+PI | 0.754(0.274) | 0.564(0.279) | 0.692(0.322) | 0.641(0.307) | 0.65(0.29) | 0.604(0.312) |
| GAIN+PI | 0.651(0.216) | 0.345(0.211) | 0.51(0.248) | 0.466(0.245) | 0.468(0.245) | 0.382(0.248) |
| HV | 0.772(0.242) | 0.577(0.259) | 0.695(0.292) | 0.65(0.284) | 0.651(0.269) | 0.612(0.278) |

Table 21: Tables representing the average accuracy and the standard deviation obtained from the imputation of MNAR values. The missing rate is 33%. MK corresponds to the MK model. Sample size of each scenario is 100 for each imputation method. **(a)** Data contained some strongly correlated traits ($\rho = 0.8$). **(b)** Data composed exclusively of independent traits.

(a) Strong correlation ($\rho = 0.8$)

| Methods | MK | MK, $\kappa = 0$ | MK, $\lambda = 0.0001$ |
|---|---|---|---|
| PI | 0.726(0.266) | 0.215(0.272) | 0.555(0.364) |
| MICE | 0.693(0.129) | 0.608(0.136) | 0.642(0.142) |
| MissForest | 0.739(0.248) | 0.56(0.268) | 0.682(0.288) |
| KNN | 0.749(0.266) | 0.602(0.271) | 0.695(0.294) |
| GAIN | 0.625(0.243) | 0.352(0.228) | 0.516(0.271) |
| MICE+P | 0.649(0.118) | 0.596(0.102) | 0.486(0.108) |
| MissForest+P | 0.711(0.27) | 0.329(0.256) | 0.569(0.293) |
| KNN+P | 0.749(0.27) | 0.509(0.258) | 0.663(0.303) |
| GAIN+P | 0.657(0.234) | 0.378(0.229) | 0.508(0.251) |
| MICE+PI | 0.775(0.123) | 0.608(0.117) | 0.631(0.12) |
| MissForest+PI | 0.743(0.272) | 0.449(0.314) | 0.666(0.335) |
| KNN+PI | 0.754(0.274) | 0.564(0.279) | 0.692(0.322) |
| GAIN+PI | 0.651(0.216) | 0.345(0.211) | 0.51(0.248) |
| HV | 0.772(0.242) | 0.577(0.259) | 0.695(0.292) |

(b) Low correlation

| Methods | MK | MK, $\kappa = 0$ | MK, $\lambda = 0.0001$ |
|---|---|---|---|
| PI | 0.705(0.268) | 0.226(0.265) | 0.511(0.349) |
| MICE | 0.503(0.178) | 0.267(0.144) | 0.399(0.199) |
| MissForest | 0.566(0.284) | 0.224(0.21) | 0.462(0.28) |
| KNN | 0.578(0.275) | 0.228(0.211) | 0.482(0.288) |
| GAIN | 0.574(0.234) | 0.261(0.179) | 0.424(0.22) |
| MICE+P | 0.611(0.128) | 0.352(0.11) | 0.386(0.127) |
| MissForest+P | 0.664(0.276) | 0.271(0.221) | 0.447(0.272) |
| KNN+P | 0.617(0.286) | 0.241(0.222) | 0.482(0.289) |
| GAIN+P | 0.586(0.241) | 0.258(0.193) | 0.441(0.256) |
| MICE+PI | 0.642(0.166) | 0.36(0.131) | 0.364(0.131) |
| MissForest+PI | 0.66(0.299) | 0.225(0.243) | 0.531(0.324) |
| KNN+PI | 0.634(0.299) | 0.228(0.23) | 0.508(0.311) |
| GAIN+PI | 0.581(0.216) | 0.253(0.167) | 0.419(0.239) |
| HV | 0.623(0.284) | 0.224(0.214) | 0.47(0.288) |

Table 22: Table representing the average accuracy and the standard deviation obtained from the imputation of phyloNA values. The missing rate is 33%. MK corresponds to the MK model and TM to the threshold model. Sample size of each scenario is 100 for each imputation method. The data contained some strongly correlated traits ($\rho = 0.8$).

| Methods | MK | MK, $\kappa = 0$ | MK, $\lambda = 0.0001$ | TM | TM, $\kappa = 0$ | TM, $\lambda = 0.0001$ |
|---|---|---|---|---|---|---|
| PI | 0.841(0.191) | 0.496(0.163) | 0.658(0.166) | 0.737(0.143) | 0.652(0.201) | 0.565(0.18) |
| MICE | 0.371(0.104) | 0.34(0.089) | 0.368(0.094) | 0.355(0.098) | 0.35(0.1) | 0.356(0.083) |
| MissForest | 0.625(0.284) | 0.495(0.164) | 0.703(0.121) | 0.542(0.239) | 0.569(0.218) | 0.577(0.176) |
| KNN | 0.451(0.328) | 0.368(0.181) | 0.44(0.295) | 0.45(0.253) | 0.439(0.248) | 0.438(0.228) |
| GAIN | 0.451(0.237) | 0.358(0.138) | 0.431(0.206) | 0.391(0.171) | 0.395(0.21) | 0.384(0.147) |
| MICE+P | 0.375(0.19) | 0.305(0.095) | 0.231(0.089) | 0.38(0.116) | 0.345(0.141) | 0.26(0.099) |
| MissForest+P | 0.83(0.148) | 0.515(0.153) | 0.663(0.136) | 0.695(0.15) | 0.656(0.165) | 0.538(0.189) |
| KNN+P | 0.792(0.184) | 0.438(0.152) | 0.564(0.193) | 0.659(0.159) | 0.616(0.165) | 0.482(0.185) |
| GAIN+P | 0.521(0.264) | 0.386(0.13) | 0.465(0.203) | 0.444(0.204) | 0.408(0.175) | 0.427(0.181) |
| MICE+PI | 0.679(0.221) | 0.334(0.113) | 0.28(0.122) | 0.597(0.149) | 0.528(0.165) | 0.305(0.109) |
| MissForest+PI | 0.848(0.146) | 0.504(0.155) | 0.688(0.129) | 0.724(0.154) | 0.67(0.187) | 0.563(0.182) |
| KNN+PI | 0.852(0.144) | 0.503(0.16) | 0.678(0.147) | 0.72(0.152) | 0.662(0.192) | 0.563(0.181) |
| GAIN+PI | 0.452(0.255) | 0.368(0.138) | 0.399(0.192) | 0.417(0.188) | 0.375(0.175) | 0.398(0.16) |
| HV | 0.839(0.151) | 0.491(0.169) | 0.68(0.146) | 0.704(0.166) | 0.65(0.194) | 0.566(0.178) |

Table 23: Tables representing the average accuracy and the standard deviation obtained from the imputation of phyloNA values. The missing rate is 33%. MK corresponds to the MK model. Sample size of each scenario is 100 for each imputation method. **(a)** Data contained some strongly correlated traits ($\rho = 0.8$). **(b)** Data composed exclusively of independent traits.

(a) Strong correlation ($\rho = 0.8$)

| Methods | MK | MK, $\kappa = 0$ | MK, $\lambda = 0.0001$ |
|---|---|---|---|
| PI | 0.841(0.191) | 0.496(0.163) | 0.658(0.166) |
| MICE | 0.371(0.104) | 0.34(0.089) | 0.368(0.094) |
| MissForest | 0.625(0.284) | 0.495(0.164) | 0.703(0.121) |
| KNN | 0.451(0.328) | 0.368(0.181) | 0.44(0.295) |
| GAIN | 0.451(0.237) | 0.358(0.138) | 0.431(0.206) |
| MICE+P | 0.375(0.19) | 0.305(0.095) | 0.231(0.089) |
| MissForest+P | 0.83(0.148) | 0.515(0.153) | 0.663(0.136) |
| KNN+P | 0.792(0.184) | 0.438(0.152) | 0.564(0.193) |
| GAIN+P | 0.521(0.264) | 0.386(0.13) | 0.465(0.203) |
| MICE+PI | 0.679(0.221) | 0.334(0.113) | 0.28(0.122) |
| MissForest+PI | 0.848(0.146) | 0.504(0.155) | 0.688(0.129) |
| KNN+PI | 0.852(0.144) | 0.503(0.16) | 0.678(0.147) |
| GAIN+PI | 0.452(0.255) | 0.368(0.138) | 0.399(0.192) |
| HV | 0.839(0.151) | 0.491(0.169) | 0.68(0.146) |

(b) Low correlation

| Methods | MK | MK, $\kappa = 0$ | MK, $\lambda = 0.0001$ |
|---|---|---|---|
| PI | 0.832(0.22) | 0.488(0.173) | 0.679(0.174) |
| MICE | 0.424(0.177) | 0.373(0.134) | 0.481(0.15) |
| MissForest | 0.636(0.286) | 0.442(0.19) | 0.63(0.234) |
| KNN | 0.59(0.309) | 0.432(0.189) | 0.569(0.262) |
| GAIN | 0.484(0.242) | 0.378(0.134) | 0.437(0.23) |
| MICE+P | 0.386(0.197) | 0.344(0.093) | 0.294(0.111) |
| MissForest+P | 0.822(0.172) | 0.486(0.158) | 0.6(0.178) |
| KNN+P | 0.783(0.212) | 0.46(0.157) | 0.614(0.178) |
| GAIN+P | 0.489(0.248) | 0.367(0.153) | 0.472(0.226) |
| MICE+PI | 0.647(0.242) | 0.35(0.117) | 0.307(0.116) |
| MissForest+PI | 0.853(0.147) | 0.495(0.165) | 0.686(0.157) |
| KNN+PI | 0.843(0.176) | 0.497(0.163) | 0.667(0.17) |
| GAIN+PI | 0.526(0.258) | 0.385(0.173) | 0.462(0.211) |
| HV | 0.82(0.194) | 0.46(0.168) | 0.632(0.209) |

Table 24: Table representing the average accuracy and the standard deviation obtained from the imputation of MCAR values. The missing rate is 50%. MK corresponds to the MK model and TM to the threshold model. Sample size of each scenario is 100 for each imputation method. The data contained some strongly correlated traits ($\rho = 0.8$).

| Methods | MK | MK, $\kappa = 0$ | MK, $\lambda = 0.0001$ | TM | TM, $\kappa = 0$ | TM, $\lambda = 0.0001$ |
|---|---|---|---|---|---|---|
| PI | 0.899(0.059) | 0.501(0.162) | 0.681(0.144) | 0.745(0.122) | 0.67(0.152) | 0.539(0.186) |
| MICE | 0.666(0.11) | 0.701(0.064) | 0.626(0.089) | 0.697(0.088) | 0.704(0.094) | 0.688(0.089) |
| MissForest | 0.9(0.06) | 0.839(0.065) | 0.879(0.064) | 0.867(0.068) | 0.865(0.074) | 0.863(0.073) |
| KNN | 0.75(0.131) | 0.604(0.113) | 0.714(0.111) | 0.671(0.12) | 0.672(0.124) | 0.662(0.12) |
| GAIN | 0.728(0.109) | 0.494(0.123) | 0.6(0.142) | 0.641(0.125) | 0.611(0.122) | 0.514(0.151) |
| MICE+P | 0.663(0.12) | 0.596(0.087) | 0.497(0.097) | 0.661(0.106) | 0.635(0.094) | 0.552(0.098) |
| MissForest+P | 0.914(0.054) | 0.635(0.122) | 0.684(0.117) | 0.825(0.091) | 0.792(0.094) | 0.639(0.133) |
| KNN+P | 0.839(0.08) | 0.588(0.117) | 0.695(0.124) | 0.739(0.097) | 0.726(0.103) | 0.631(0.128) |
| GAIN+P | 0.756(0.09) | 0.51(0.122) | 0.584(0.147) | 0.676(0.113) | 0.618(0.131) | 0.528(0.156) |
| MICE+PI | 0.822(0.113) | 0.624(0.09) | 0.567(0.089) | 0.76(0.094) | 0.716(0.098) | 0.584(0.094) |
| MissForest+PI | 0.933(0.044) | 0.775(0.082) | 0.774(0.095) | 0.857(0.068) | 0.831(0.08) | 0.763(0.107) |
| KNN+PI | 0.905(0.053) | 0.704(0.084) | 0.766(0.09) | 0.829(0.072) | 0.795(0.083) | 0.727(0.097) |
| GAIN+PI | 0.719(0.118) | 0.496(0.12) | 0.59(0.135) | 0.637(0.118) | 0.62(0.128) | 0.503(0.146) |
| HV | 0.941(0.038) | 0.806(0.071) | 0.854(0.069) | 0.875(0.058) | 0.854(0.061) | 0.816(0.08) |

Table 25: Table representing the average accuracy and the standard deviation obtained from the imputation of MAR values. The missing rate is 50%. MK corresponds to the MK model and TM to the threshold model. Sample size of each scenario is 100 for each imputation method. The data contained some strongly correlated traits ($\rho = 0.8$).

| Methods | MK | MK, $\kappa = 0$ | MK, $\lambda = 0.0001$ | TM | TM, $\kappa = 0$ | TM, $\lambda = 0.0001$ |
|---|---|---|---|---|---|---|
| PI | 0.724(0.215) | 0.381(0.237) | 0.607(0.248) | 0.545(0.231) | 0.494(0.232) | 0.432(0.286) |
| MICE | 0.67(0.227) | 0.822(0.13) | 0.698(0.16) | 0.786(0.127) | 0.779(0.145) | 0.762(0.161) |
| MissForest | 0.887(0.133) | 0.882(0.125) | 0.926(0.095) | 0.894(0.125) | 0.885(0.121) | 0.892(0.138) |
| KNN | 0.835(0.159) | 0.823(0.175) | 0.877(0.154) | 0.852(0.148) | 0.827(0.151) | 0.829(0.193) |
| GAIN | 0.699(0.216) | 0.532(0.233) | 0.645(0.237) | 0.6(0.215) | 0.595(0.224) | 0.546(0.248) |
| MICE+P | 0.561(0.22) | 0.769(0.155) | 0.587(0.169) | 0.684(0.174) | 0.677(0.18) | 0.673(0.186) |
| MissForest+P | 0.762(0.203) | 0.669(0.242) | 0.792(0.178) | 0.727(0.227) | 0.696(0.229) | 0.693(0.27) |
| KNN+P | 0.807(0.17) | 0.698(0.229) | 0.813(0.17) | 0.784(0.18) | 0.752(0.185) | 0.705(0.256) |
| GAIN+P | 0.642(0.225) | 0.479(0.198) | 0.595(0.233) | 0.581(0.214) | 0.554(0.205) | 0.52(0.249) |
| MICE+PI | 0.624(0.197) | 0.703(0.136) | 0.556(0.168) | 0.679(0.127) | 0.664(0.141) | 0.632(0.165) |
| MissForest+PI | 0.761(0.209) | 0.577(0.314) | 0.737(0.206) | 0.685(0.231) | 0.619(0.248) | 0.573(0.33) |
| KNN+PI | 0.798(0.193) | 0.717(0.235) | 0.821(0.183) | 0.782(0.195) | 0.74(0.208) | 0.731(0.264) |
| GAIN+PI | 0.621(0.227) | 0.546(0.225) | 0.627(0.222) | 0.608(0.237) | 0.582(0.22) | 0.541(0.232) |
| HV | 0.877(0.143) | 0.849(0.143) | 0.912(0.104) | 0.877(0.127) | 0.849(0.135) | 0.849(0.17) |

Table 26: Table representing the average accuracy and the standard deviation obtained from the imputation of MNAR values. The missing rate is 50%. MK corresponds to the MK model and TM to the threshold model. Sample size of each scenario is 100 for each imputation method. The data contained some strongly correlated traits ($\rho = 0.8$).

| Methods | MK | MK, $\kappa = 0$ | MK, $\lambda = 0.0001$ | TM | TM, $\kappa = 0$ | TM, $\lambda = 0.0001$ |
|---|---|---|---|---|---|---|
| PI | 0.624(0.31) | 0.17(0.237) | 0.452(0.344) | 0.416(0.291) | 0.37(0.286) | 0.281(0.305) |
| MICE | 0.519(0.155) | 0.436(0.11) | 0.531(0.117) | 0.494(0.127) | 0.465(0.138) | 0.447(0.133) |
| MissForest | 0.578(0.327) | 0.348(0.244) | 0.635(0.256) | 0.442(0.302) | 0.418(0.303) | 0.404(0.287) |
| KNN | 0.627(0.3) | 0.425(0.256) | 0.669(0.244) | 0.521(0.289) | 0.502(0.279) | 0.469(0.268) |
| GAIN | 0.523(0.264) | 0.273(0.215) | 0.495(0.241) | 0.387(0.256) | 0.365(0.24) | 0.323(0.241) |
| MICE+P | 0.541(0.101) | 0.447(0.069) | 0.437(0.074) | 0.54(0.095) | 0.507(0.102) | 0.424(0.08) |
| MissForest+P | 0.602(0.32) | 0.237(0.217) | 0.506(0.274) | 0.409(0.291) | 0.382(0.288) | 0.306(0.259) |
| KNN+P | 0.63(0.304) | 0.365(0.242) | 0.622(0.244) | 0.5(0.279) | 0.469(0.28) | 0.414(0.255) |
| GAIN+P | 0.539(0.265) | 0.275(0.211) | 0.502(0.243) | 0.411(0.24) | 0.372(0.242) | 0.332(0.264) |
| MICE+PI | 0.652(0.13) | 0.49(0.11) | 0.518(0.123) | 0.563(0.132) | 0.536(0.142) | 0.484(0.113) |
| MissForest+PI | 0.633(0.328) | 0.262(0.27) | 0.624(0.301) | 0.448(0.302) | 0.407(0.315) | 0.349(0.317) |
| KNN+PI | 0.628(0.327) | 0.375(0.262) | 0.662(0.265) | 0.493(0.291) | 0.464(0.297) | 0.419(0.284) |
| GAIN+PI | 0.52(0.26) | 0.274(0.203) | 0.448(0.244) | 0.397(0.25) | 0.374(0.234) | 0.345(0.259) |
| HV | 0.621(0.319) | 0.365(0.243) | 0.644(0.251) | 0.479(0.291) | 0.445(0.291) | 0.412(0.278) |

Table 27: Table representing the average accuracy and the standard deviation obtained from the imputation of phyloNA values. The missing rate is 50%. MK corresponds to the MK model and TM to the threshold model. Sample size of each scenario is 100 for each imputation method. The data contained some strongly correlated traits ($\rho = 0.8$).

| Methods | MK | MK, $\kappa = 0$ | MK, $\lambda = 0.0001$ | TM | TM, $\kappa = 0$ | TM, $\lambda = 0.0001$ |
|---|---|---|---|---|---|---|
| PI | 0.801(0.207) | 0.483(0.159) | 0.665(0.156) | 0.696(0.17) | 0.655(0.187) | 0.542(0.196) |
| MICE | 0.367(0.093) | 0.322(0.077) | 0.345(0.078) | 0.368(0.086) | 0.362(0.077) | 0.339(0.07) |
| MissForest | 0.646(0.235) | 0.483(0.17) | 0.698(0.125) | 0.533(0.22) | 0.546(0.219) | 0.575(0.18) |
| KNN | 0.489(0.299) | 0.377(0.18) | 0.437(0.279) | 0.423(0.236) | 0.432(0.236) | 0.436(0.232) |
| GAIN | 0.459(0.237) | 0.365(0.125) | 0.433(0.205) | 0.401(0.191) | 0.443(0.205) | 0.419(0.195) |
| MICE+P | 0.396(0.146) | 0.319(0.079) | 0.269(0.083) | 0.393(0.108) | 0.369(0.104) | 0.299(0.087) |
| MissForest+P | 0.829(0.113) | 0.493(0.157) | 0.658(0.14) | 0.683(0.159) | 0.66(0.165) | 0.537(0.173) |
| KNN+P | 0.769(0.159) | 0.453(0.135) | 0.541(0.204) | 0.648(0.165) | 0.605(0.174) | 0.481(0.169) |
| GAIN+P | 0.51(0.23) | 0.368(0.131) | 0.43(0.188) | 0.439(0.196) | 0.466(0.199) | 0.419(0.182) |
| MICE+PI | 0.671(0.179) | 0.344(0.096) | 0.305(0.091) | 0.56(0.148) | 0.504(0.15) | 0.319(0.125) |
| MissForest+PI | 0.84(0.11) | 0.5(0.16) | 0.682(0.134) | 0.702(0.165) | 0.66(0.184) | 0.564(0.183) |
| KNN+PI | 0.842(0.11) | 0.495(0.158) | 0.669(0.154) | 0.694(0.164) | 0.658(0.175) | 0.549(0.19) |
| GAIN+PI | 0.47(0.222) | 0.371(0.133) | 0.439(0.222) | 0.403(0.173) | 0.425(0.166) | 0.397(0.17) |
| HV | 0.821(0.128) | 0.488(0.162) | 0.671(0.148) | 0.679(0.165) | 0.641(0.183) | 0.555(0.186) |

#### 4.2 Lines plots

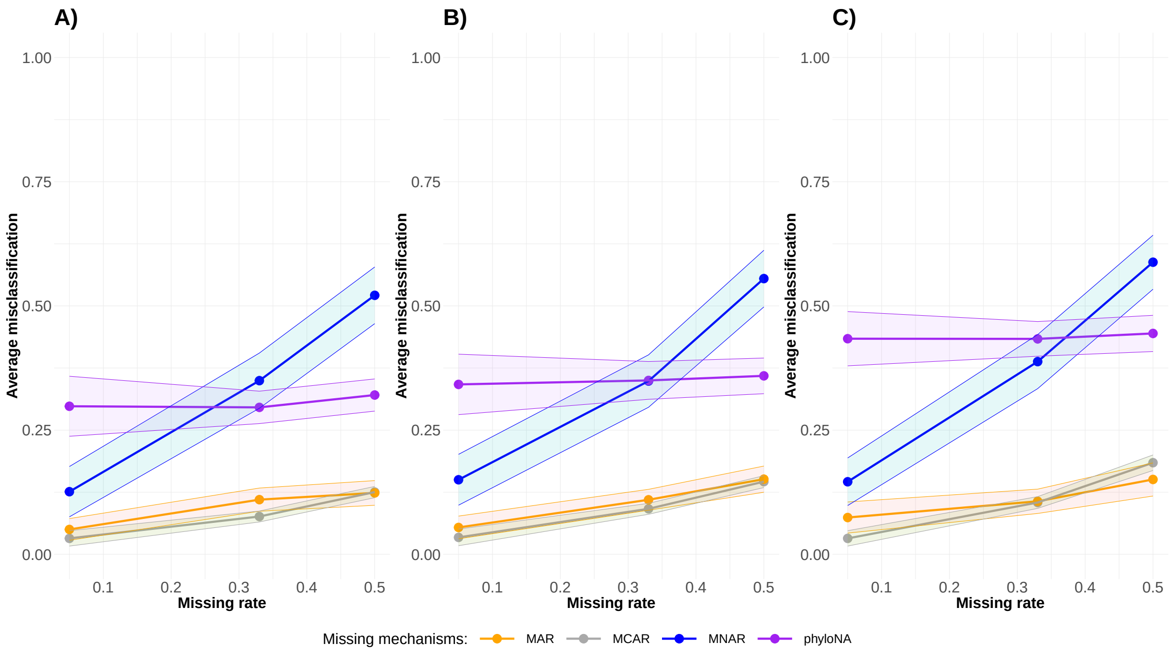

Figure 11: Plots representing the effect of the missing mechanisms and of the missing rate according to the phylogenetic signal. The dots represent the averaged misclassification of the HV approach. The shade represents the 95% confidence interval. **A)** Traits simulated with $\kappa = 1$ and $\lambda = 1$. **B)** Traits simulated with $\kappa = 0$ and $\lambda = 1$. **C)** Traits simulated with $\kappa = 1$ and $\lambda = 0.0001$. The data were simulated through a threshold model and contained strongly correlated traits ($\rho = 0.8$).

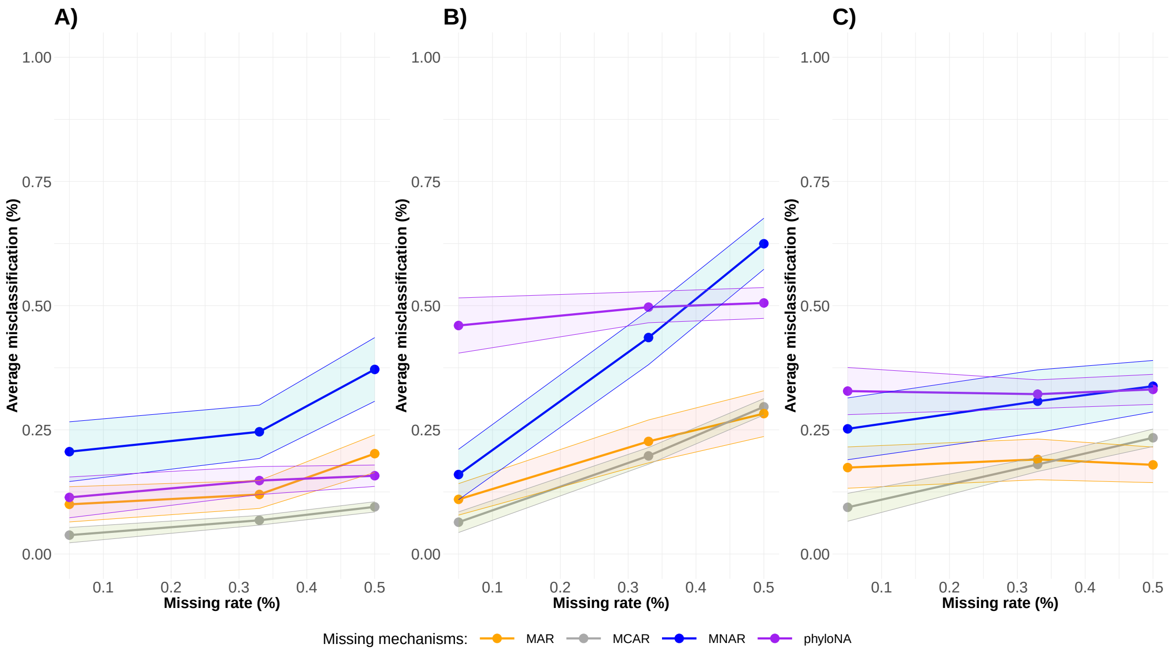

Figure 12: Plots representing the effect of the missing mechanisms and of the missing rate according to the phylogenetic signal. The dots represent the averaged misclassification of the kNN+PI approach. The shade represents the 95% confidence interval. **A)** Traits simulated with $\kappa = 1$ and $\lambda = 1$. **B)** Traits simulated with $\kappa = 0$ and $\lambda = 1$. **C)** Traits simulated with $\kappa = 1$ and $\lambda = 0.0001$. The data were simulated through a MK model and contained strongly correlated traits ($\rho = 0.8$).

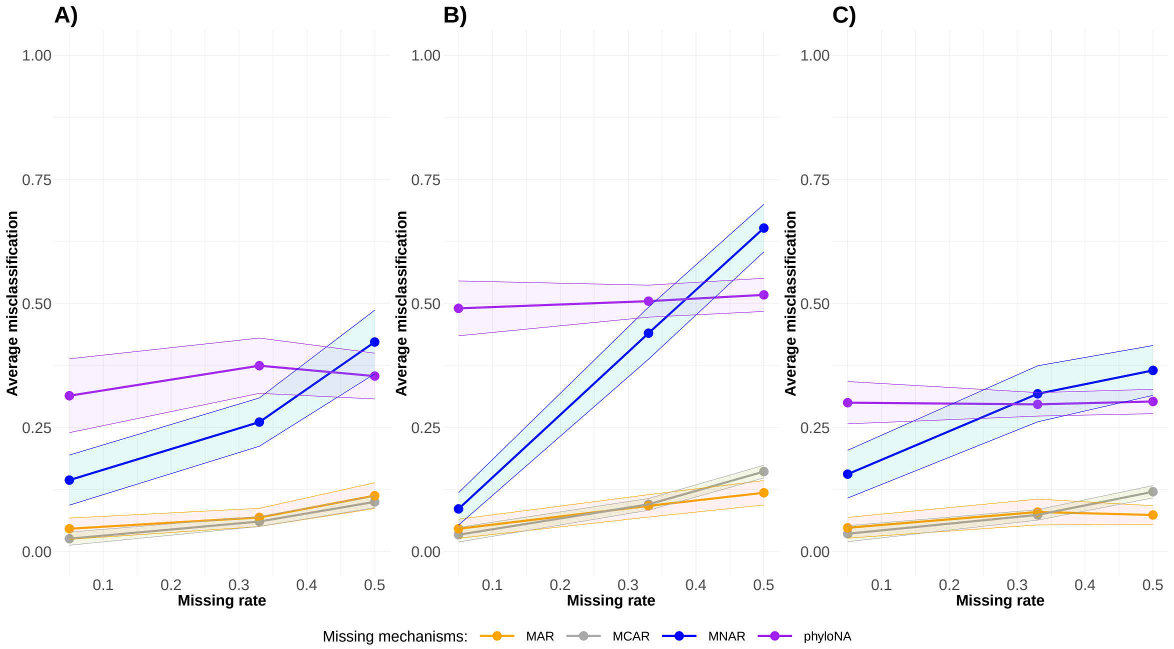

Figure 13: Plots representing the effect of the missing mechanisms and of the missing rate according to the phylogenetic signal. The dots represent the averaged misclassification of the MissForest approach. The shade represents the 95% confidence interval. **A)** Traits simulated with $\kappa = 1$ and $\lambda = 1$. **B)** Traits simulated with $\kappa = 0$ and $\lambda = 1$. **C)** Traits simulated with $\kappa = 1$ and $\lambda = 0.0001$. The data were simulated through a MK model and contained strongly correlated traits ($\rho = 0.8$).

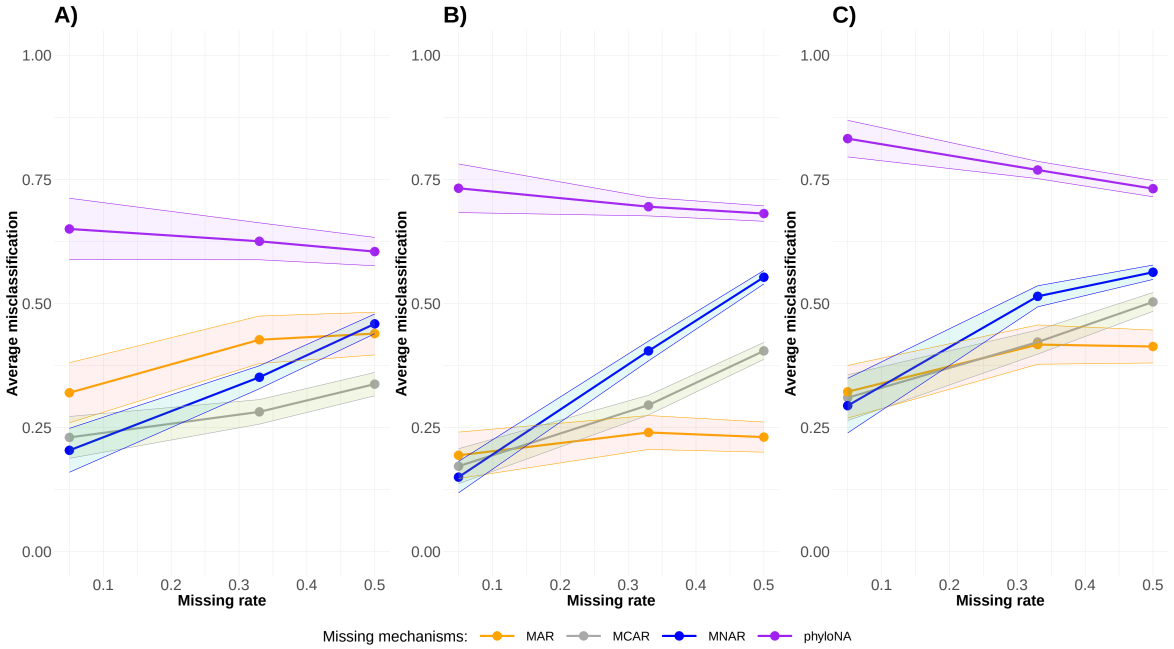

Figure 14: Plots representing the effect of the missing mechanisms and of the missing rate according to the phylogenetic signal. The dots represent the averaged misclassification of the MICE+P approach. The shade represents the 95% confidence interval. **A)** Traits simulated with $\kappa = 1$ and $\lambda = 1$. **B)** Traits simulated with $\kappa = 0$ and $\lambda = 1$. **C)** Traits simulated with $\kappa = 1$ and $\lambda = 0.0001$. The data were simulated through a MK model and contained strongly correlated traits ($\rho = 0.8$).

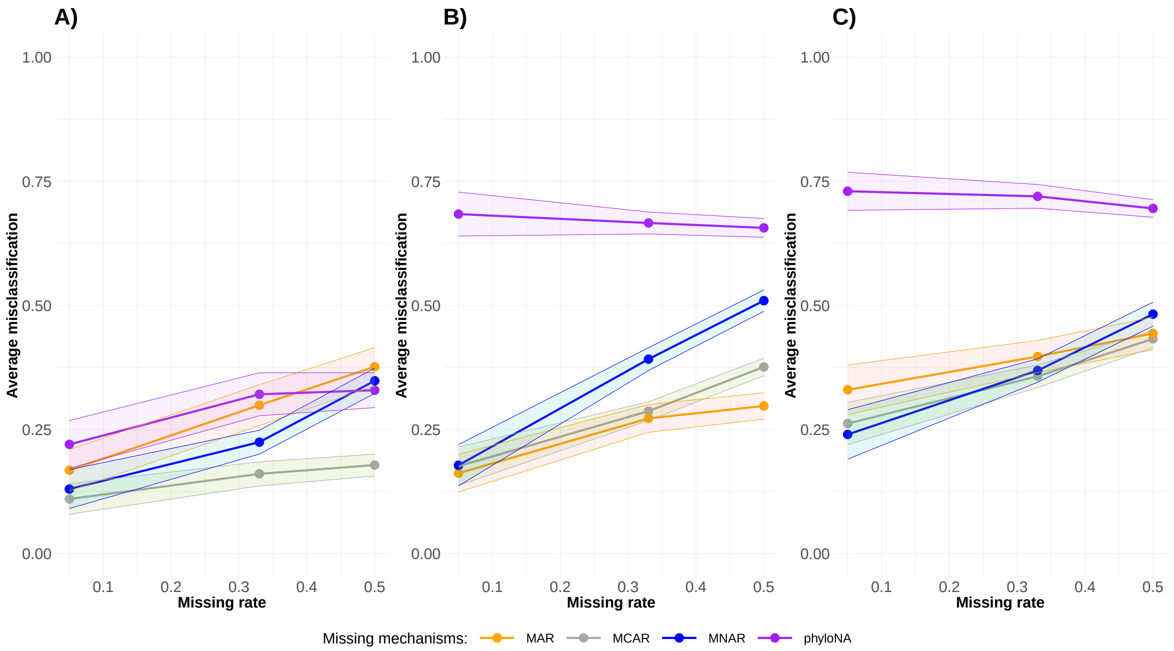

Figure 15: Plots representing the effect of the missing mechanisms and of the missing rate according to the phylogenetic signal. The dots represent the averaged misclassification of the MICE+PI approach. The shade represents the 95% confidence interval. **A)** Traits simulated with $\kappa = 1$ and $\lambda = 1$. **B)** Traits simulated with $\kappa = 0$ and $\lambda = 1$. **C)** Traits simulated with $\kappa = 1$ and $\lambda = 0.0001$. The data were simulated through a MK model and contained strongly correlated traits ($\rho = 0.8$).

#### 4.3 Improvement boxplots

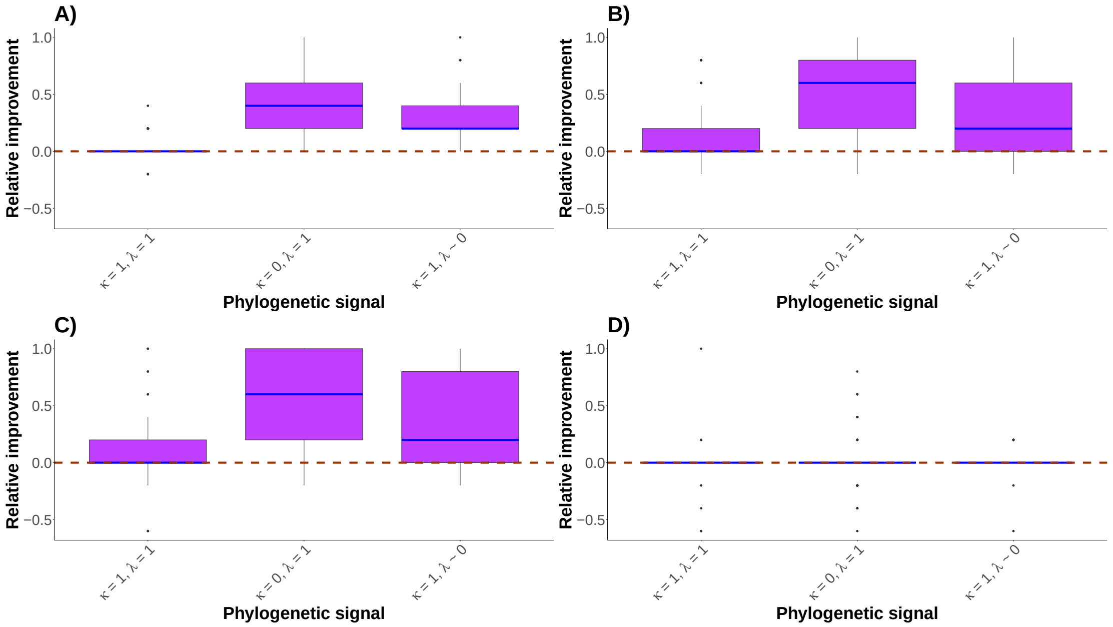

Figure 16: Boxplots representing the performance of the HV method according to different phylogenetic signals compared to PI method for the 4 missing mechanisms (MCAR (**A**), MAR (**B**), MNAR (**C**) and phyloNA (**D**)). Boxplots represent the difference between PI and HV accuracy. Therefore, the dashed line indicates no difference in performance between PI method and HV. The blue line in the boxplots is the median. Data simulated by a MK model and contained strongly correlated traits ($\rho = 0.8$). The missing rate is of 5%.

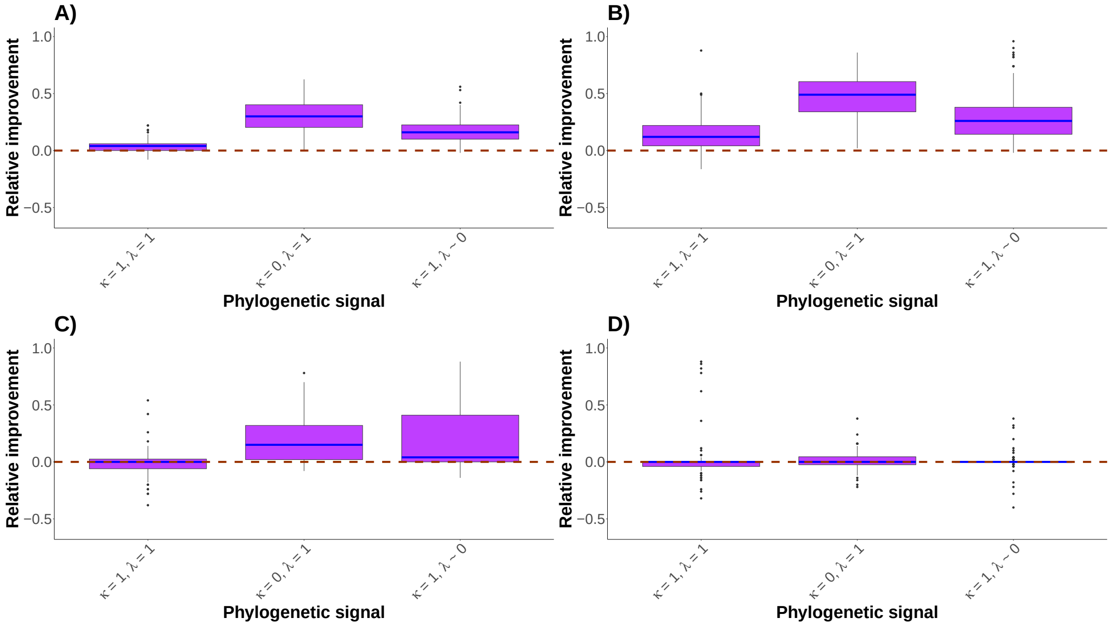

Figure 17: Boxplots representing the performance of the HV method according to different phylogenetic signals compared to PI method for the 4 missing mechanisms (MCAR (**A**), MAR (**B**), MNAR (**C**) and phyloNA (**D**)). Boxplots represent the difference between PI and HV accuracy. Therefore, the dashed line indicates no difference in performance between PI method and HV. The blue line in the boxplots is the median. Data simulated by a MK model and contained strongly correlated traits ($\rho = 0.8$). The missing rate is of 50%.

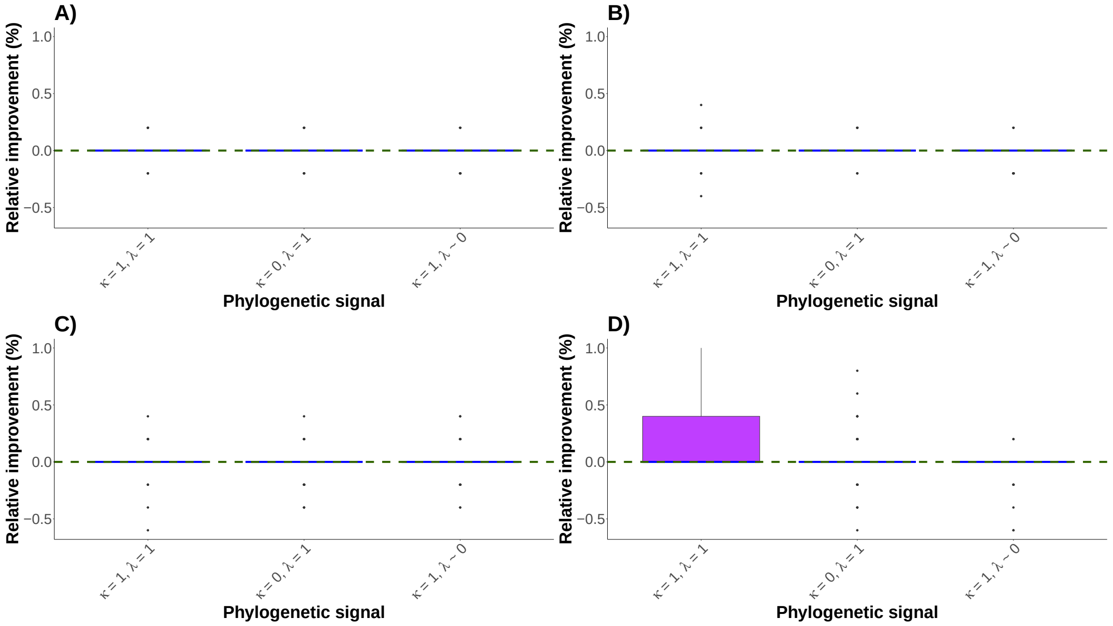

Figure 18: Boxplots representing the performance of the HV method according to different phylogenetic signals compared to MissForest method for the 4 missing mechanisms (MCAR (**A**), MAR (**B**), MNAR (**C**) and phyloNA (**D**)). Boxplots represent the difference between MissForest and HV accuracy. Therefore, the dashed line indicates no difference in performance between MissForest and HV. The blue line in the boxplots is the median. Data simulated by a MK model and contained strongly correlated traits ($\rho = 0.8$). The missing rate is of 5%.

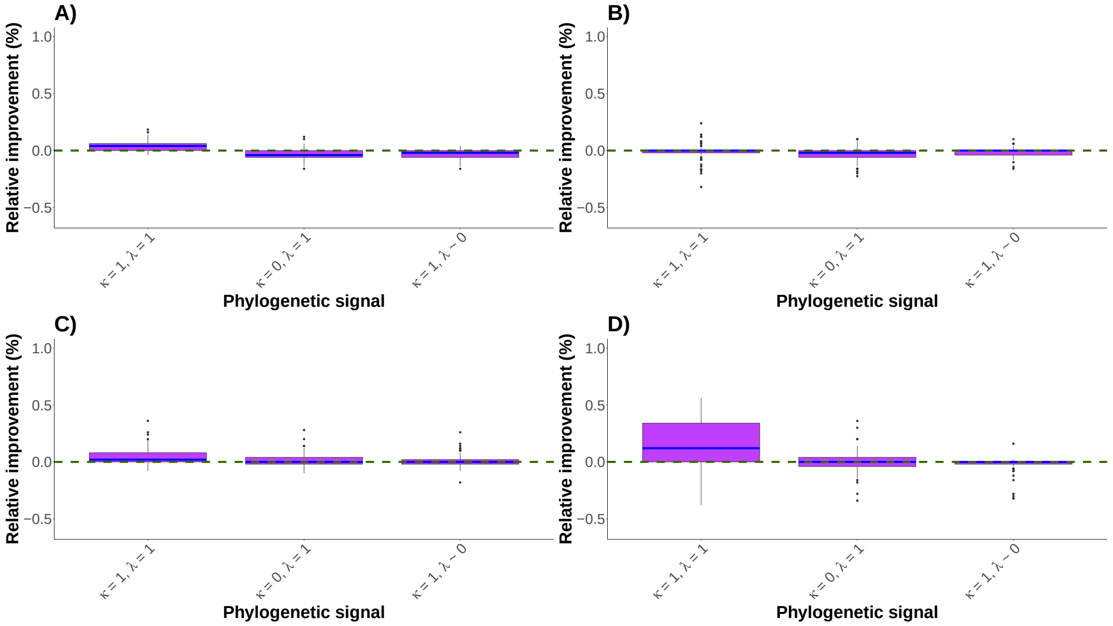

Figure 19: Boxplots representing the performance of the HV method according to different phylogenetic signals compared to MissForest method for the 4 missing mechanisms (MCAR (**A**), MAR (**B**), MNAR (**C**) and phyloNA (**D**)). Boxplots represent the difference between MissForest and HV accuracy. Therefore, the dashed line indicates no difference in performance between MissForest and HV. The blue line in the boxplots is the median. Data simulated by a MK model and contained strongly correlated traits ($\rho = 0.8$). The missing rate is of 50%.

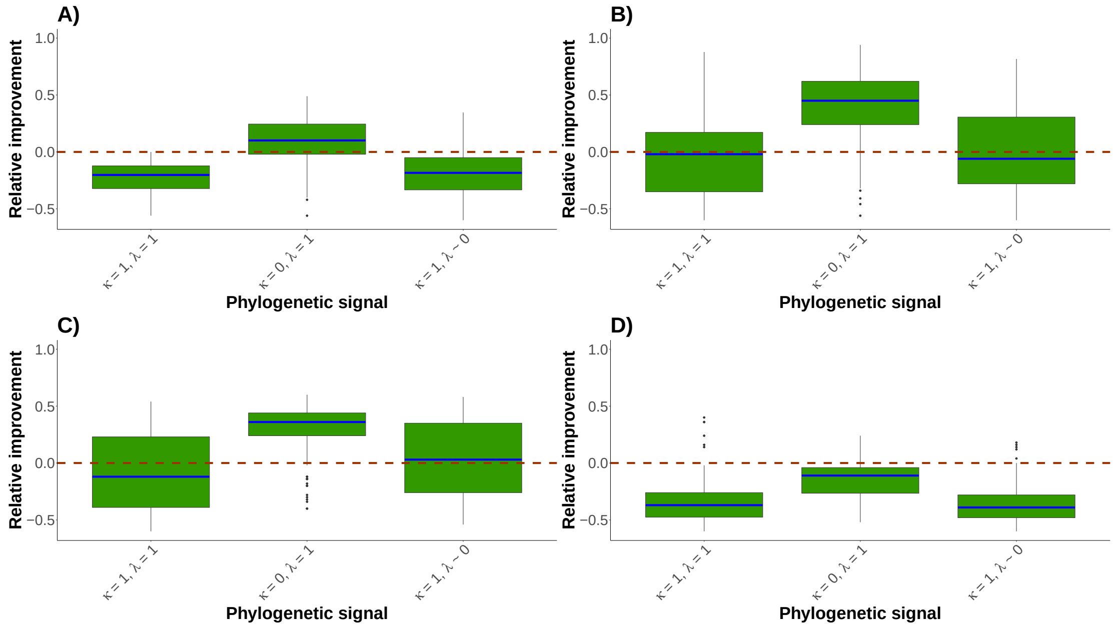

Figure 20: Boxplots representing the performance of the MICE+P method according to different phylogenetic signals compared to PI method for the 4 missing mechanisms (MCAR (**A**), MAR (**B**), MNAR (**C**) and phyloNA (**D**)). Boxplots represent the difference between PI and MICE+P accuracy. Therefore, the dashed line indicates no difference in performance between PI method and MICE+P. The blue line in the boxplots is the median. Data simulated by a MK model and contained strongly correlated traits ($\rho = 0.8$). The missing rate is of 50%.

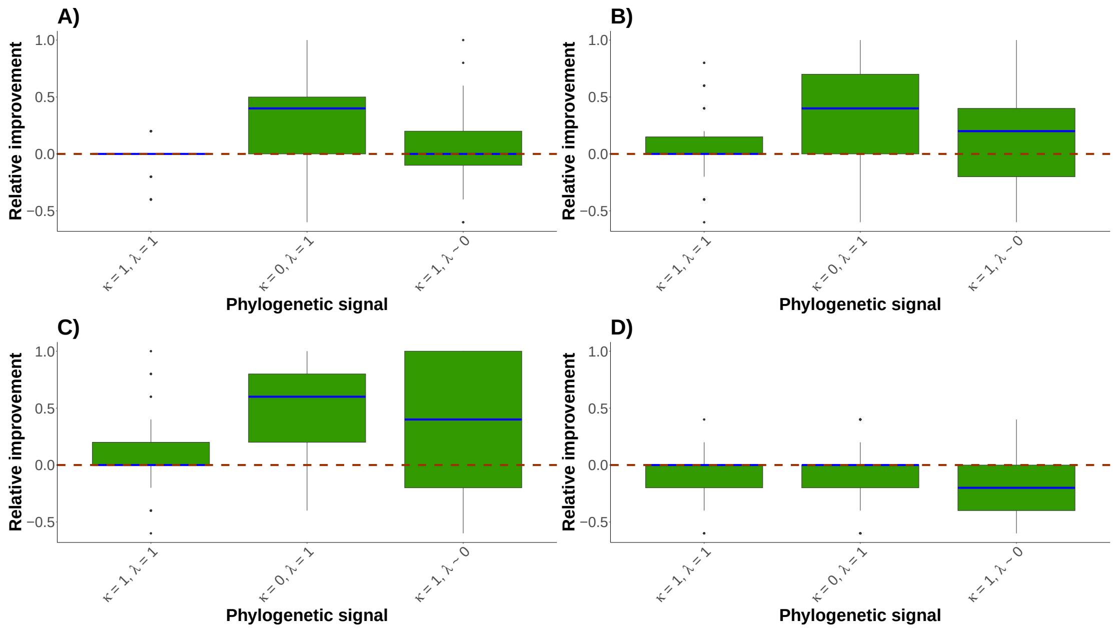

Figure 21: Boxplots representing the performance of the MICE+PI method according to different phylogenetic signals compared to PI method for the 4 missing mechanisms (MCAR (**A**), MAR (**B**), MNAR (**C**) and phyloNA (**D**)). Boxplots represent the difference between PI and MICE+PI accuracy. Therefore, the dashed line indicates no difference in performance between PI method and MICE+PI. The blue line in the boxplots is the median. Data simulated by a MK model and contained strongly correlated traits ($\rho = 0.8$). The missing rate is of 5%.

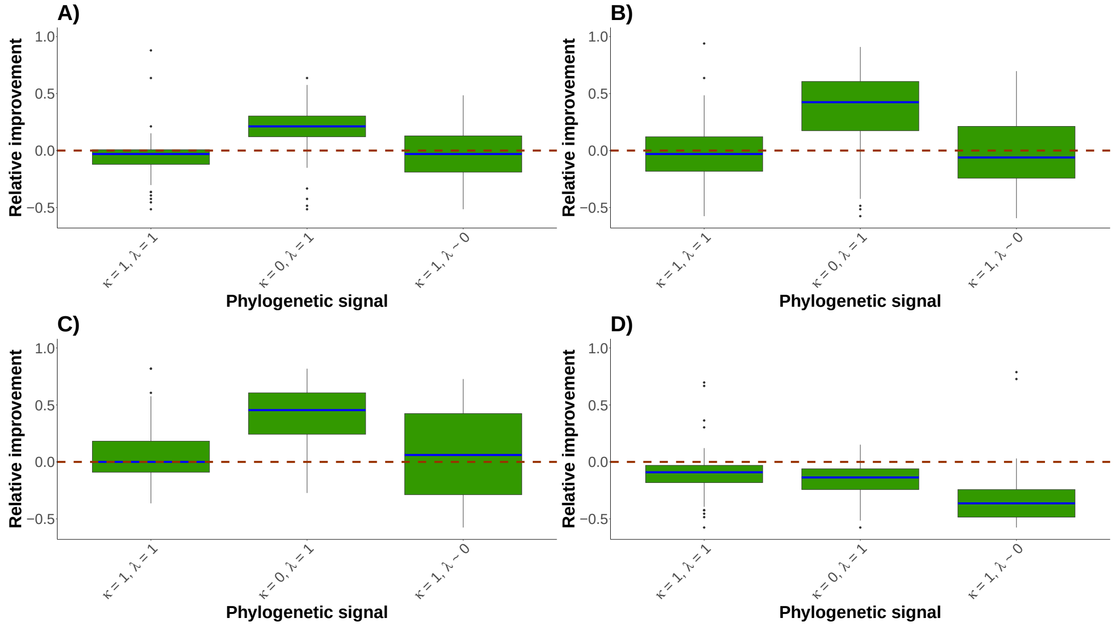

Figure 22: Boxplots representing the performance of the MICE+PI method according to different phylogenetic signals compared to PI method for the 4 missing mechanisms (MCAR (**A**), MAR (**B**), MNAR (**C**) and phyloNA (**D**)). Boxplots represent the difference between PI and MICE+PI accuracy. Therefore, the dashed line indicates no difference in performance between PI method and MICE+PI. The blue line in the boxplots is the median. Data simulated by a MK model and contained strongly correlated traits ($\rho = 0.8$). The missing rate is of 33%.

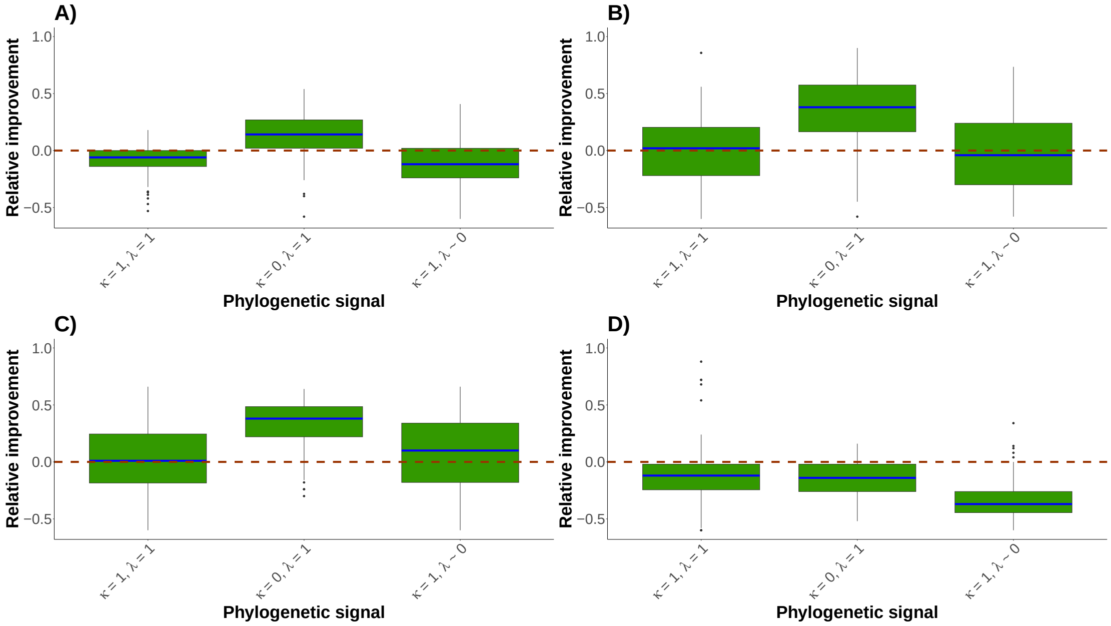

Figure 23: Boxplots representing the performance of the MICE+PI method according to different phylogenetic signals compared to PI method for the 4 missing mechanisms (MCAR (**A**), MAR (**B**), MNAR (**C**) and phyloNA (**D**)). Boxplots represent the difference between PI and MICE+PI accuracy. Therefore, the dashed line indicates no difference in performance between PI method and MICE+PI. The blue line in the boxplots is the median. Data simulated by a MK model and contained strongly correlated traits ($\rho = 0.8$). The missing rate is of 50%.

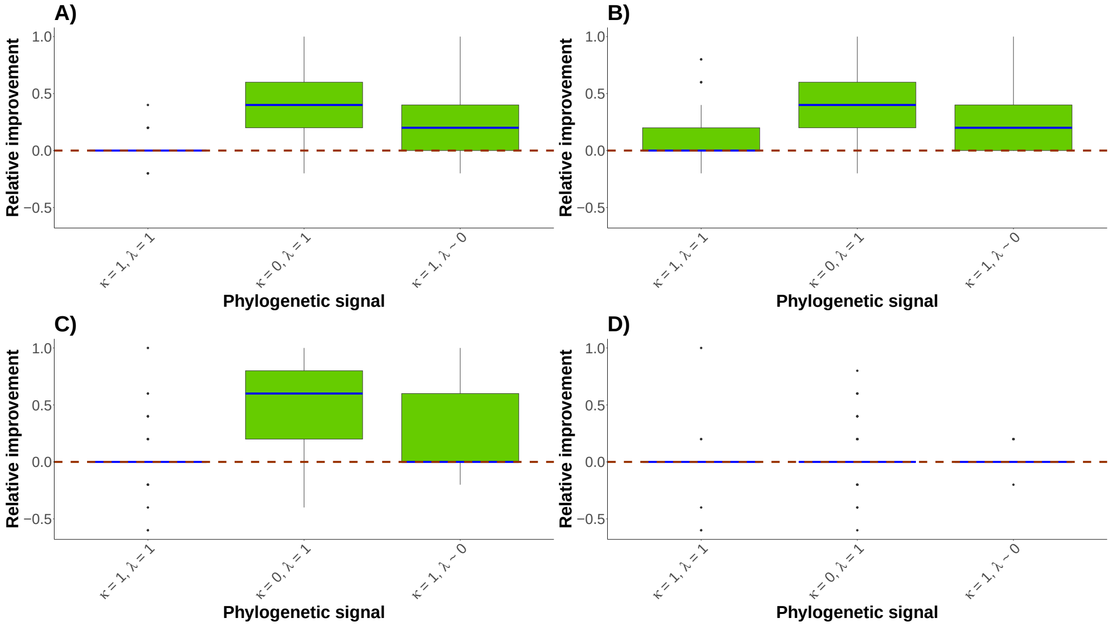

Figure 24: Boxplots representing the performance of the kNN+PI method according to different phylogenetic signals compared to PI method for the 4 missing mechanisms (MCAR (**A**), MAR (**B**), MNAR (**C**) and phyloNA (**D**)). Boxplots represent the difference between PI and kNN+PI accuracy. Therefore, the dashed line indicates no difference in performance between PI method and kNN+PI. The blue line in the boxplots is the median. Data simulated by a MK model and contained strongly correlated traits ($\rho = 0.8$). The missing rate is of 5%.

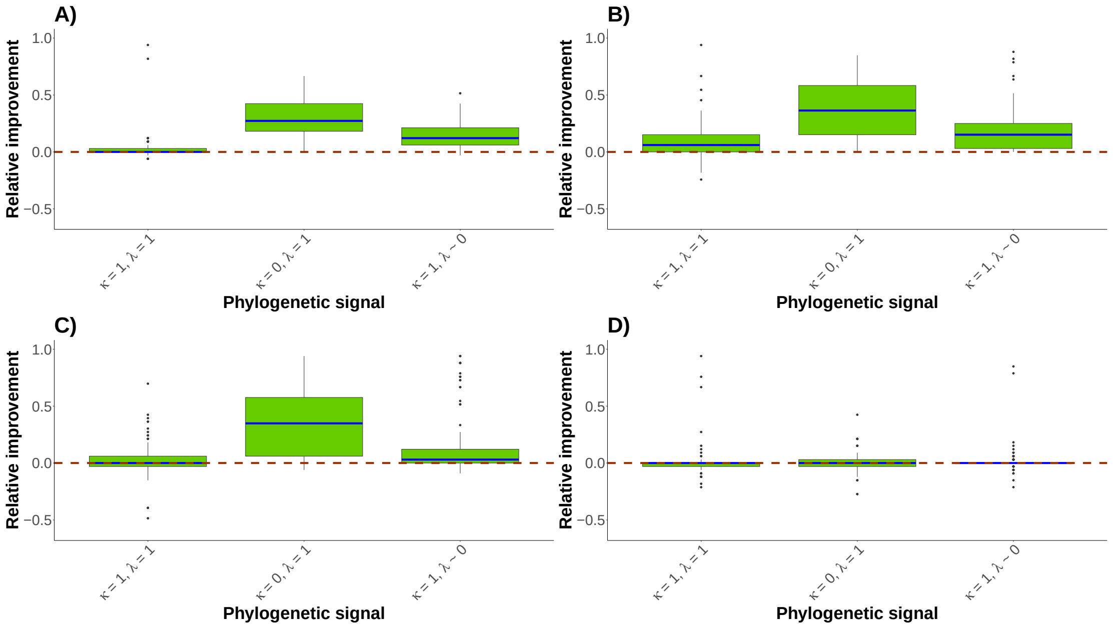

Figure 25: Boxplots representing the performance of the kNN+PI method according to different phylogenetic signals compared to PI method for the 4 missing mechanisms (MCAR (**A**), MAR (**B**), MNAR (**C**) and phyloNA (**D**)). Boxplots represent the difference between PI and kNN+PI accuracy. Therefore, the dashed line indicates no difference in performance between PI method and kNN+PI. The blue line in the boxplots is the median. Data simulated by a MK model and contained strongly correlated traits ($\rho = 0.8$). The missing rate is of 33%.

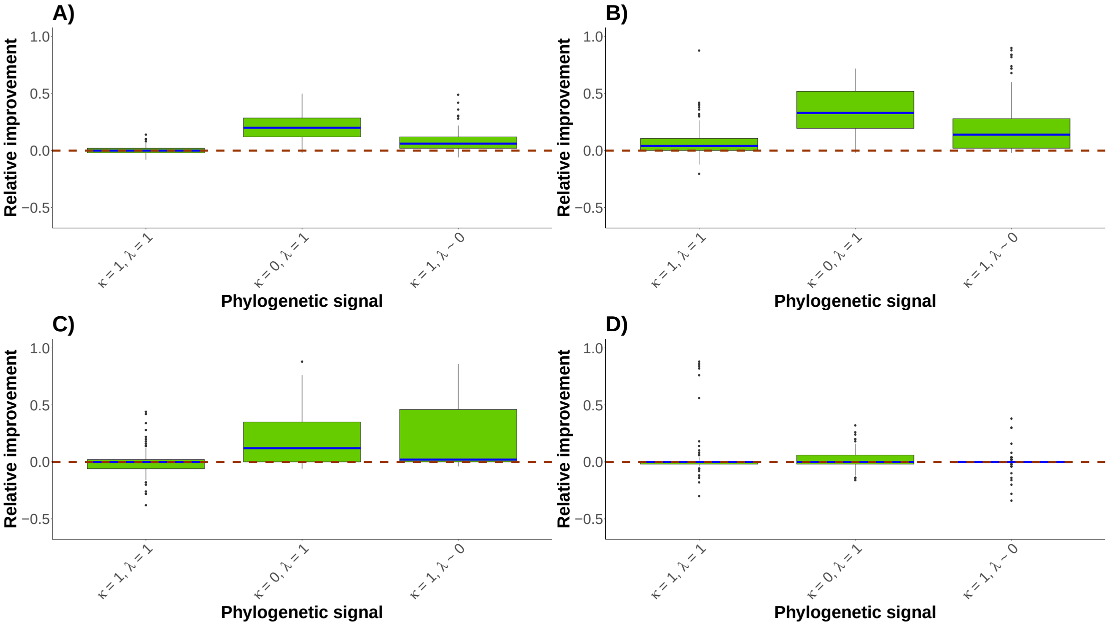

Figure 26: Boxplots representing the performance of the kNN+PI method according to different phylogenetic signals compared to PI method for the 4 missing mechanisms (MCAR (**A**), MAR (**B**), MNAR (**C**) and phyloNA (**D**)). Boxplots represent the difference between PI and kNN+PI accuracy. Therefore, the dashed line indicates no difference in performance between PI method and kNN+PI. The blue line in the boxplots is the median. Data simulated by a MK model and contained strongly correlated traits ($\rho = 0.8$). The missing rate is of 50%.

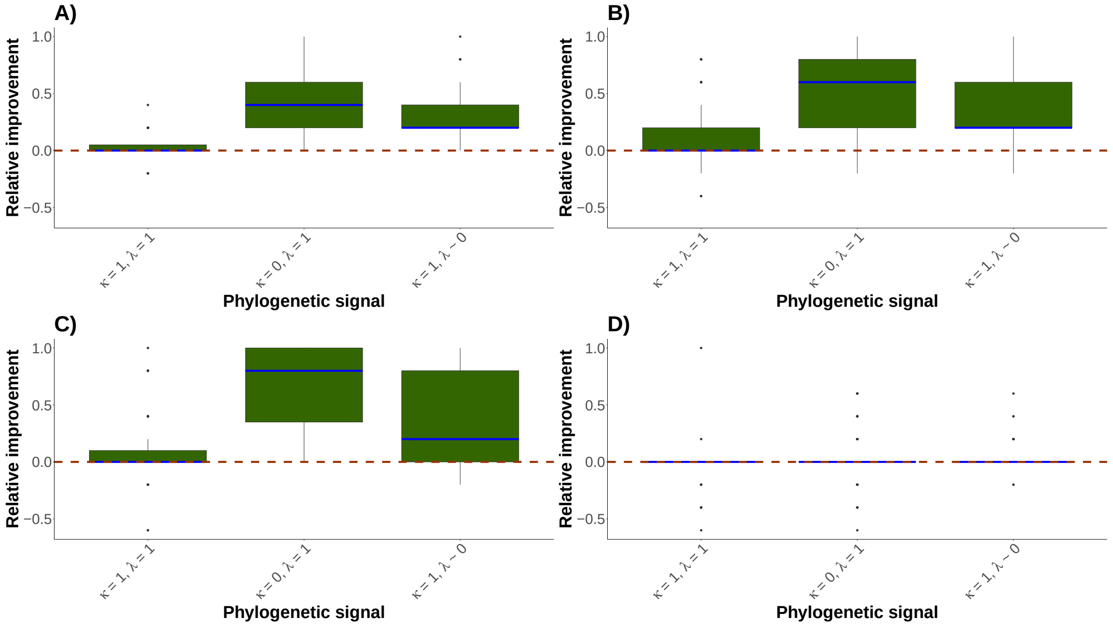

Figure 27: Boxplots representing the performance of the MissForest method according to different phylogenetic signals compared to PI method for the 4 missing mechanisms (MCAR (**A**), MAR (**B**), MNAR (**C**) and phyloNA (**D**)). Boxplots represent the difference between PI and MissForest accuracy. Therefore, the dashed line indicates no difference in performance between PI method and MissForest. The blue line in the boxplots is the median. Data simulated by a MK model and contained strongly correlated traits ($\rho = 0.8$). The missing rate is of 5%.

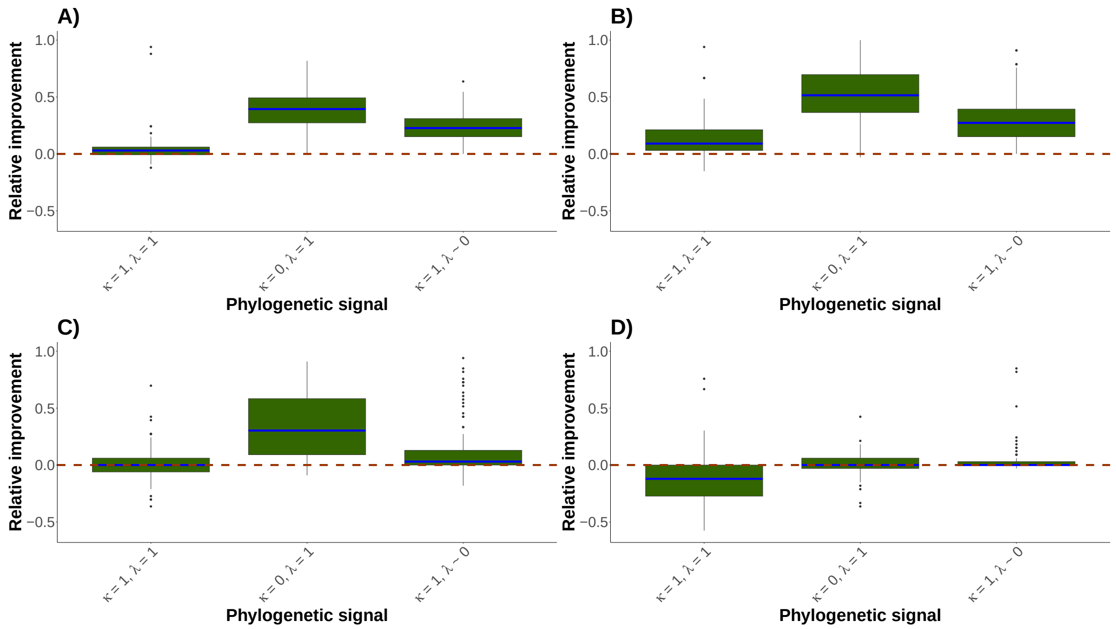

Figure 28: Boxplots representing the performance of the MissForest method according to different phylogenetic signals compared to PI method for the 4 missing mechanisms (MCAR (**A**), MAR (**B**), MNAR (**C**) and phyloNA (**D**)). Boxplots represent the difference between PI and MissForest accuracy. Therefore, the dashed line indicates no difference in performance between the PI method and MissForest. The blue line in the boxplots is the median. Data simulated by a MK model and contained strongly correlated traits ($\rho = 0.8$). The missing rate is of 33%.

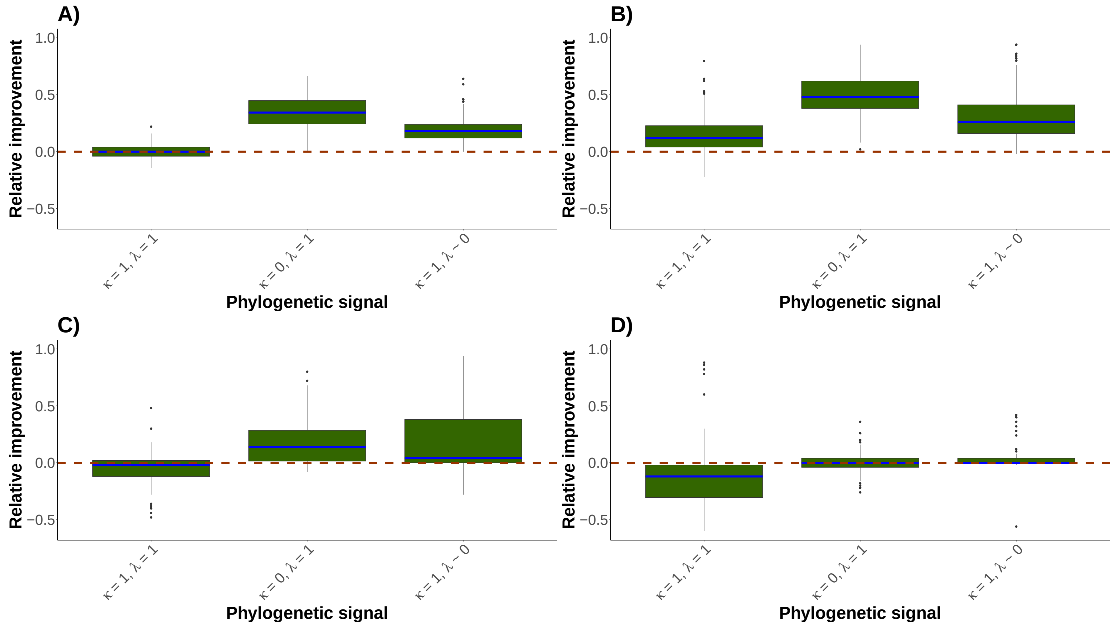

Figure 29: Boxplots representing the performance of the MissForest method according to different phylogenetic signals compared to PI method for the 4 missing mechanisms (MCAR (**A**), MAR (**B**), MNAR (**C**) and phyloNA (**D**)). Boxplots represent the difference between PI and MissForest accuracy. Therefore, the dashed line indicates no difference in performance between the PI method and MissForest. The blue line in the boxplots is the median. Data simulated by a MK model and contained strongly correlated traits ($\rho = 0.8$). The missing rate is of 50%.

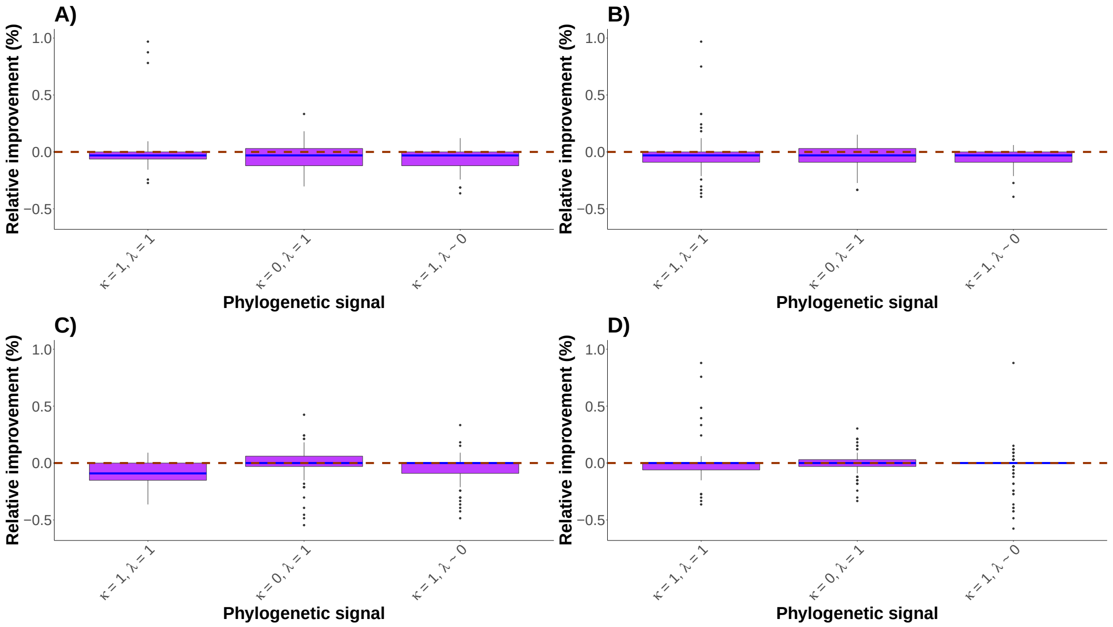

Figure 30: Boxplots representing the performance of the HV method according to different phylogenetic signals compared to PI method for the 4 missing mechanisms (MCAR (**A**), MAR (**B**), MNAR (**C**) and phyloNA (**D**)). Boxplots represent the difference between PI and HV accuracy. Therefore, the dashed line indicates no difference in performance between PI method and HV. The blue line in the boxplots is the median. Data simulated by a MK model and contained only independent traits. The missing rate is of 33%.

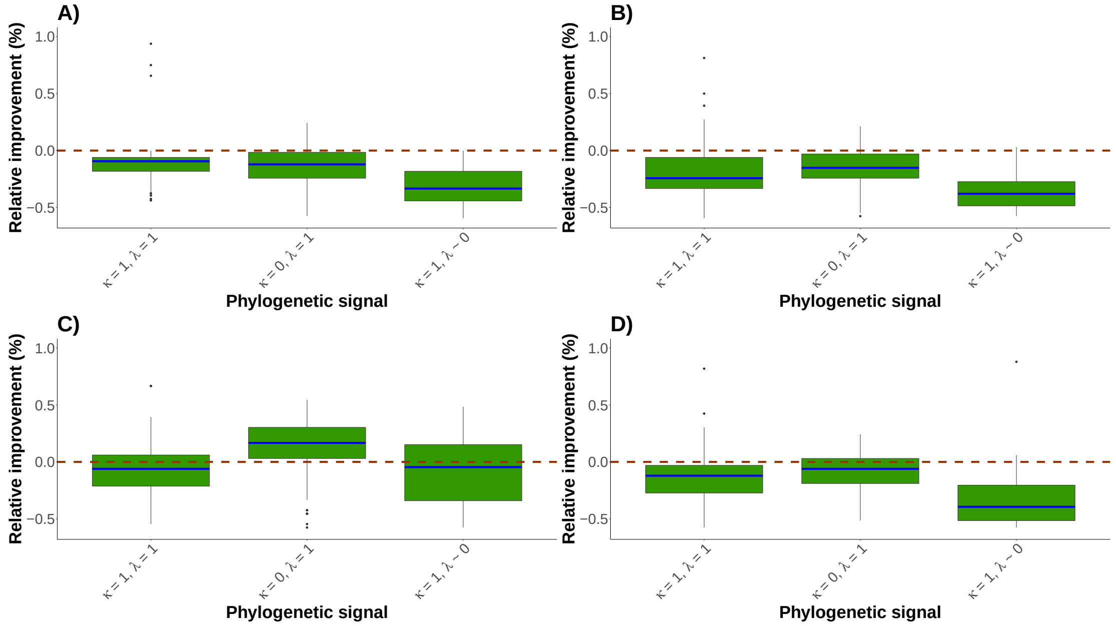

Figure 31: Boxplots representing the performance of the MICE+PI method according to different phylogenetic signals compared to PI method for the 4 missing mechanisms (MCAR (**A**), MAR (**B**), MNAR (**C**) and phyloNA (**D**)). Boxplots represent the difference between PI and MICE+PI accuracy. Therefore, the dashed line indicates no difference in performance between PI method and MICE+PI. The blue line in the boxplots is the median. Data simulated by a MK model and contained only independent traits. The missing rate is of 33%.

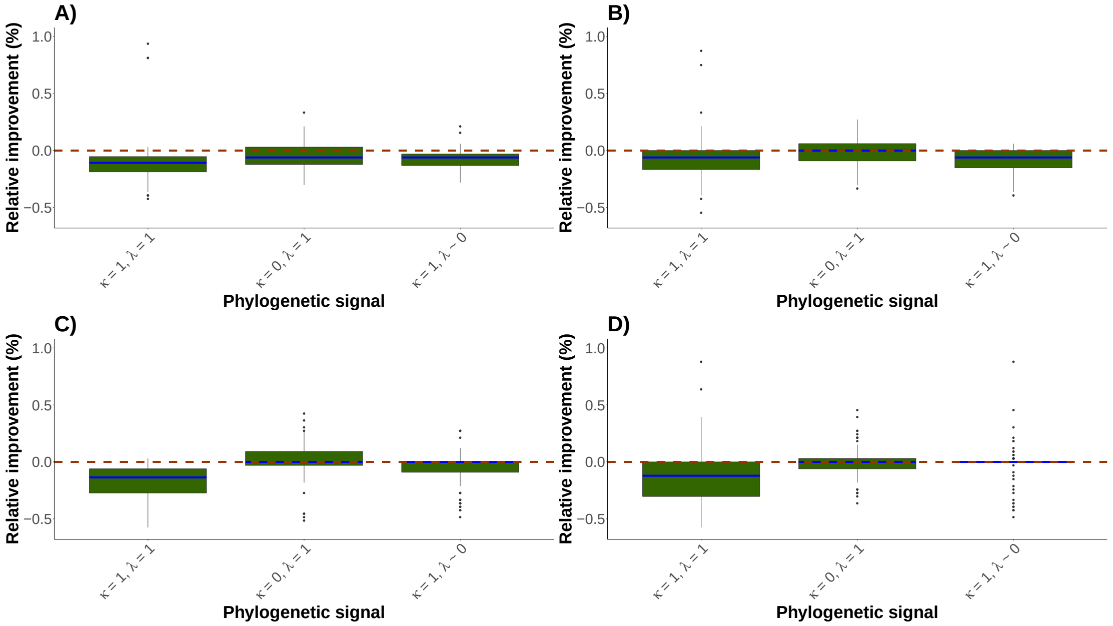

Figure 32: Boxplots representing the performance of the MissForest method according to different phylogenetic signals compared to PI method for the 4 missing mechanisms (MCAR (**A**), MAR (**B**), MNAR (**C**) and phyloNA (**D**)). Boxplots represent the difference between PI and MissForest accuracy. Therefore, the dashed line indicates no difference in performance between PI method and MissForest. The blue line in the boxplots is the median. Data simulated by a MK model and contained only independent traits. The missing rate is of 33%.

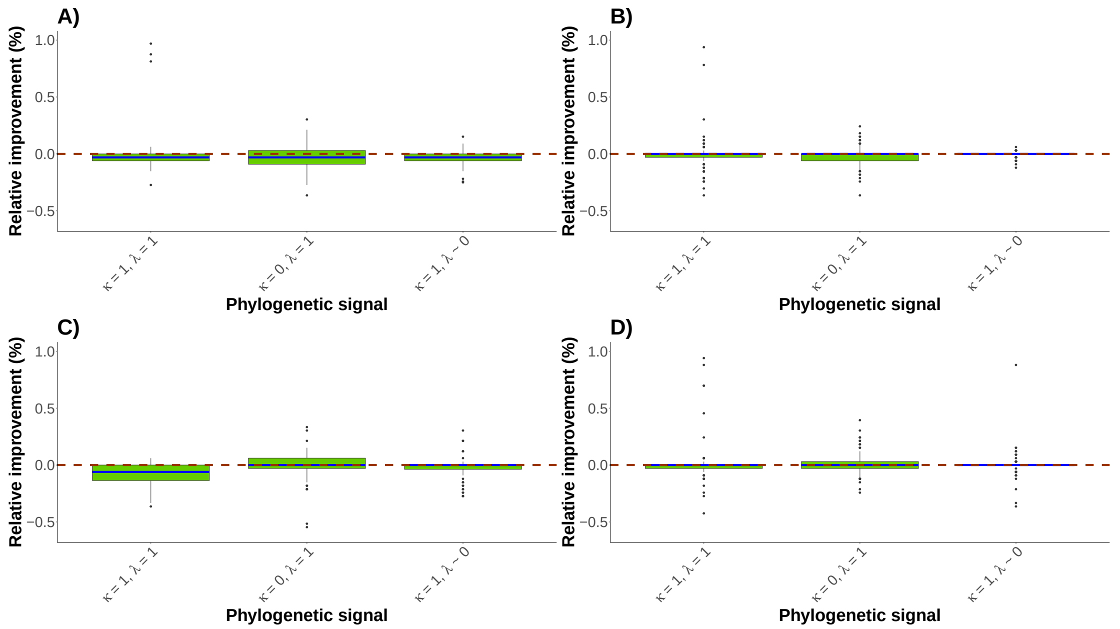

Figure 33: Boxplots representing the performance of the kNN+PI method according to different phylogenetic signals compared to PI method for the 4 missing mechanisms (MCAR (**A**), MAR (**B**), MNAR (**C**) and phyloNA (**D**)). Boxplots represent the difference between PI and kNN+PI accuracy. Therefore, the dashed line indicates no difference in performance between PI method and kNN+PI. The blue line in the boxplots is the median. Data simulated by a MK model and contained only independent traits. The missing rate is of 33%.

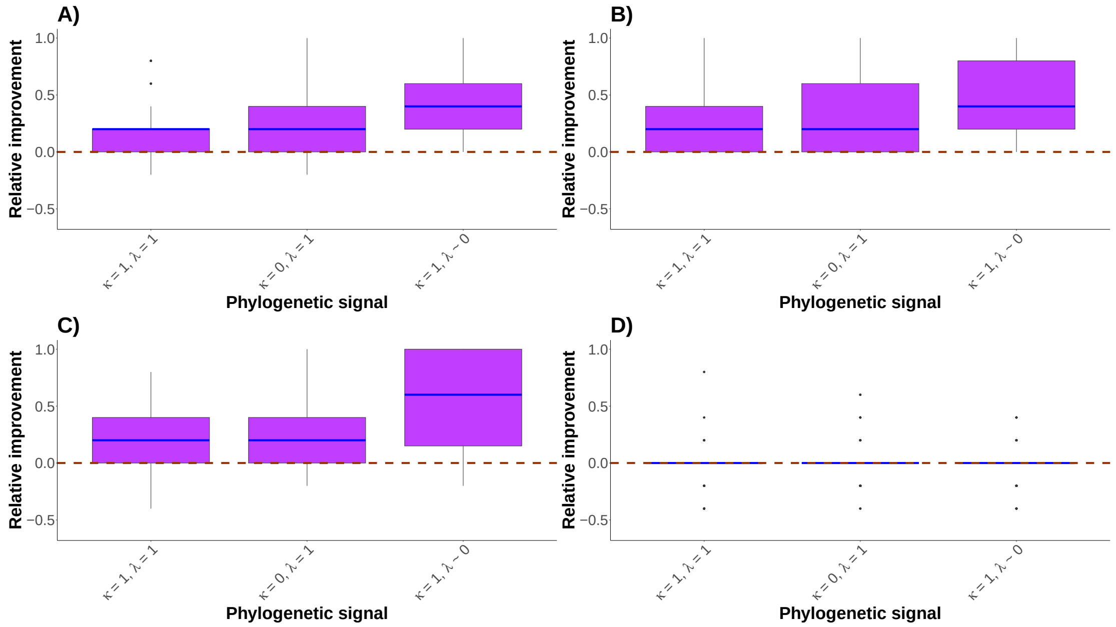

Figure 34: Boxplots representing the performance of the HV method according to different phylogenetic signals compared to PI method for the 4 missing mechanisms (MCAR (**A**), MAR (**B**), MNAR (**C**) and phyloNA (**D**)). Boxplots represent the difference between PI and HV accuracy. Therefore, the dashed line indicates no difference in performance between PI method and HV. The blue line in the boxplots is the median. Data simulated by a threshold model and contained strongly correlated traits ($\rho = 0.8$). The missing rate is of 5%.

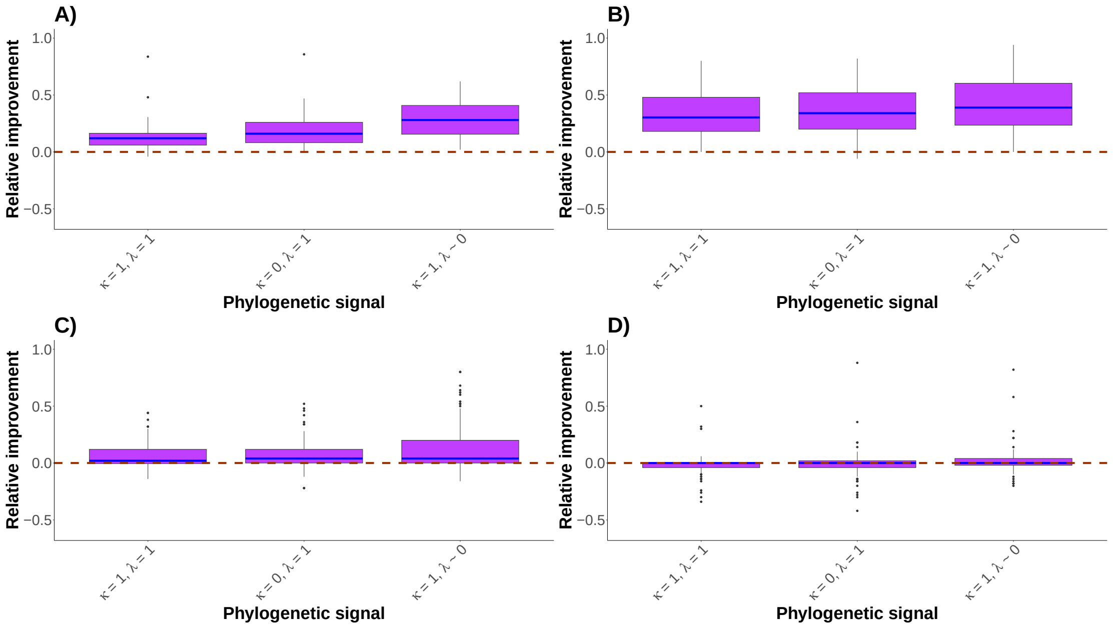

Figure 35: Boxplots representing the performance of the HV method according to different phylogenetic signals compared to PI method for the 4 missing mechanisms (MCAR (**A**), MAR (**B**), MNAR (**C**) and phyloNA (**D**)). Boxplots represent the difference between PI and HV accuracy. Therefore, the dashed line indicates no difference in performance between PI method and HV. The blue line in the boxplots is the median. Data simulated by a threshold model and contained strongly correlated traits ($\rho = 0.8$). The missing rate is of 50%.

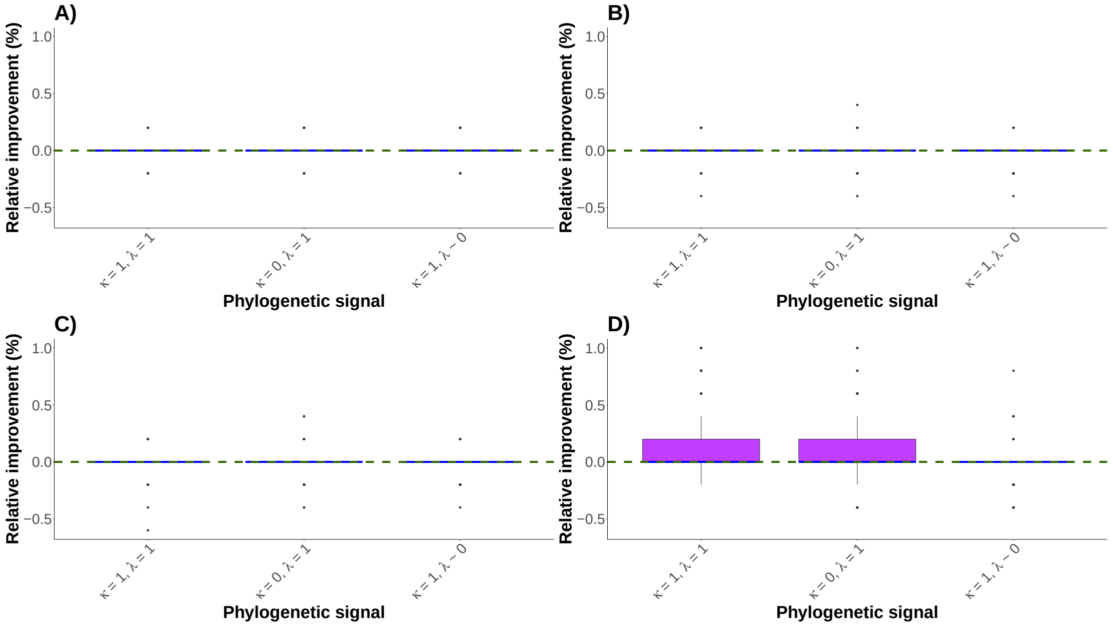

Figure 36: Boxplots representing the performance of the HV method according to different phylogenetic signals compared to MissForest method for the 4 missing mechanisms (MCAR (**A**), MAR (**B**), MNAR (**C**) and phyloNA (**D**)). Boxplots represent the difference between MissForest and HV accuracy. Therefore, the dashed line indicates no difference in performance between the MissForest method and HV. The blue line in the boxplots is the median. Data simulated by a threshold model and contained strongly correlated traits ($\rho = 0.8$). The missing rate is of 5%.

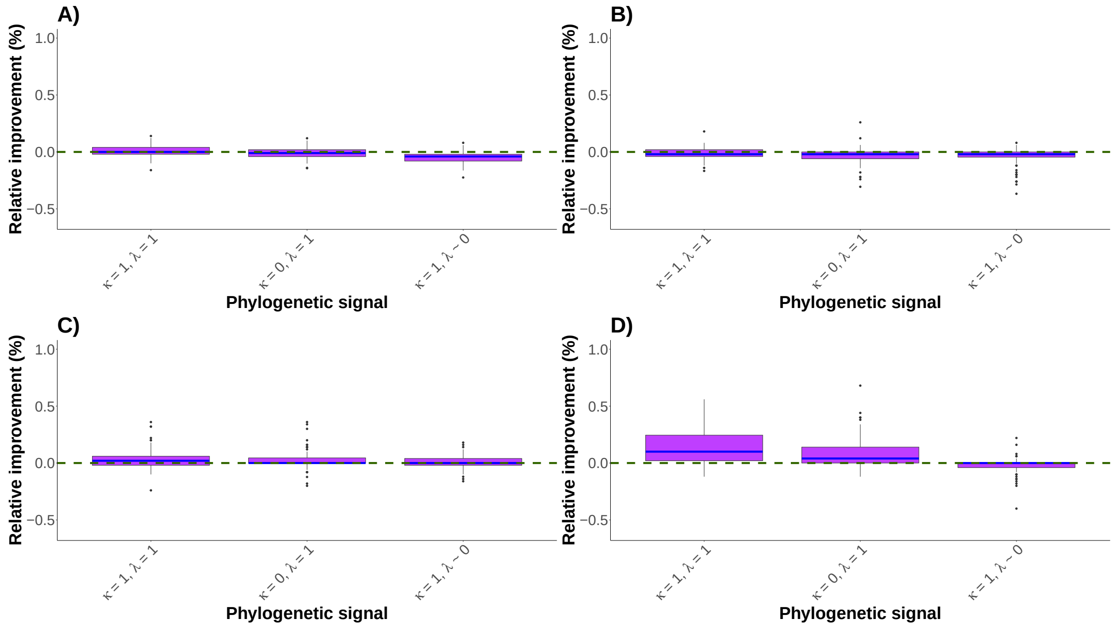

Figure 37: Boxplots representing the performance of the HV method according to different phylogenetic signals compared to MissForest method for the 4 missing mechanisms (MCAR (**A**), MAR (**B**), MNAR (**C**) and phyloNA (**D**)). Boxplots represent the difference between MissForest and HV accuracy. Therefore, the dashed line indicates no difference in performance between the MissForest method and HV. The blue line in the boxplots is the median. Data simulated by a threshold model and contained strongly correlated traits ($\rho = 0.8$). The missing rate is of 50%.

### 5 Outputs of the empirical data

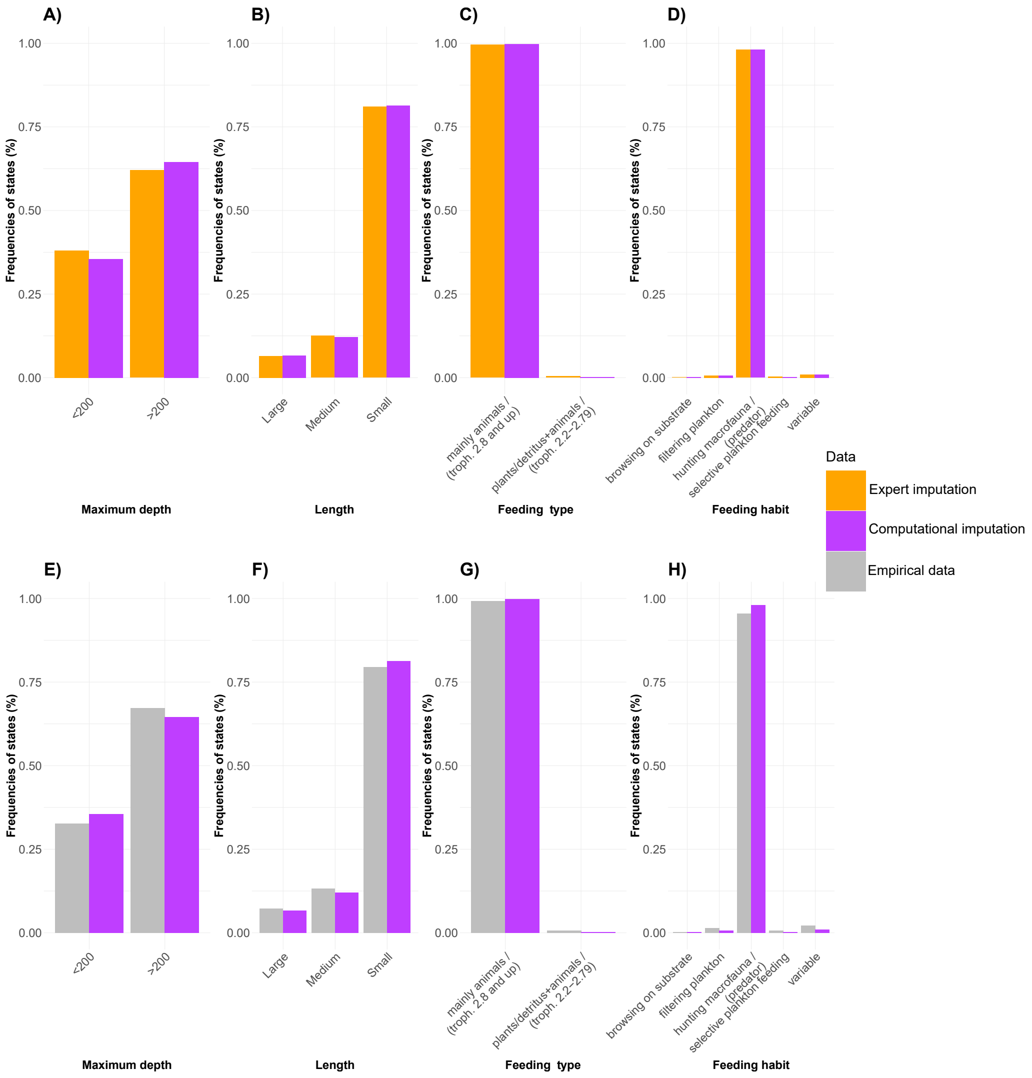

Figure 38: State frequencies in empirical data compared to state frequencies after expert imputation by an expert and after HV method imputation. The figures **A)** - **D)** compare the state frequency in the expert imputed data with the imputed data by HV while the figures **E)** - **H)** compare the state frequency in the original data and the imputed data by HV while the figures. **A)** and **E)** States frequencies of the Maximum depth trait. **B)** and **F)** States frequencies of the Length trait. **C)** and **G)** States frequencies of the Feeding type trait. **D)** and **H)** States frequencies of the Feeding habitat trait.

Table 28: Description of the four traits of interest: Maximum depth, Length, Feeding type and Feeding habit with 3 contingency columns describing the counts of each state before and after the expert imputation and the artificial addition of 1% MCAR values (random sampling). The last 2 columns highlight the amount of NA in each state.

| | | | Counts / Amount of NAs | | |
|---|---|---|---|---|---|
| Traits | Amount of NAs in traits | States | Empirical data | Expert data | Random sampling |
| Maximum depth | 0.14 | <200 | 285 / 0.26 | 383 | 282 / 0.01 |
| | | >200 | 587 / 0.06 | 624 | 581 / 0.01 |
| Length | 0.10 | Small | 727 / 0.12 | 822 | 720 / 0.01 |
| | | Medium | 121 / 0.05 | 127 | 120 / 0.01 |
| | | Large | 66 / 0.00 | 66 | 65 / 0.02 |
| Feeding type | 0.55 | mainly animals (troph. 2.8 and up) | 451 / 0.33 | 672 | 446 / 0.01 |
| | | plants/detritus+animals (troph. 2.2-2.79) | 3 / 0.00 | 3 | 2 / 0.33 |
| Feeding habit | 0.58 | filtering plankton | 6 / 0.00 | 6 | 5 / 0.17 |
| | | hunting macrofauna (predator) | 407 / 0.58 | 975 | 403 / 0.01 |
| | | selective plankton feeding | 3 / 0.00 | 3 | 2 / 0.33 |
| | | variable | 9 / 0.00 | 9 | 8 / 0.11 |

Table 29: Accuracies obtained from the expert-based approach and random sampling approach using the methods MICE+PI, missForest, kNN+PI and HV. The traits evaluated were Maximum depth, Length, Feeding type and Feeding habit.

| Methods | Traits | Expert-based | Random sampling |
|---|---|---|---|
| MICE+PI | Maximum depth | 0.80 | 0.61(0.20) |
| | Length | 0.85 | 0.80(0.08) |
| | Feeding type | 1.00 | 0.82(0.05) |
| | Feeding habit | 0.93 | 0.55(0.06) |
| MissForest | Maximum depth | 0.81 | 0.87(0.13) |
| | Length | 0.96 | 0.88(0.11) |
| | Feeding type | 1.00 | 0.84(0.00) |
| | Feeding habit | 0.98 | 0.61(0.07) |
| kNN+PI | Maximum depth | 0.80 | 0.64(0.22) |
| | Length | 0.94 | 0.81(0.07) |
| | Feeding type | 1.00 | 0.83(0.00) |
| | Feeding habit | 0.99 | 0.61(0.07) |
| HV | Maximum depth | 0.81 | 0.80(0.15) |
| | Length | 0.94 | 0.89(0.07) |
| | Feeding type | 1.00 | 0.83(0.00) |
| | Feeding habit | 0.99 | 0.57(0.00) |

Table 30: Table comparing the state frequencies of 3 traits of the empirical data (Maximum depth, Length, Feeding type and Feeding habit) before and after the imputation of the missing values.

| States | Original data | Computationally data | Manually imputed data |
|---|---|---|---|
| $< 200$ | 0.327 | 0.355 | 0.380 |
| $> 200$ | 0.673 | 0.645 | 0.620 |

(a) Maximum depth

| States | Original data | Computationally data | Manually imputed data |
|---|---|---|---|
| Small | 0.795 | 0.813 | 0.810 |
| Medium | 0.132 | 0.121 | 0.125 |
| Large | 0.072 | 0.066 | 0.065 |

(b) Length

| States | Original data | Computationally data | Manually imputed data |
|---|---|---|---|
| mainly animals (troph. 2.8 and up) | 0.993 | 0.998 | 0.996 |
| plants/ detritus+animals (troph. 2.2-2.79) | 0.007 | 0.002 | 0.004 |

(c) Feeding type

| States | Original data | Computationally imputed data | Manually imputed data |
|---|---|---|---|
| browsing on substrate | 0.002 | 0.002 | 0.001 |
| filtering plankton | 0.014 | 0.006 | 0.006 |
| hunting macrofauna (predator) | 0.955 | 0.981 | 0.981 |
| selective plankton feeding | 0.007 | 0.002 | 0.003 |
| variable | 0.021 | 0.009 | 0.009 |

(d) Feeding habit

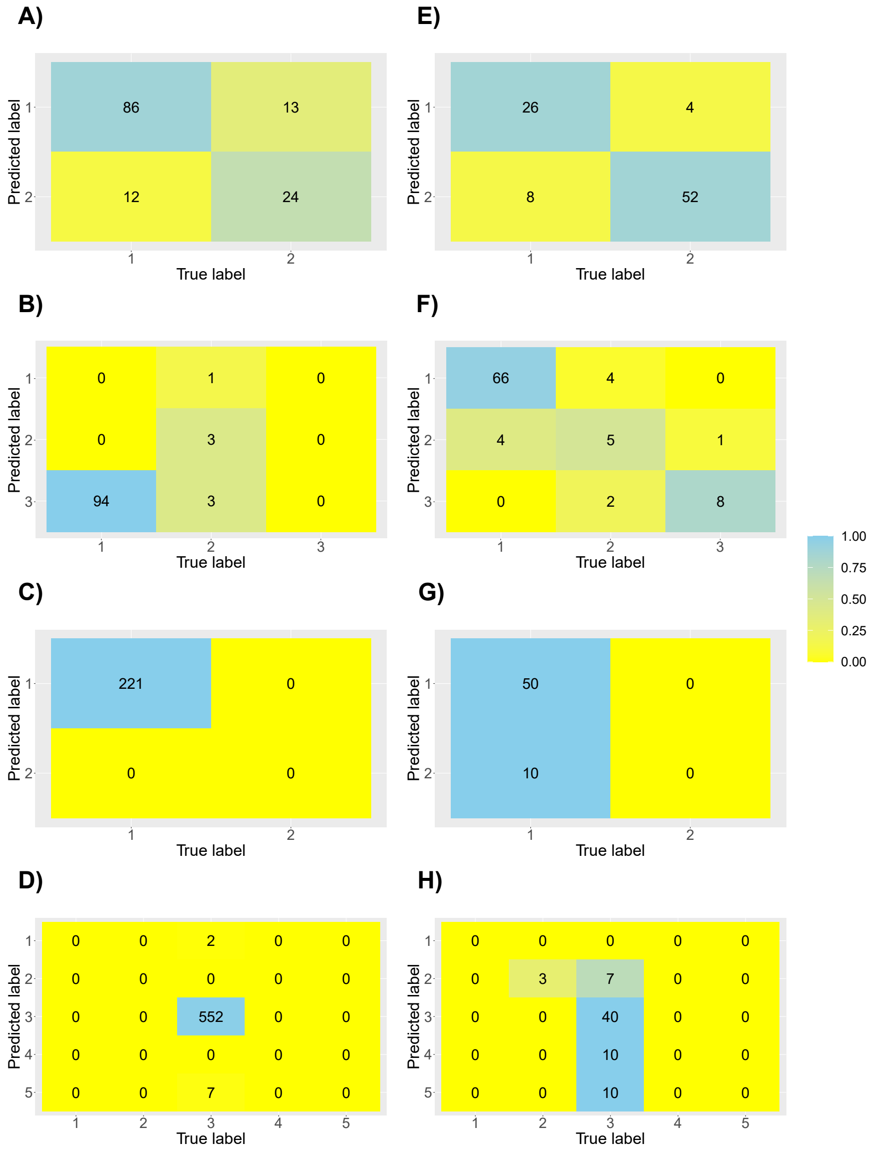

Figure 39: Confusion matrix obtained after imputation of missing values in the empirical dataset with MissForest. The true label for **A)** to **D)** is the expert filled dataset while for **E)** to **D)** the true label is the empirical data in which we applied the random sampling approach. In this case, the matrices correspond to the sum of the 10 replicated confusion matrices. **A)** and **E)** correspond to the quality of imputation of the "Maximum depth" trait (1: "$< 200$", 2: "$>= 200$"). **B)** and **F)** are for the "Length" trait (1: "Small", 2: "Medium", 3: "Large"). **C)** and **G)** are for the "Feeding type" trait (1: "mainly animals", 2: "plants/detritus+animals"). **D)** and **H)** are for the "Feeding habit" trait (1: "browsing on substrate", 2: "filtering plankton", 3: "hunting macrofauna (predator)", 4: "selective plankton feeding" , 5: "variable"). The color scale correspond to the proportion of value that are correctly imputed (True positive rate).

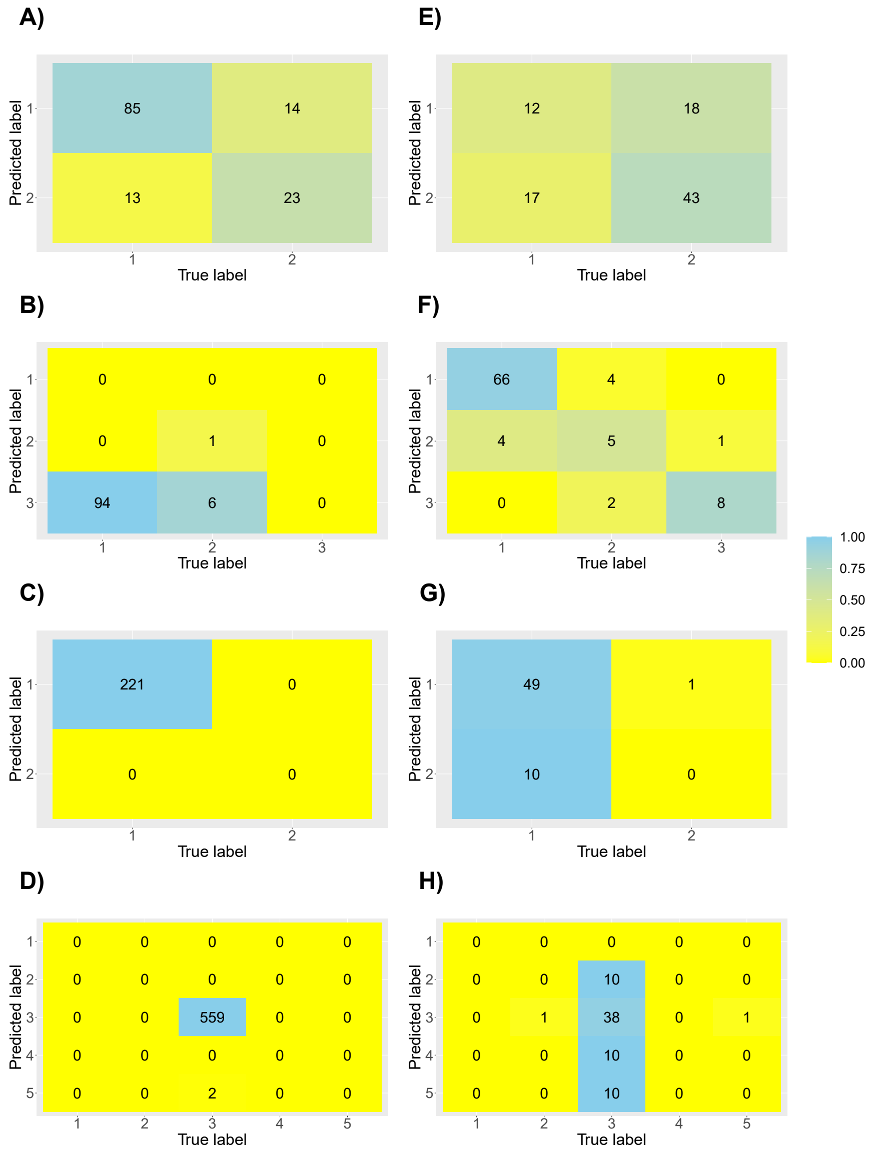

Figure 40: Confusion matrix obtained after imputation of missing values in the empirical dataset with kNN+PI. The true label for **A)** to **D)** is the expert filled dataset while for **E)** to **D)** the true label is the empirical data in which we applied the random sampling approach. In this case, the matrices correspond to the sum of the 10 replicated confusion matrices. **A)** and **E)** correspond to the quality of imputation of the "Maximum depth" trait (1: "$<200$", 2: "$>=200$"). **B)** and **F)** are for the "Length" trait (1: "Small", 2: "Medium", 3: "Large"). **C)** and **G)** are for the "Feeding type" trait (1: "mainly animals", 2: "plants/detritus+animals"). **D)** and **H)** are for the "Feeding habit" trait (1: "browsing on substrate", 2: "filtering plankton", 3: "hunting macrofauna (predator)", 4: "selective plankton feeding" , 5: "variable"). The color scale correspond to the proportion of value that are correctly imputed (True positive rate).

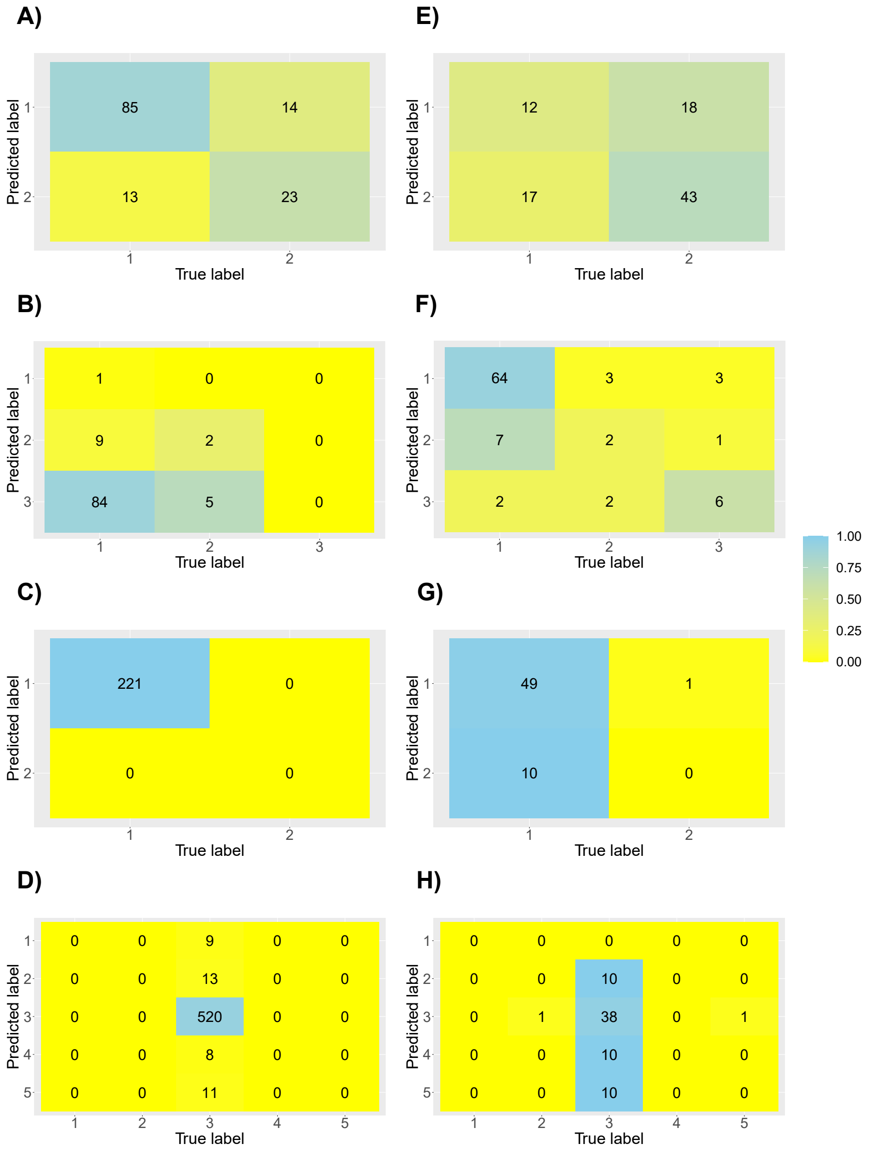

Figure 41: Confusion matrix obtained after imputation of missing values in the empirical dataset with MICE+PI. The true label for **A)** to **D)** is the expert filled dataset while for **E)** to **D)** the true label is the empirical data in which we applied the random sampling approach. In this case, the matrices correspond to the sum of the 10 replicated confusion matrices. **A)** and **E)** correspond to the quality of imputation of the "Maximum depth" trait (1: "$< 200$", 2: "$>= 200$"). **B)** and **F)** are for the "Length" trait (1: "Small", 2: "Medium", 3: "Large"). **C)** and **G)** are for the "Feeding type" trait (1: "mainly animals", 2: "plants/detritus+animals"). **D)** and **H)** are for the "Feeding habit" trait (1: "browsing on substrate", 2: "filtering plankton", 3: "hunting macrofauna (predator)", 4: "selective plankton feeding" , 5: "variable"). The color scale correspond to the proportion of value that are correctly imputed (True positive rate).

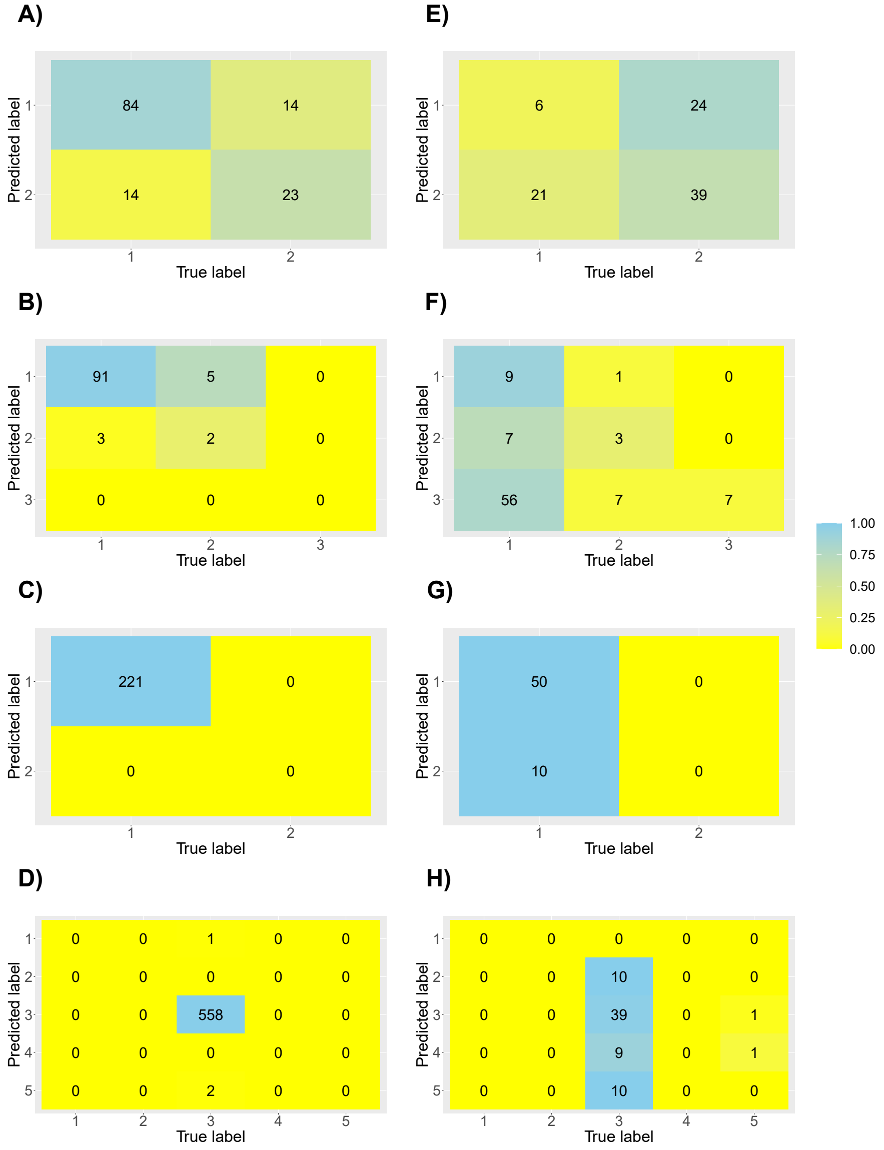

Figure 42: Confusion matrix obtained after imputation of missing values in the empirical dataset with PI. The true label for **A)** to **D)** is the expert filled dataset while for **E)** to **D)** the true label is the empirical data in which we applied the random sampling approach. In this case, the matrices correspond to the sum of the 10 replicated confusion matrices. **A)** and **E)** correspond to the quality of imputation of the "Maximum depth" trait (1: "$< 200$", 2: "$>= 200$"). **B)** and **F)** are for the "Length" trait (1: "Small", 2: "Medium", 3: "Large"). **C)** and **G)** are for the "Feeding type" trait (1: "mainly animals", 2: "plants/detritus+animals"). **D)** and **H)** are for the "Feeding habit" trait (1: "browsing on substrate", 2: "filtering plankton", 3: "hunting macrofauna (predator)", 4: "selective plankton feeding" , 5: "variable"). The color scale correspond to the proportion of value that are correctly imputed (True positive rate).

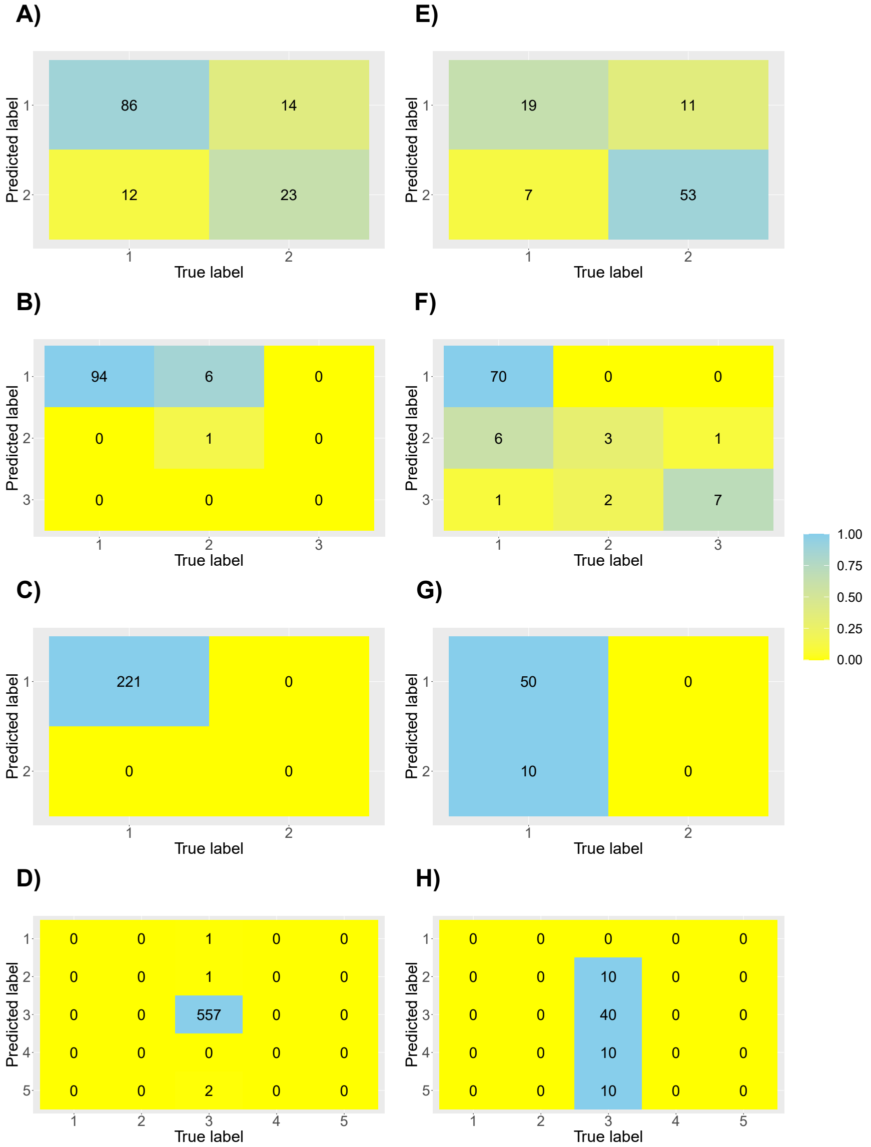

Figure 43: Confusion matrix obtained after imputation of missing values in the empirical dataset with HV method. The true label for **A)** to **D)** is the expert filled dataset while for **E)** to **D)** the true label is the empirical data in which we applied the random sampling approach. In this case, the matrices correspond to the sum of the 10 replicated confusion matrices. **A)** and **E)** correspond to the quality of imputation of the "Maximum depth" trait (1: "$< 200$", 2: "$>= 200$"). **B)** and **F)** are for the "Length" trait (1: "Small", 2: "Medium", 3: "Large"). **C)** and **G)** are for the "Feeding type" trait (1: "mainly animals", 2: "plants/detritus+animals"). **D)** and **H)** are for the "Feeding habit" trait (1: "browsing on substrate", 2: "filtering plankton", 3: "hunting macrofauna (predator)", 4: "selective plankton feeding" , 5: "variable"). The color scale correspond to the proportion of value that are correctly imputed (True positive rate).
